# Slow spatial migration can help eradicate cooperative antimicrobial resistance in time-varying environments

**DOI:** 10.1101/2024.12.30.630406

**Authors:** Lluís Hernández-Navarro, Kenneth Distefano, Uwe C. Täuber, Mauro Mobilia

**Affiliations:** Department of Applied Mathematics, School of Mathematics, University of Leeds, Leeds LS2 9JT, U.K.; Department of Physics & Center for Soft Matter and Biological Physics, MC 0435, Robeson Hall, 850 West Campus Drive, Virginia Tech, Blacksburg, VA 24061, USA; Faculty of Health Sciences, Virginia Tech, Blacksburg, VA 24061, USA

## Abstract

Antimicrobial resistance (AMR) is a global threat and combating its spread is of paramount importance. AMR often results from a cooperative behaviour with shared drug protection. Microbial communities generally evolve in volatile, spatially structured settings. Migration, space, fluctuations, and environmental variability all have a significant impact on the development and proliferation of AMR. While drug resistance is enhanced by migration in static conditions, this changes in time-fluctuating spatially structured environments. Here, we consider a two-dimensional metapopulation consisting of demes in which drug-resistant and sensitive cells evolve in a time-changing environment. This contains a toxin against which protection can be shared (cooperative AMR). Cells migrate between demes and connect them. When the environment and the deme composition vary on the same timescale, strong population bottlenecks cause fluctuation-driven extinction events, countered by migration. We investigate the influence of migration and environmental variability on the AMR eco-evolutionary dynamics by asking at what migration rate fluctuations can help clear resistance and what are the near-optimal environmental conditions ensuring the quasi-certain eradication of resistance in the shortest possible time. By combining analytical and computational tools, we answer these questions by determining when the resistant strain goes extinct across the entire metapopulation. While dispersal generally promotes strain coexistence, here we show that slow-but-nonzero migration can speed up and enhance resistance clearance, and determine the near-optimal conditions for this phenomenon. We discuss the impact of our findings on laboratory-controlled experiments and outline their generalisation to lattices of any spatial dimension.

**Author summary:** As the number of microbes resisting antimicrobial drugs grows alarmingly, it is of paramount importance to tackle this major societal issue. Resistant microbes often inactivate antibiotic drugs in the environment around them, and hence offer protection to drug-sensitive bacteria in a form of cooperative behaviour. Moreover, microbes typically are distributed in space and live in time-changing environments, where they are subject to random fluctuations. Environmental variability, fluctuations, and spatial dispersal all have a strong influence on the drug resistance of microbial organisms. Here we investigate the temporal evolution of antimicrobial resistance in time-varying spatial environments by combining computational and mathematical means. We study the dynamics of drug-resistant and sensitive cells in the presence of an antimicrobial drug, when microbes are spatially distributed across a (two-dimensional) grid of well-mixed sub-populations (demes). Cells migrate between neighbouring demes, connecting these sub-populations, and are subject to sudden changes of environment, homogeneously across all demes. We show that when the environment and the deme composition vary on the same timescale, the joint effect of slow migration and fluctuations can help eradicate drug resistance by speeding up and enhancing the extinction probability of resistant bacteria. We also discuss how our findings can be probed in laboratory experiments, and outline their generalisation to lattices of any dimension.

## Introduction

Microbial communities generally live in volatile, time-varying environments embedded in complex spatial structures connected through cellular migration, e.g., in soil [1], seabed [2], on wet surfaces [3], in plants [4], animals [5], and humans [6, 7]. How the environment helps shape microbial populations and species diversity [8, 9, 10, 11] is a subject of intense research [11, 12, 13, 14, 15, 16, 17, 18, 19, 20, 21, 22]. Moreover, environmental variability and microbiome-environment interactions greatly influence the temporal evolution of microbial communities, with a growing interest in their eco-evolutionary dynamics [6, 10, 22, 23, 24, 25, 26, 27, 28]. Despite significant recent progress [13, 14, 21, 22], a general understanding of the combined influence of spatial structure, migration and environmental variability on the evolution of microbial populations remains an open question that is notably relevant to the spread of antimicrobial resistance (AMR) [29, 30, 31, 32, 33, 34, 35, 36, 37, 38, 39]. Understanding the spread of AMR is of paramount societal importance, and is influenced by spatial structure, environmental changes and fluctuations. The latter are often associated with population bottlenecks [10], when the community size is drastically reduced, e.g., due to the effects of drugs [40, 41] or other causes [42, 43]. The size and composition of microbial populations are often interdependent, leading to coupled environmental and demographic fluctuations [44, 23, 45, 46, 47, 48, 49, 26, 27, 28, 50, 51, 52, 53, 54, 55]. These are particularly relevant when antibiotics cause bottlenecks following which surviving cells may replicate and AMR can spread [41]. Recent studies have investigated the impact of space on the emergence and spread of non-cooperative AMR mutants [11, 15], even in the presence of environmental bottlenecks [20, 22]. However, AMR often results from cooperative behaviour with resistant microbes inactivating toxins and sharing their protection with drug-sensitive bacteria [21, 43, 56]. Recent research on cooperative AMR has mostly focused on microbiome-environment interactions [14, 21], without considering external environmental changes.

Moreover, recent studies on *rescue dynamics*, which investigate the recovery of near-extinction populations, have also focused on the role of spatial recolonisation – that is, the reoccupation of a previously emptied local area through cell migration (demographic rescue). These processes were first investigated in single-strain systems subject to constant environments [57], and then in the presence of different forms of environmental variability, e.g., [58, 59, 60, 61, 62, 63]. These works showed that environmental stochasticity typically limits the rescue effect and is detrimental for population survival (but see Ref. [64], where population rescue can occur more readily in the face of harsher environmental shifts). Additionally, recent computational studies [65, 66, 67, 68] and experimental works [69, 70, 71, 72, 73] have reported that the survival of a population is enhanced when the rate of cell migration is intermediate, i.e. when individuals are neither organised in entirely isolated patches nor in fully connected (well-mixed) subpopulations. This stems from recolonisation events following local extinction, and has notably been reported for the case of cooperative AMR [14].

Here, inspired by the cooperative nature of *β-*lactamase-mediated AMR [43, 56, 74], we study how the migration of cells shapes the temporal evolution of cooperative resistance (modelled using a public good threshold, see below) in a spatially structured microbial population subject to environmental variability causing bottlenecks and fluctuations. To this end, we investigate the *in silico* temporal evolution of cooperative AMR in a two-dimensional (2D) metapopulation consisting of cells that are either sensitive or resistant to a drug. (Note that in this work we use the term *evolution* to refer to the competition dynamics between drug-resistant and sensitive strains, hence ignoring mutations that often occur on a longer timescale than the phenomenon studied here; see Model & Methods and Discussion). We consider a spatially explicit model consisting of a grid of demes whose well-mixed sub-populations are connected by cell migration, as commonly used to model microbial communities living on surfaces, in theory and experiments [75, 76, 31, 77]. (The case of a one-dimensional lattice of demes is discussed in S1 Appendix Sec. 5.5, see below). The metapopulation is subject to a constant antimicrobial input and a time-fluctuating environment, that is homogeneous across all demes. We model environmental variability by letting the carrying capacity of each deme change simultaneously in time to represent harsh and mild environmental conditions (Fig 1A), thereby implementing bottlenecks [51, 52, 53, 78, 55, 54, 26, 28, 27, 79]. These are critical for microbial dynamics and can be engineered in laboratory-controlled experiments [80, 55, 81, 41, 82, 83, 84, 50, 85, 86, 87, 88, 89, 90, 91, 92]. Changes in the carrying capacity can potentially encode different exogenous sources of environmental variability. Here, inspired by recent chemostat and microfluidic experiments [25, 93, 94], we interpret environmental variability as representing a time-varying influx of nutrients (or sequential changes with a secondary antibiotic [95]).

**Figure 1.**
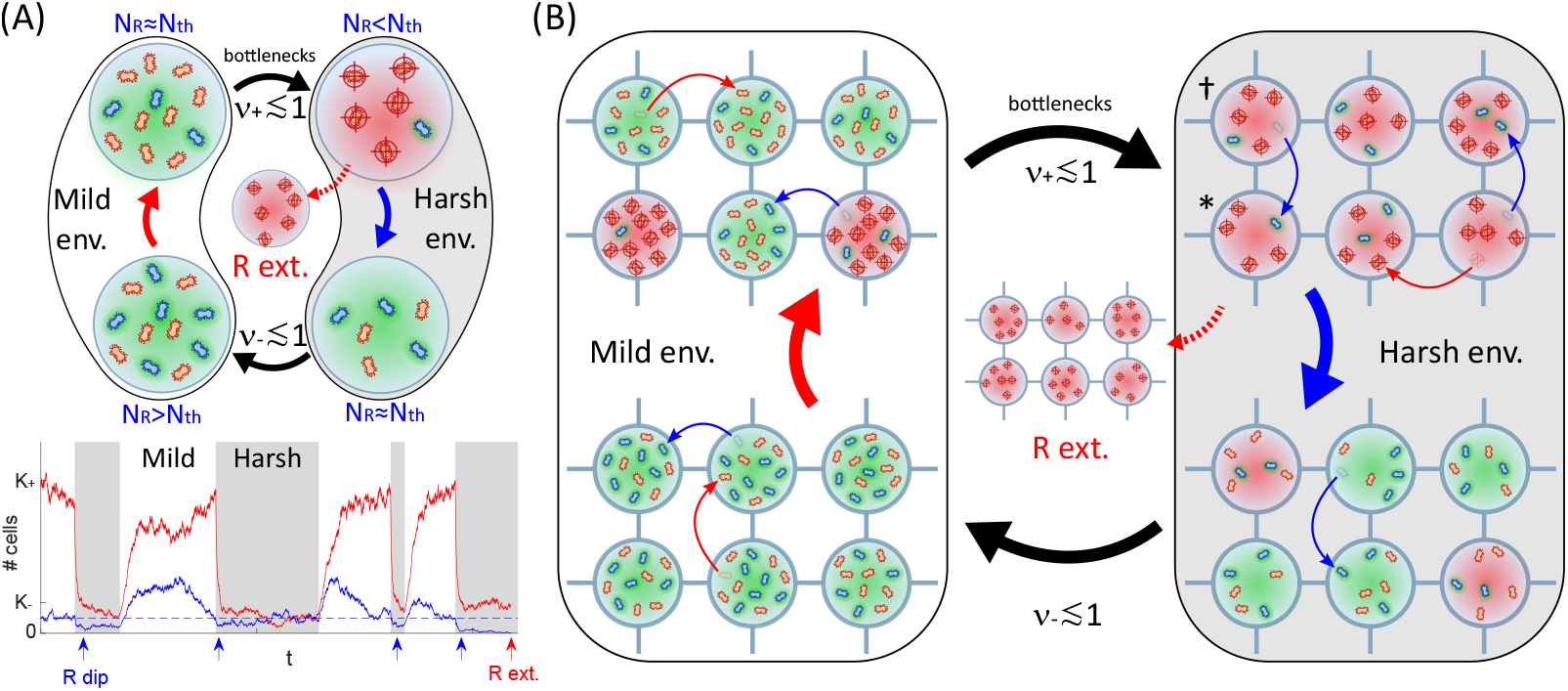
Microbial community model. Panel A: Eco-evolutionary dynamics in an isolated deme (*m* = 0) subject to constant antimicrobial input rate and intermediate environmental switching (Model & Methods). Top: Illustrative temporal evolution when the environment switches between mild (*K*_+_ = 12) and harsh (*K*_−_ = 6) environments (env.) at rates *ν*_*±*_ ≲ 1, with cooperation threshold *N*_th_ = 3 (Model & Methods). Resistant microbes (blue, *R*) produce a resistance enzyme that locally inactivates the drug (green shade) at a metabolic cost. When *N*_*R*_ *≥ N*_th_, the drug is inactivated in the entire deme and sensitive cells (red, *S*) benefit from the protection at no cost (e.g., bottom-left green shade). The fraction of *S* thus increases (solid red arrow). When *N*_*R*_ < *N*_th_, the drug hampers the spread of *S* (top-right red crosshairs) while *R*’s remain protected and thrive (blue arrow). In the mild environment (left, *K* = *K*_+_), *N*_*R*_ *→ N*_th_, whereas *N*_*S*_ *→ K*_+_ − *N*_th_ (solid red arrow). Similarly, in the harsh environment (right, grey background, *K* = *K*_−_), we still have *N*_*R*_ *→ N*_th_ while *N*_*S*_ *→ K*_−_ − *N*_th_ (blue arrow). *K* is assumed to switch suddenly between *K*_+_ and *K*_−_ (environmental variability), driving the deme size (*N* = *N*_*S*_ + *N*_*R*_) that fluctuates in time (Model & Methods, see S1 Appendix Fig S1 and Sec. 1.2.3). When *ν*_*±*_ ≲ 1 (intermediate switching), the deme experiences bottlenecks at every mild (*K* = *K*_+_) to harsh (*K* = *K*_−_) switch. When *K*_+_*/K*_−_ ≳ *N*_th_ [26] (Model & Methods), demographic fluctuations may cause the extinction (ext.) of *R* cells after each bottleneck (curved dotted red arrow). Bottom: Stochastic realisation of *N*_*S*_ (red) and *N*_*R*_ (blue) in a deme vs. time, with parameters *N*_th_ = 40 (dashed), *K*_+_ = 400, *K*_−_ = 80, *ν*_+_ = 0.075, and *ν*_−_ = 0.125 (Model & Methods). White/grey background indicates mild/harsh environment. Population bottlenecks (white-to-grey) enforce transient *N*_*R*_ dips (blue arrows) promoting fluctuation-driven *R* eradication (red arrow) [26] (Model & Methods). Panel B: Eco-evolutionary metapopulation dynamics; legend and parameters are as in A. The metapopulation is structured as a (two-dimensional) grid of connected demes, all with carrying capacity *K*(*t*) *∈ {K*_−_, *K*_+_*}* given by Eq. (3). Each *R* and *S* cell can migrate onto a neighbouring deme at rate *m* (curved thin arrows, Model & Methods). Owed to local fluctuations of *N*_*R*_, drug inactivation varies across demes (different shades). Bottlenecks can locally eradicate *R*, e.g. in deme (∗), but migration from a neighbouring deme (*†*) can rescue resistance (curved thin blue arrow). Resistance is fully eradicated when no *R* cells survive across the entire grid (curved dashed red arrow).

It has recently been shown that coupled environmental and demographic fluctuations shape the temporal evolution of cooperative antimicrobial resistance in well-mixed populations (e.g., in isolated demes), where the environmental conditions for the eradication of resistance were determined [26]. Cell migration between demes generally promotes the coexistence of strains in static environments [96, 39], and is thus expected to enhance cooperative drug resistance in the absence of environmental variability. In this context, we investigate how cell migration and environmental fluctuations influence the dynamics of antimicrobial resistance by asking: *(1) For what migration rates can environmental and demographic fluctuations clear resistance?* and *(2) What are the near-optimal environmental conditions ensuring the quasi-certain fluctuation-driven eradication of resistance in the shortest possible time?* Here, we answer these questions by combining analytical and simulation tools to determine under which circumstances the resistant strain goes extinct from the grid.

In the next section, we detail the model and introduce our main methods, including a background account of Refs. [26, 28]. We then present our main results answering the above central questions (1) and (2): we first determine the conditions ensuring the fluctuation-driven eradication of the cooperative resistant strain across the metapopulation, and then find the near-optimal conditions for the quasi-certain clearance of resistance in the shortest possible time. The biological relevance of these findings are then discussed alongside the model assumptions (robustness, limitations) and possible experimental impact, in light of the existing literature. Finally, we present our conclusions. Our study is complemented by a series of appendices and supporting movies (see S1 Appendix).

## Model & Methods

Motivated by *β-*lactamase cooperative antimicrobial resistance [43, 56, 74], and inspired by chemostat laboratory set-ups [25, 88, 93], we study how the migration of cells shapes the AMR eco-evolutionary dynamics in a *time-varying environment* of a spatially structured microbial population consisting of two cell types, denoted by *S a*nd *R*, competing for the same resources in the presence of a constant input of an antimicrobial drug to which *S c*ells are sensitive and *R m*icrobes are resistant, and against which the protection can be shared. Further details on the biological underpinning of our modelling approach are provided in the Discussion, and additional technical points can be found in S1 Appendix Sections 1-3.

### Metapopulation model

For the sake of concreteness, we consider a two-dimensional (2D) microbial metapopulation that can be envisioned as a grid of linear size *L*, containing *L*×*L d*emes (or sites) labelled by a vector 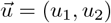 where *u*_1,2_ ∈ {1, 2, …, *L*} (with *L* = 20 in our examples), and periodic boundary conditions [75, 76, 77, 97, 18, 20]. The demes of this spatially explicit model are connected to their four nearest neighbours via cell migration, at per capita rate proportional to the migration parameter *m* (later simply referred to as “migration rate”), and are subject to a constant input rate of an antimicrobial drug [29, 30, 35]. Each deme 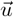 has the same carrying capacity, denoted by *K*, and at time *t* consists of a well-mixed subpopulation of 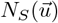 cells of type *S* that are drug-sensitive and 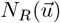 microbes of the AMR-resistant strain *R*, with deme size 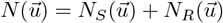, while the total number of *R*/*S* cells at time *t* across the metapopulation is 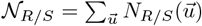 and the overall time-fluctuating number of microbes is 𝒩= 𝒩_*S*_ + 𝒩_*R*_, see Fig 1.

Antimicrobial resistance can often be seen as a form of cooperative behaviour [56, 43, 21], for example in the case of *β*-lactam antibiotics, microbes with resistance gene-bearing plasmids produce a *β*-lactamase resistance enzyme hydrolysing the antimicrobial drug in their surroundings [98, 99, 56, 43, 26, 28, 21]. In this context, when there are enough resistant microbes, the local concentration of resistance enzymes can reduce the drug concentration below the minimum inhibitory concentration (MIC), so that antimicrobial resistance acts as a public good, protecting both resistant and sensitive cells. Resistant microbes can either secrete the resistance enzymes (extracellular enzymes, typically produced by Gram-positive bacteria) or retain them within the cell (intracellular enzymes, Gram-negative bacteria) [100, 101]. Our theoretical model can account for both scenarios as it relies on the catalytic inactivation of the antimicrobial drug, either inside or outside *R* cells [56, 102, 103]. In any case, the public good (shared drug protection) can be interpreted as encoding the local decrease in the concentration of the active drug [56] (Introduction; see Fig 1).

Here, we consider a metapopulation model where *S* and *R* cells compete in each deme for finite resources in a *time-fluctuating environment* and in the presence of an antimicrobial drug. We assume that *R* cells share the benefit of drug protection with *S* cells within a deme 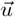 when the local number of resistant cells reaches or exceeds a certain fixed cooperation threshold *N*_th_ that is constant across demes, i.e., *R* cells act as cooperators in deme 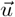 when 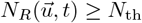 [26, 56, 43] (i.e., drug inactivation occurs at a faster timescale than microbial replication, each deme has a constant drug influx/outflux, and the deme volume is fixed; see *Extra and intracellular drug inactivation* in the subsection “Robustness of results across parameters and scenarios” of Discussion, and [28] for the case of a cooperation threshold set by the fraction of *R* cells in a single deme due to a time-varying volume). We therefore assume that microbes of the cooperative resistant strain *R* have the same constant growth fitness *f*_*R*_ = 1 − *s* in all demes, where the parameter *s* (with 0 < *s* < 1) represents a resistance-production metabolic cost. Moreover, cells of type *S* that are sensitive to the drug have a baseline fitness *f*_*S*_ = 1 when 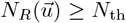, and *f*_*S*_ = 1 − *a* when 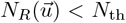, where *s* < *a* < 1, which denotes the growth fitness reduction caused by the drug [26, 28] (see *Population size and microbial parameter values* in the Discussion subsection “Robustness of results across parameters and scenarios” for further considerations on biostatic and biocidal drug action). Hence, with *f*_*R*_ − *f*_*S*_ = *a* − *s* > 0, the fitness of the *R* strain exceeds that of *S* when 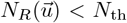, whereas the *R* type has a lower fitness than *S*, with *f*_*R*_ −*f*_*S*_ = −*s* < 0, when 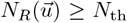. Both strains, *R* and *S*, are subject to the same single-deme carrying capacity (*K*), which is homogeneous across all demes of the metapopulation, at all times.

Here, we particularly focus on the metapopulation’s eco-evolutionary dynamics under slow migration, a biologically relevant dispersal regime known to increase population fragmentation and hence influence its evolution and diversity [104, 105, 106, 18, 107] (see *Population size and microbial parameter values* in Discussion for further details on the slow migration regime). Environmental variability is here encoded in the time-varying carrying capacity of each single deme, *K*(*t*), which changes simultaneously across all demes, and that we assume to switch endlessly between values representing mild and harsh conditions [51, 52, 53, 54], see Fig 1A and below. (A detailed discussion of the time-fluctuating *K*(*t*) is provided in *Environmental assumptions* in the Discussion subsection “Robustness of results across parameters and scenarios”).

### Intra- and inter-deme processes: Bacterial division, death, and cell migration

In close relation to the Moran process [108, 109, 110, 111], a reference model in biology for evolutionary processes in finite populations [112], the intra-deme dynamics within a lattice site 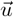 is represented by a birth-death process defined by the reactions 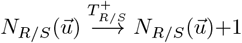 and 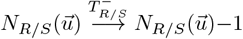 of birth (cell division) and death of *R*/*S* cells, occurring at local transition rates [51, 52, 53, 26, 27, 28, 79]

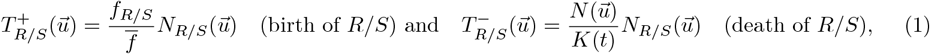

where 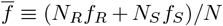 is the average fitness in deme 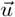 at time *t*. The continuous time variable *t* is measured in units of microbial replication cycles, i.e., microbial generations (see “Background” below and Results, as well as S1 Appendix Sec. 3).

The inter-deme dynamics on the 2D grid stems from the migration of one cell of *R*/*S* type from the site 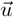 to one of its four nearest-neighbour demes denoted by 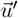. Cells’ dispersal in microbial populations is generally density-dependent, with movement often directed towards areas that are rich in resources [113], but simpler assumptions are commonly used [18, 20, 114, 79]. Here, we have considered two forms of migration: 1) We have first assumed a local density-dependent per-capita migration rate 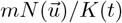, with increasing migration rate as the deme’s population size approaches the carrying capacity (less available resources in 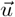). 2) We have also studied the simpler case of a constant per capita migration rate *m*, corresponding to the same dispersal in all spatial directions (symmetric migration) of all *R* and *S* cells in a deme 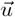. The inter-deme dynamics is therefore implemented by picking randomly a cell (*R* or *S*) from deme 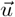 and moving it into a nearest-neighbour 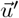 according to the reactions 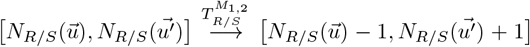 occurring at the migration transition rates

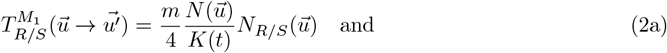

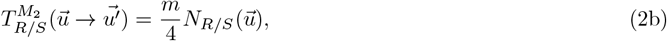

where 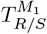 and 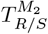 are the two forms of local migration rates (respectively, with density-dependent and density-independent per capita rate). We have found that the specific form of migration does not qualitatively affect our main findings, see Discussion. For notational simplicity, we may refer to *m* as the “migration rate”, being it clear from the context which of the two forms of migration, 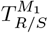 or 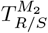, is used.

### Environmental variability

Microbial populations generally live in time-varying environments, and are often subject to conditions changing suddenly and drastically, e.g., experiencing cycles of harsh and mild environmental states [80, 55, 81, 41, 82, 83, 84, 50, 85, 86, 87, 88, 89, 90, 91, 92], see Fig 1A. Here, environmental variability is encoded in the time-variation of the binary carrying capacity [51, 52, 53, 78, 55, 54, 26, 28, 27, 79]

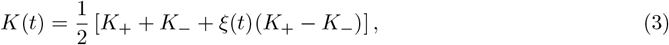

which can take the values *K*(*t*) ∈ {*K*_−_, *K*_+_} (see *Environmental assumptions* in the Discussion subsection “Robustness of results across parameters and scenarios” for further discussion and interpretation of *K*(*t*)). The carrying capacity is thus driven by the coloured dichotomous Markov noise (DMN), also called telegraph process, *ξ*(*t*) ∈ {−1, 1} that switches between ±1 according to *ξ* → −*ξ* at rate *ν*_*±*_ when *ξ* = ±1 [115, 116, 117]. It is convenient to write *ν*_*±*_ in terms of the mean switching rate *ν* ≡ (*ν*_−_ + *ν*_+_)/2 and switching bias *δ* ≡ (*ν*_−_ *ν*_+_)/(2*ν*), where |*δ*| ≤ 1, and *δ* > 0 indicates that, on average, more time is spent in the environmental state *ξ* = +1 than in *ξ* = −1 (*δ* = 0 corresponds to symmetric switching) [53, 54, 26, 28, 27]. In all our simulations, the DMN is at stationarity, and is therefore initialised from its long-time distribution, see S1 Appendix Sec. 3. All the DMN instantaneous correlations are thus time-independent while its auto-covariance reads ⟨*ξ*(*t*)*ξ*(*t*^*′*^)⟩ − ⟨*ξ*(*t*)⟩⟨*ξ*(*t*^*′*^)⟩ = (1 − *δ*^2^)*e*^−2*ν*|*t*−*t′*|^ [115, 116, 117, 53], where ⟨·⟩ denotes the ensemble average and 1/(2*ν*) is the finite correlation time (when *t, t*^*′*^ → ∞). Following Eq. (3), the carrying capacity switches back and forth at rates *ν*_*±*_ = *ν*(1 ∓ *δ*) between a value *K* = *K*_+_ (*ξ* = 1) corresponding to a mild environment, e.g., where there is abundance of nutrients and/or lack of toxins, and *K* = *K*_−_ < *K*_+_ (*ξ* = −1) under harsh environmental conditions (e.g., lack of nutrients, abundance of toxins) according to 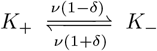, and thus describes (“feast and famine”) cycles of mild and harsh conditions. As the DMN, the time-fluctuating *K*(*t*) is always at stationarity: its expected value is 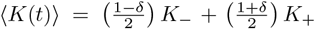, and its auto-covariance is 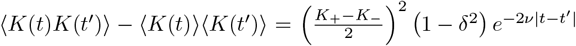 [116, 115, 117, 53]. Accordingly, in our simulations the initial value of the carrying capacity is drawn from its stationary distribution, i.e. *K*(0) = *K*_*±*_ with a probability (1 ± *δ*)/2, see S1 Appendix Sec. 3. The randomly time-switching *K*(*t*) drives the deme size of all demes simultaneously, and is hence responsible for the coupling of demographic fluctuations with environmental variability [51, 52, 53, 78, 55, 54, 27, 79]. This effect is particularly important when the dynamics is characterised by population bottlenecks [26, 28, 79]; see below and S1 Appendix Sec. 1.2.3.

The stochastic metapopulation model is therefore a continuous-time multivariate Markov process – defined by the transition rates given by Eqs. (1), (2a) and (2b) – that satisfies the master equation given in S1 Appendix Sec. 1.1. The individual-based dynamics encoded in Eqs. (1), (2a) and (2b) has been simulated using the Monte Carlo method described in S1 Appendix Sec. 3. It is worth noting that *N, N*_*R/S*_,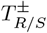 and 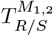 all depend on the deme 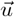, time *t*, and environmental state *ξ*. However, for notational simplicity, we often drop the explicit dependence on some or all of the variables 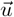, *t*, and *ξ*.

### Background: Eco-evolutionary dynamics in an isolated deme

Since the metapopulation consists of a grid of connected demes, all with the same carrying capacity *K*(*t*), it is useful to review and summarise the properties of the eco-evolutionary dynamics in a single isolated deme (when *m* = 0), studied in Ref. [26] (see also Refs. [51, 52, 28]). Further technical details can be found in S1 Appendix Sec. 1.2.

#### Mean-field approximation of the eco-evolutionary dynamics of an isolated deme

In an isolated deme, there is only cell division and death according to the intra-deme processes with rates given by Eq. (1). Upon ignoring all fluctuations, the mean-field dynamics in an isolated deme subject to a carrying capacity of constant and very large value *K* = *K*_0_ is characterised by the rate equations for the deme size *N* and the local fraction *x* ≡ *N*_*R*_/*N* of resistant cells [51, 52] (see details in S1 Appendix Sec. 1.2.1):

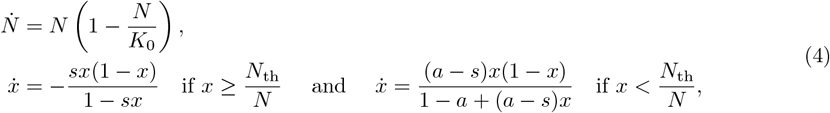

where the dot indicates the time derivative. The logistic rate equation for *N* predicts the relaxation of the deme size towards the carrying capacity *N* → *K*_0_ on a timescale *t* ∼ 1 (one microbial generation). The fraction of *R* cells is coupled to *N*: *x* decreases when *x* > *N*_th_/*N*, and increases otherwise (shared protection). Since 0 < *s* < *a* < 1, in this mean-field picture, *x* approaches *N*_th_/*K*_0_ on a timescale *t* ∼ 1/|*f*_*R*_ − *f*_*S*_| > 1 (*x* → *N*_th_/*K*_0_ and *N*_*S*_/*N* → 1 − *N*_th_/*K*_*0*_) [26, 28]. In our examples, we have |*f*_*R*_ −*f*_*S*_|∼ *s* ≪ 1 yielding a clear timescale separation between the dynamics of the deme size and its make-up: *N* and *x* are respectively the fast and slow variables; see S1 Appendix Sec. 1.2.1.

#### Eco-evolutionary dynamics of an isolated deme in a static environment (finite N and constant K)

An isolated deme of finite size, subject to a large and constant carrying capacity *K*_0_, can be aptly approximated by a Moran process by assuming that the deme size *N* = *K*_0_ is constant (see details in S1 Appendix Sec. 1.2.2) [108, 112, 51, 52, 26, 28]. In this static environment setting, *R* or *S* cells eventually take over and the process is characterised by the probability and mean time of fixation [108, 112, 109, 110]. Using classical techniques, the probability and mean time for the fixation of *R* and *S* can be computed exactly, showing that resistant cells are most likely to fix in the deme when the long-time fraction of *R* is high enough (*N*_th_/*K*_0_ ≳ ln (1 − *s*)/ ln (1 −*a*)) [26, 28]. Otherwise, *R* and *S* cells are likely to coexist for extended periods. Therefore, resistant cells generally persist in an isolated deme when the environment is static; see S1 Appendix Sec. 1.2.2. This picture is drastically altered when environmental variability generates strong population bottlenecks, as briefly reviewed below.

#### Eco-evolutionary dynamics in an isolated deme subject to a fluctuating environment

When the deme size is sufficiently large to neglect demographic fluctuations and randomness only stems from environmental variability via Eq. (3), the deme size dynamics is well approximated by the piecewise deterministic Markov process (*N*-PDMP) [118, 51, 52, 26, 28, 53, 78, 55, 27, 54, 79] defined by

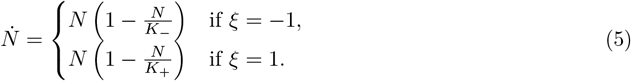

In the realm of the *N*-PDMP approximation, the deme size thus satisfies a deterministic logistic equation in each environmental state *ξ* = ±1, subject to the time-switching carrying capacity, see Eq. (3) (and details in S1 Appendix Sec. 1.2.3). The properties of the *N*-PDMP, defined by Eq. (5) and discussed in S1 Appendix Sec. 1.2.3, shed light on how the deme size changes with the rate of environmental changes. In particular, *N* tracks *K*(*t*) when the environmental switching is slower than the logistic dynamics (*ν* ≲ 1 and *δ* ≠ ±1). In this biologically relevant intermediate switching regime [119, 94], the deme experiences a *bottleneck* whenever the carrying capacity switches from *K*_+_ to *K*_−_ < *K*_+_ and its size is drastically reduced, see Fig 1A, with subpopulation prone to fluctuation-driven phenomena [50, 86, 87, 120, 51, 52, 10, 26, 28].

#### Fluctuation-driven resistance eradication in an isolated deme

When *ν* ∼ *s* ≲ 1 and 0 ≤ *δ* ≲ 1, the deme undergoes bottlenecks at an average frequency *ν*(1−*δ*^2^)/2, comparable to *s*, the rate at which the deme composition changes (more slowly than *N* that relaxes after *t* ∼ 1); see details in S1 Appendix Sec. 1.2.3. Assuming 1 ≪ *N*_th_ < *K*_−_ ≪ *K*_+_, the deme extinction is unlikely to be observed and, between two environmental switches, *N*_*R*_ and *N*_*S*_ fluctuate respectively about *N*_th_ and *K* − *N*_th_, with the deme consisting of a majority of *S* cells in the mild state (*K* = *K*_+_) (S1 Appendix Secs. 1.2.2, 1.2.3, and 5.4; see also Fig 1A and S1 Appendix Fig S8). Following each bottleneck, the coupling of *N* and *x* causes transient “dips” in the number of *R* cells [26], see Fig 1A, S1-S5 Movies and S1 Appendix Sec. 4. Using the *N*-PDMP approximation, it was shown that strong enough bottlenecks, whose strength is measured by *K*_+_/*K*_−_ (that can be roughly interpreted as the resource supply ratio in mild/harsh conditions in a chemostat set-up [55], or similarly as the inverse of an effective dilution factor), can eradicate resistance. Namely, when *K*_+_/*K*_−_ ≳ *N*_th_, demographic fluctuations are strong enough to lead to the extinction of *R* after a finite number of bottlenecks, i.e., in a time scaling as ∼1/*s* [26]. (In Fig 1A, *R* extinction occurs after four bottlenecks). This phenomenon where resistance is cleared by the coupled effect of environmental and demographic fluctuations is called “fluctuation-driven eradication”, and also holds for realistically large systems (e.g., *N* > 10^6^) [26] (Discussion and S1 Appendix Sec. 1.2.3). The ensuing resistance eradication, occurring under intermediate switching, where *ν* ∼ *s* ≲ 1 and 0 ≤ *δ* ≲ 1, is in stark contrast with the persistence of resistance characterising the regimes of slow and fast switching (*ν* ≪ 1 and *ν* ≫ 1); see S1 Appendix Sec. 1.2.3 and Sec. 2 Fig S3.

In this study, we investigate how the joint effects of migration, demographic fluctuations, and environmental variability influence the eco-evolutionary dynamics of the spatially structured metapopulation. We particularly focus on finding the conditions for the efficient clearance of drug resistance from the microbial community via *spatial fluctuation-driven eradication* of *R* cells.

## Results

In an isolated deme resistance is likely to persist for extended periods when the environment varies either quickly (*ν* ≫1) or slowly (*ν* ≪1) compared to the intra-deme dynamics (see S1 Appendix Sec. 1.2.3), whereas strong enough bottlenecks can cause *R* eradication in the regime of intermediate switching [26, 28] (*ν* ∼*s* ≲ 1, 0 ≤ *δ* ≤ 1, see “Background” in Model & Methods). However, determining the conditions for survival, fixation, or coexistence of strains in a metapopulation remains a challenging open problem. Since all demes of the metapopulation have the same time-switching carrying capacity *K*(*t*) given by Eq. (3), they have the same size distribution (S1 Appendix Sec. 1.2.3). The long-term coexistence of *R* and *S* across the grid is likely in the regimes of slow and fast environmental switching with non-zero migration, while *R* and *S* can take over under zero or slow migration (S1 Appendix Sec. 2 and Fig S3). This behaviour is similar to what happens in static environments (where *K* is constant), as discussed in S1 Appendix Sec. 2; see S1 Appendix Fig S2. By contrast, in the regime of intermediate switching, with *ν* ∼*s* ≲ 1 and 0 ≤ *δ* ≲ 1, all demes are subject to environmental bottlenecks [79] that can cause significant fluctuations in the number of *R* and *S* cells, and can eradicate *R* (see S1 Appendix Fig S6 and S7).

In this work, we focus on the intermediate switching regime where the size of each deme tracks its carrying capacity *K*(*t*) ∈ {*K*_−_, *K*_+_}, and strong bottlenecks can lead to fluctuation-driven eradication of resistance from the metapopulation (Fig 1); see below. We thus investigate under which conditions the coupled effect of environmental and demographic fluctuations leads to the clearance of resistance from the two-dimensional metapopulation. This is an important and intriguing question since microbial communities generally evolve in spatial settings, and locally *R*-free demes can be recolonized by cells migrating from neighbouring sites (Fig 1B). Migration is generally expected to favour diversity within demes by promoting the local coexistence of *R* and *S*, thus increasing alpha-diversity, while at the same time it reduces beta-diversity across the metapopulation, since all demes approach a similar composition of coexisting *R* and *S* cells [96, 39]; see S1 Appendix Fig S2. However, we show that fluctuation-driven eradication also occurs across the two-dimensional metapopulation and reveals a biologically relevant regime in which migration even *enhances* resistance clearance. Since *K*_+_/*K*_−_, referred to as the bottleneck strength, governs the fluctuation-driven eradication of *R* in an isolated deme [26] (Model & Methods), while *m* controls the homogenizing effect of dispersal [96, 39], their influences on the clearance of resistance are antagonistic. It is therefore enlightening to investigate the interplay between *K*_+_/*K*_−_ and *m* in determining the eradication of resistance.

Here, the metapopulation eco-evolutionary dynamics is studied by performing a large number ℛof long simulation runs (realizations) for each dataset, and its statistical properties are obtained by sampling all ℛ realizations; see S1 Appendix Sec. 3. In our simulations, the time is measured in “microbial generations” (or Monte Carlo steps), with one generation being the (mean) time for attempting 2𝒩 birth-death events (see S1 Appendix Sec. 3 for details). We choose a carrying capacity that is never too low, so that demes are always occupied by *R* and/or *S* individuals and the extinction of all cells in a deme is unobservable (S1 Appendix Secs. 1.1 & 3.1).

### Critical migration rate and bottleneck strength to eradicate antimicrobial resistance

To study under which circumstances environmental variability coupled to demographic fluctuations leads to the eradication of resistance in the two-dimensional metapopulation, we focus on the regime of intermediate environmental switching, with *ν* ∼ *s* ≲ 1 and 0 ≤ *δ* ≲ 1, and assume 1 ≪ *N*_th_ < *K*_−_ < *K*_+_. In this biologically relevant regime [119, 94] (see *Population size and microbial parameter values* in the Discussion subsection “Robustness of results across parameters and scenarios”), each deme experiences a sequence of bottlenecks of strength *K*_+_/*K*_−_, occurring at a rate *ν*(1 − *δ*^2^)/2, accompanied by “transient dips” in the number of cells, and the demes generally consist of a majority of *S* cells when it is in the mild state (*K* = *K*_+_) [26, 28] (“Background” in Model & Methods; see also S1 Appendix Secs. 1.2.3 and Fig S8 in Sec. 5.4). In addition, we consider a slow/moderate migration rate, with 0 < *m* ≲ 1 (see proper definitions just after Eq. (6)). The demes of the metapopulations are thus neither entirely isolated (*m* = 0), nor fully connected (*m* ≫ 1). We know that, in this intermediate environmental switching regime, fluctuation-driven eradication of *R* is likely to occur in isolated demes [26], i.e., in metapopulations with *m* → 0 (see S1 Appendix Secs. 1.2.3 and 5.1), while the probability of long-lived coexistence of *R* and *S* is expected to increase with *m* (the latter as in S1 Appendix Sec. 2 Figs S2 and S3, for constant and very slow/fast switching environments, respectively). In this context, when 0 < *m* ≲ 1, fluctuations can clear resistance in some demes, but these *R*-free demes can be recolonised following migration events from neighbouring sites still containing *R* cells; see Figs 1B, 2 and 3, and S1 Appendix Sec. 5.1 Figs S4 and S5. The effect of fluctuations caused by environmental bottlenecks is thus countered by migration, and it is not obvious whether eradication of resistance can arise in the spatial metapopulation.

We ask for what migration rates can environmental and demographic fluctuations clear resistance. The first central question that we address is therefore *whether there is a critical migration rate m*_*c*_ *above which the fluctuation-driven eradication of resistance is unlikely, and below which it is either possible or likely*. Since the amplitude of the fluctuations generated by the bottlenecks increases with their strength, we expect *m*_*c*_ to be an increasing function of *K*_+_/*K*_−_.

We have computed the probability *P*(*N*_*R*_(*t*) = 0) that there are no resistant cells across the entire metapopulation after a time *t* (by sampling ℛ realizations, see S1 Appendix Sec. 3). In Fig 2A we report *P*(*N*_*R*_(*t*) = 0) as a function of bottleneck strength *K*_+_/*K*_−_ and migration rate *m* when (*ν, δ*) = (1, 0.75), finding *P*(*N*_*R*_(*t* ≫ 1) = 0) ≈ 1 when *m* is below a certain value. Similar results are found in Fig 4B-D for other environmental parameters (*ν, δ*) at different times *t*. This indicates the existence of a trade-off between the rate of migration and bottleneck strength, see Figs 2A and 4A-D: For a given bottleneck strength *K*_+_/*K*_−_, when the migration rate is below the critical value *m*_*c*_, shown as red/dark phases in Figs 2A and 4A-D, the fluctuations caused by bottlenecks can clear resistance across the whole metapopulation in a finite time (that scales with 1/*s*, see below). A few bottlenecks thus suffice to eradicate *R*, as shown in, e.g., Fig 2C and S1 Appendix Fig S6C,D,F, where each red spike corresponds to a bottleneck (see “Breaking it down” subsection below). An approximate expression for *m*_*c*_ is obtained by matching the total number of *R* migration events across the metapopulation, during the time between two successive bottlenecks, with the number of new *R*-free demes due to a bottleneck, yielding (see the derivation at the end of this subsection)

**Figure 2.**
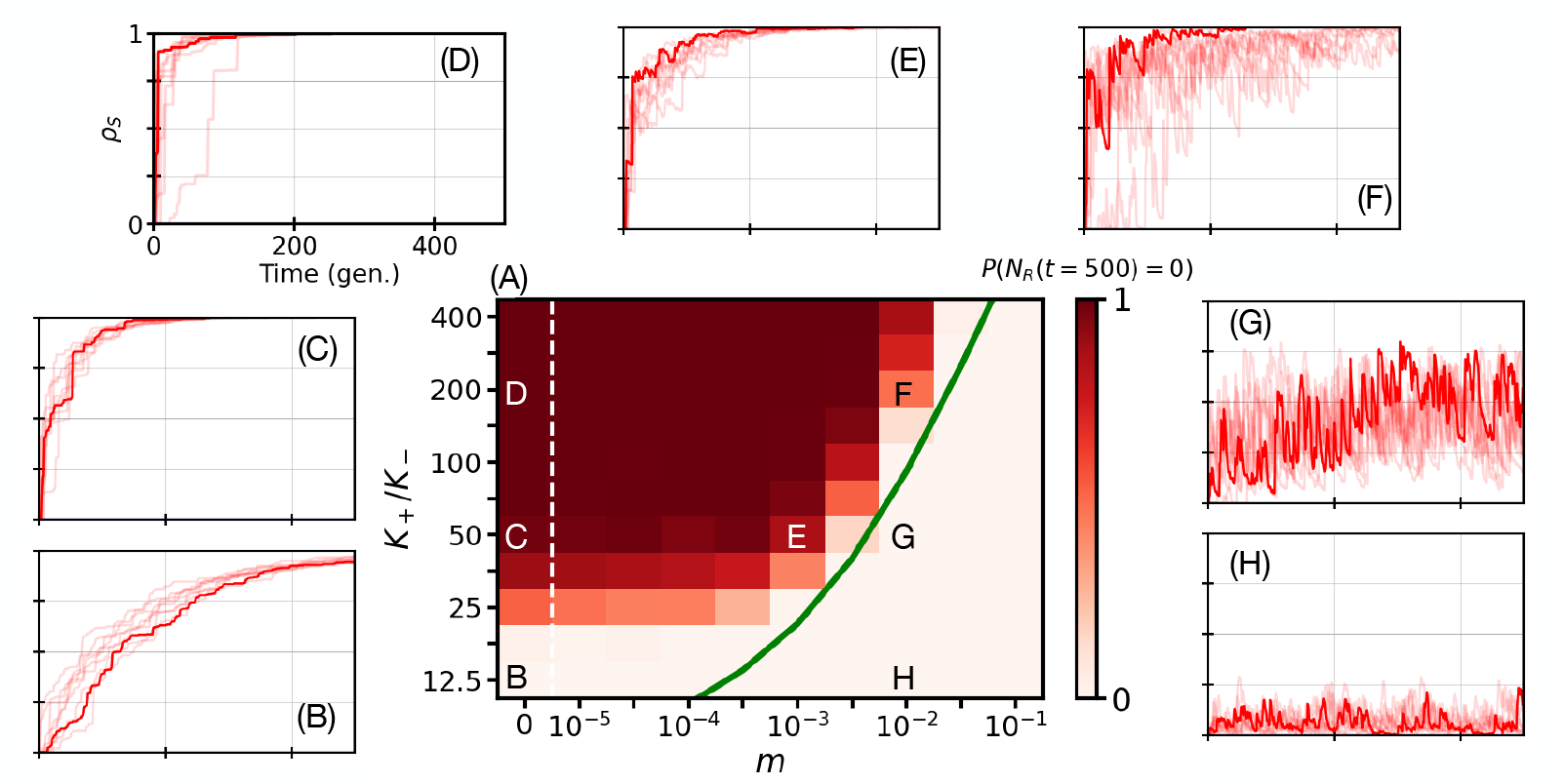
The eradication of *R* cells depends on the bottleneck strength and migration rate. The shared parameters in all panels are *ν* = 1, *δ* = 0.75, *L* = 20, *a* = 0.25, *s* = 0.1, *N*_th_ = 40, and *K*_−_ = 80 (Model & Methods) with migration according to Eq. (2a). Other parameters are as listed in Table 1. Panel A: Heatmap of the probability *P* (*N*_*R*_(*t*) = 0) of total extinction of *R* (resistant) cells as a function of bottleneck strength, *K*_+_*/K*_−_, and migration rate *m* at time *t* = 500. Each (*m, K*_+_*/K*_−_) value pair represents an ensemble average of *R* = 200 independent simulations, where we show the fraction of realisations resulting in complete extinction of *R* (resistant) microbes after 500 microbial generations (standard error of the mean in *P* (*N*_*R*_(*t* = 500) = 0) below 4%; see S1 Appendix Sec. 3.3.2). The colour bar ranges from light to dark red, where darkest red indicates complete *R* extinction in all 200 simulations at time *t* = 500, *P* (*N*_*R*_(*t* = 500) = 0) = 1. The green line is the theoretical prediction of Eq. (6) and the white dashed vertical line indicates an axis break separating *m* = 0 and *m* = 10^−5^ (Model & Methods). The black and white annotated letters point to the specific (*m, K*_+_*/K*_−_) values used in the outer panels. Panels B-H: Typical example trajectories of the fraction of demes *ρ*_*S*_ (*t*) without *R* cells, up to *t* = 500 microbial generations (gen.), defined by Eq. (7) and corresponding to the fixation of *S* in the metapopulation. (The fraction of demes without *S* cells, *ρ*_*R*_(*t*), is vanishingly small and unnoticeable.) Here, *ρ*_*S*_ (*t*) is shown as a function of time (microbial generations) for the (*m, K*_+_*/K*_−_) value pairs indicated in Panel A (see S1 Appendix Sec. 3).

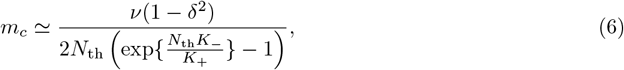

**Table 1.** Summary of simulation parameters for Figures 2-4 and S1-S5 Movies. Parameters kept fixed are listed by a single value, other parameters are listed as ranges. The average number of sensitive cells *S* per deme at 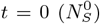 equals the metapopulation’s carrying capacity at *t* = 0 minus the constant threshold value for cooperation, *K*(*t* = 0) − *N*_th_, which depends on whether the system begins in a harsh or mild environment, *K ∈ {K*_+_, *K*_−_*}* (S1 Appendix Sec. 3.1). See S1 Appendix Fig S10 (as well as Fig S3) for the extended range in *ν* and *δ*, and S1 Appendix Sec. 5.5 for the discussion of results obtained on a periodic one-dimensional lattice (cycle) of length *L* = 100.

| Parameter | Description | Value |
| --- | --- | --- |
| $L$ | side length of square lattice (of size $L^2$) | 20 |
| $t_{max}$ | maximum number of microbial generations | 500 |
| $N_S^0$ | average number of $S$ cells per deme at $t = 0$ | $K(t = 0) - N_{th}$ |
| $N_R^0$ | average number of $R$ cells per deme at $t = 0$ | $N_{th}$ |
| $a$ | reduction in the birth rate of $S$ cells due to drug exposure | 0.25 |
| $s$ | resistance metabolic cost for $R$ cells | 0.1 |
| $m$ | migration rate | $0-10^{-1}$ |
| $K_+$ | carrying capacity per deme in the mild environment | $10^3-3.2 \cdot 10^4$ |
| $K_-$ | carrying capacity per deme in the harsh environment | 80 |
| $N_{th}$ | cooperation threshold | 40 |
| $\nu$ | environmental switching rate | $10^{-3}-10^1$ |
| $\delta$ | environmental switching bias | $0-0.75$ |

whose graph is shown in the green curve of Fig 2A where it approximately captures the border between the red/white phases and how *m*_*c*_ increases with *K*_+_/*K*_−_ (see also Fig 4A-D where *m*_*c*_ ranges from 10^−4.5^ to 10^−2^). Here, the regime of slow migration is defined by *m* ≲ *m*_*c*_ (with moderate migration when *m* ≳ *m*_*c*_).

Further light into the phenomenon of fluctuation-driven *R* eradication is shed by computing the fraction of demes *ρ*_*S/R*_(*t*) that consist only of *S*/*R* microbes at time *t*:

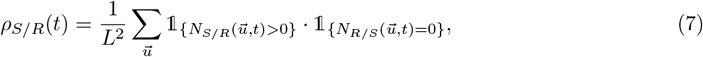

where 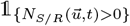 is the indicator function defined as 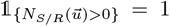 if 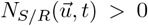 and 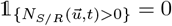 otherwise. Since each deme is never empty, *ρ*_*S/R*_(*t*) also corresponds to the fraction of demes without any *R*/*S* cells, i.e. *ρ*_*S*_(*t*) thus gives the fraction of *R*-free demes across the metapopulation at time *t*. In Figures 2 and 3, and those of S1 Appendix, *ρ*_*S/R*_(*t*) correspond to the fraction of *R*/*S*-free demes in a *single realization* of the metapopulation. In the results of Fig 2B-H (and 3I), *ρ*_*S*_(*t*) increases sharply coincidentally with each bottleneck and then transiently decreases due to the recolonisation of *R*-free demes via migration, whereas *ρ*_*R*_(*t*) → 0 at all times. When bottlenecks are strong and *m* ≲ *m*_*c*_ (*K*_+_/*K*_−_ and *m* in the red/dark phase of Fig 2A), recolonisation cannot counter bottlenecks and eventually *ρ*_*S*_(*t*) → 1 with the eradication of resistance from all demes; see Figs 2C-E and 3I, and S1 Appendix Figs S5E-H, S6C,D,F, and S7D,F, while *ρ*_*R*_(*t*) → 0 (*R*-only demes are very unlikely, but see the blue lines in S1 Appendix Figs S5D, S6E and S7C,E when *m* → 0). When *m* > *m*_*c*_ (without *K*_+_/*K*_−_ scaling as the system size, see below), *ρ*_*S*_(*t*) remains finite, while generally *ρ*_*R*_(*t*) → 0 regardless of *m*. This indicates the persistence of *R* in the metapopulation, which then consists of demes where *R* and *S* coexist, and other demes that are resistance free (*S*-free demes are very unlikely, *ρ*_*R*_(*t*) → 0, when *m* > *m*_*c*_; see S1 Appendix Fig S5I-L).

**Figure 3.**
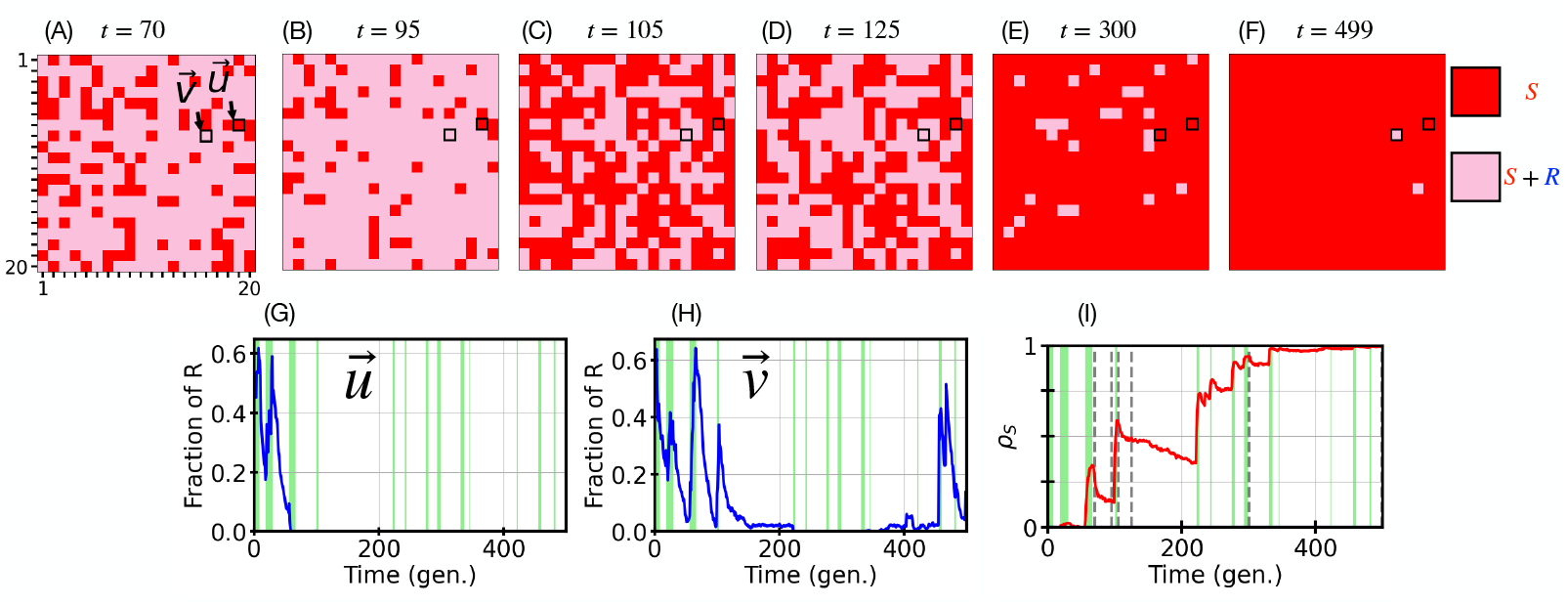
A closer look to individual demes: Migration and intermediate environmental switches shape local eradication of *R* cells. Example eco-evolutionary dynamics of the metapopulation in a single simulation realisation. Parameters are *K*_+_ = 2000, *ν* = 0.1, *δ* = 0.5, and *m* = 0.001, with density-dependent migration according to Eq. (2a); other parameters are as in Table 1. Panels A-F: Snapshots of the 20 *×* 20 metapopulation at six microbial generation times *t ∈ {*70, 95, 105, 125, 300, 499*}*. Red pixels indicate *R*-free demes (containing only *S* cells) and pink pixels are demes where *R* and *S* cells coexist. The two demes, 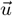 and 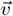, whose time composition is tracked in Panels G and H are indicated by a black border. Panel A shows the metapopulation a few generations after an environmental bottleneck. From panels A to B no bottleneck occurs, and many *S*-only demes are recolonised by *R* cells (many red pixels become pink). Between B and C, the metapopulation experiences a bottleneck causing a burst of local *R* extinctions (with burst of randomly located red pixels, see also the spike of *ρ*_*S*_ (*t* = 105) in Panel I). Panel D: Pink clusters spread across the grid due to the migration of *R* cells causing many recolonisation events (*ρ*_*S*_ (*t*) in Panel I decreases for *t ∈* [105, 125]). Panels E-F: After a sequence of bottlenecks starting at *t ≈* 220, the number of *S*-only demes increases overwhelmingly across the grid (*ρ*_*S*_ (*t* ⪅ 220) *→* 1 in Panel I), and resistance persists only in a few demes where *R* and *S* coexist. See S3 Movie and S1 Appendix Sec. 4 for a video of the full spatial metapopulation dynamics for this example realisation and its detailed description. Panels G-H: Temporal evolution of the fraction of resistant cells 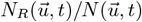 and 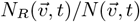, up to *t* = 500 microbial generations (gen.), in the example demes 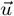 and 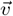 indicated as highlighted pixels in Panels A-F. Green bands indicate periods in the harsh environment (where *K*_−_ = 80); harsh periods shorter than 1 microbial generation are not shown (S1 Appendix Sec. 3.3.1). Each transition from white background to a green band indicates an environmental bottleneck. The deme 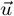 of panel G first exhibits *R/S* coexistence, followed by fluctuation-driven *R* eradication at *t* ≃ 70 due to environmental bottlenecks. In Panel H, similar dynamical development is followed by the restoration of resistance through recolonisation of the deme by *R* cells, as indicated by the blue spikes at long times (*t ≈* 350, Discussion). Panel I: Temporal evolution of the fraction *ρ*_*S*_ (*t*) of demes without *R* cells (red pixels within Panels A-F, see Eq. (7)). From left to right, the dashed vertical lines indicate the corresponding snapshot times in Panels A-F. Green background areas as in Panels G-H.

**Figure 4.**
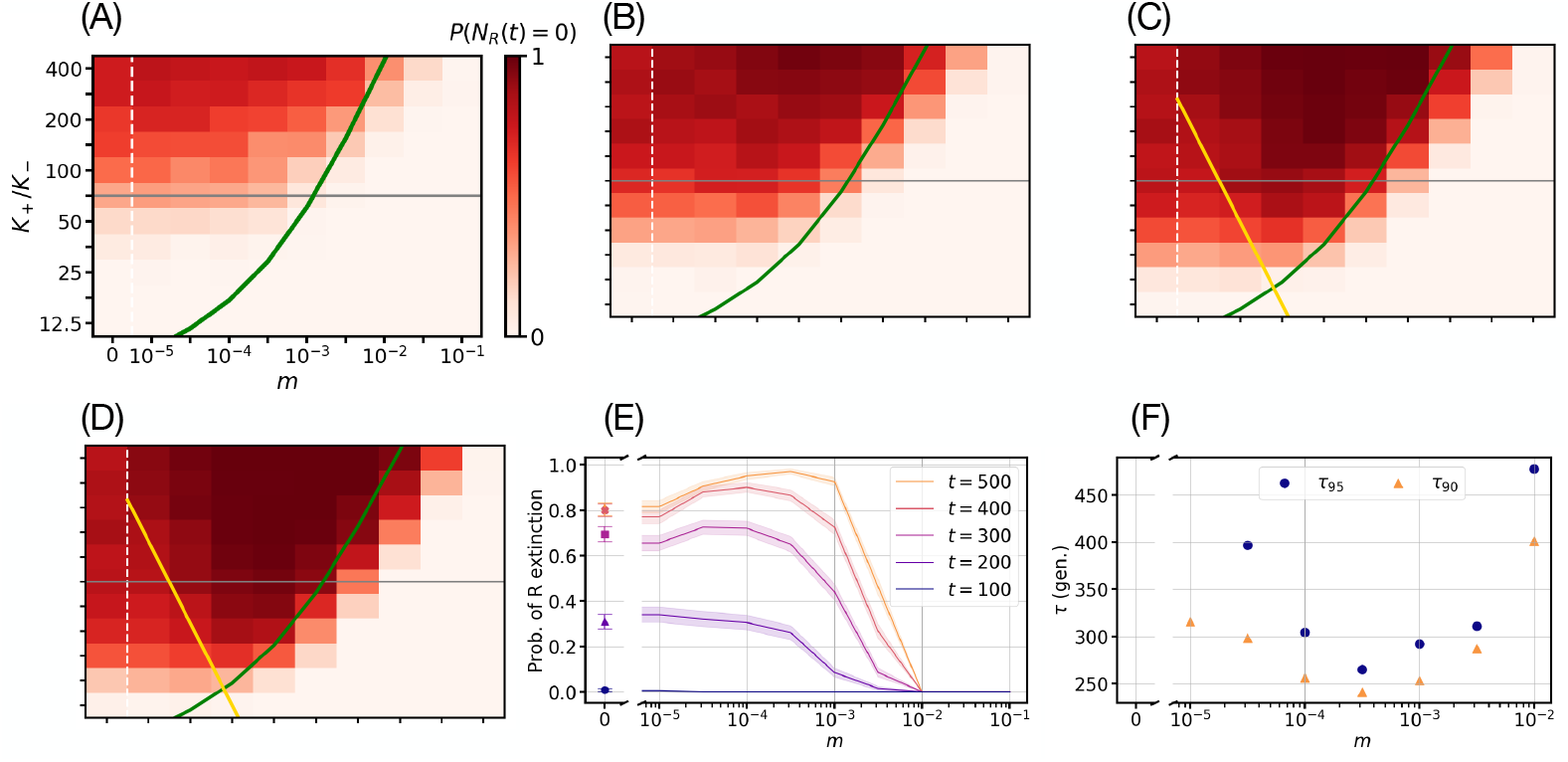
Near-optimal conditions for resistance clearance: Slow migration can speed up and enhance the eradication of *R* cells. Temporal evolution of the heatmap showing the probability *P* (*N*_*R*_(*t*) = 0) of *R* extinction as a function of bottleneck strength, *K*_+_*/K*_−_, and migration rate *m* (implemented according to Eq. (2a)) at *t* = 200 (Panel A), *t* = 300 (Panel B), *t* = 400 (Panel C), and *t* = 500 (Panel D) with environmental switching rate *ν* = 0.1 and bias *δ* = 0.5; other parameters are as in Table 1. As in Fig 2A, each (*m, K*_+_*/K*_−_) value pair is an ensemble average over 200 independent metapopulation simulations and the *P* (*N*_*R*_(*t*) = 0) colour bar ranges from light to dark red indicating the fraction of simulations that have eradicated *R* cells at each snapshot in time (standard error of the mean in *P* (*N*_*R*_(*t*) = 0) is below 4%; see S1 Appendix Sec. 3.3.2). The green and dashed white lines represent the theoretical prediction of Eq. (6) and an eye-guiding axis break, respectively (as in Fig 2A). The golden lines in Panels D-E show 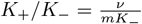, with *P* (*N*_*R*_(*t*) = 0) *≈* 1 in the (upper) region between the golden and green lines, according to Eq. (9). The grey horizontal lines in Panels A-E indicate the example bottleneck strength used in Panel E. Panel E: Probability of *R* extinction *P* (*N*_*R*_(*t*) = 0) as a function of migration rate *m* at bottleneck strength *K*_+_*/K*_−_ = 70.7 for *t* = 100, 200, 300, 400, 500 microbial generations (bottom to top). Solid lines (full symbols at *m* = 0) show results averaged over 200 realisations; shaded areas (error bars at *m* = 0) indicate binomial confidence interval computed via the Wald interval (see S1 Appendix Sec. 3.3.2). Panel F: 90th and 95th percentile (*τ*_90_ and *τ*_95_ respectively) of *R* eradication times as function of the migration rate with a bottleneck strength *K*_+_*/K*_−_ = 400 (see S1 Appendix Sec. 3.3.3). Panel F shows a single minimum at *m*^∗^ *≈* 10^−3.5^ corresponding to *τ*_90*/*95_(*m*^∗^) = *τ* (*m*^∗^) = *t*^∗^ *≈* 240 − 270.

The fluctuation-driven eradication of *R* across the two-dimensional metapopulation hence requires intermediate environmental switching, strong enough bottlenecks, and slow migration, which can be summarised by the necessary conditions

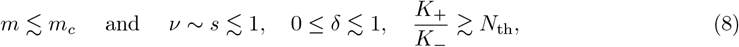

where the first condition indicates that the demes are not fully connected. In the limit of fast migration, here defined as *m* ≫ *m*_*c*_ (see below), the metapopulation can be regarded as *L*^2^ fully connected demes (island model [121, 122]), all subject to the same fluctuating carrying capacity given by Eq. (3). The fraction of *R* cells just after an environmental bottleneck still fluctuates about *x* = *N*_th_/*K*_+_ in each deme (same *x* as in the mild environment, see Eq (4)). All *L*^2^ demes experience the same carrying capacity *K*_−_ during the bottleneck. The approximate total number of *R* cells across the metapopulation right after a bottleneck is thus 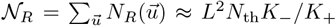. When *m* ≫ *m*_*c*_, resistance can typically be eradicated only if all *R* cells can be eliminated simultaneously during a single bottleneck. This is because *m* ≫ *m*_*c*_ ensures quick and efficient deme mixing between each bottleneck: If *R* is not eradicated from each deme after a single strong bottleneck, fast migration restores resistance in all demes before the next bottleneck (as opposed to the case of slow migration, where *R*-free demes can gradually accumulate after several consecutive bottlenecks). The eradication of *R* can occur under fast migration when 𝒩_*R*_ ≲ 1 following a single bottleneck, i.e. for 𝒩_*R*_ ≈ *L*^2^*N*_th_*K*_−_/*K*_+_ ≲ 1. Hence, when *ν* ∼ *s* ≲ 1 and 0 ≤ *δ* ≲ 1, fluctuation-driven eradication of resistance occurs for very strong bottlenecks, *K*_+_/*K*_−_ ≳ *N*_th_*L*^2^, regardless of the actual value of the migration rate.

#### Derivation of the critical migration rate m_c_

To derive Eq. (6), we remember that the fluctuation-driven eradication of *R* in a deme arises when *K*(*t*) switches between *K*_+_ and *K*_−_ at rate *ν* ∼ *s* ≲ 1 (with 0 ≤ *δ* ≲ 1), generating strong enough population bottlenecks (*K*_+_/*K*_−_ ≳ *N*_th_); see the end of the “Background” subsection in Model & Methods. In this regime, the number of microbes in each deme (*N*) continuously tracks the same carrying capacity *K*(*t*) on a fast timescale *t* ∼ 1, while each deme’s composition (*x*) changes on a slower timescale *t* ∼ 1/*s*. After each bottleneck, the local fraction of *R* cells initially fluctuates about *x* ∼*N*_th_/*K*_+_ ≪ 1, and their expected number in the harsh environment, *N*_*R*_ *≈ N*_th_*K*_−_/*K*_+_ ≲ 1, is sufficiently low for demographic fluctuations to effect the eradication of resistance [26] (“Background” in Model & Methods).

We assume that, in each deme, approximately *K*_−_ cells are randomly drawn to survive a bottleneck. Since there is a large number of cells before the onset of a bottleneck (*K*_+_ ≫ 1), each *R* cell has the same independent probability to survive the bottleneck (random draws with replacement), from a deme consisting of an approximate fraction *x* ≈ *N*_th_/*K*_+_ of *R* cells [26] (Model & Methods). Therefore, the approximate number of *R* cells surviving one bottleneck can be drawn from a Poisson distribution of mean *N*_th_*K*_−_/*K*_+_, and we thus estimate the probability that a bottleneck eradicates resistance as exp 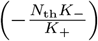. In this regime, the fluctuation-driven clearance of AMR is attempted at each bottleneck, see Fig 1A. AMR fluctuation-driven eradication thus occurs at the average bottleneck frequency *ν* (1 − *δ*^2^ )/2. Consequently, the rate at which each deme becomes *R*-free is approximately 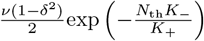.

The demes of the metapopulation are connected by cell migration, which generally homogenizes the local population make-up [123] and here tends to favour the coexistence of *R* and *S* cells [96, 39] (see S1 Appendix Sec. 2). Noting that the fraction of demes where resistance survives a single bottleneck is approximately 1−exp 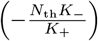, and that the number of surviving *R* cells in a deme tends to *N*_th_ (see Eq. (4), Model & Methods, [26]), the estimated rate of migration of *R* cells from each of these demes is *mN* 1 – exp 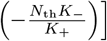. Matching this *R* cell migration rate with the rate 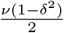 exp 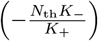 at which a deme becomes *R*-free corresponds to migration and fluctuation-driven eradication balancing each other, and hence yields the expression of Eq. (6) for the critical migration rate *m*_*c*_.

If *m* ≫ *m*_*c*_, unless *K*_+_/*K*_−_ ≳ *N*_th_*L*^2^ (see after conditions (8)), migration generally promotes long-time *R* and *S* coexistence (see above and S1 Appendix Figs S3, S4D, and S5I-L). Moreover, when *m* ≪ *m*_*c*_, the fluctuation-driven eradication of *R* is essentially the same as in an isolated deme (*m* = 0), see Fig 4A-E and below. We note that Eq. (6) and its derivation are independent of the spatial dimension of the metapopulation (see *Impact of the spatial dimension and accuracy of the critical migration prediction* in the Discussion subsection “Robustness of results across parameters and scenarios” for further details; and S1 Appendix Sec. 5.5 and Fig S9 for the case of a one-dimensional metapopulation). It is also worth noting that the conditions (8) are essentially independent of the spatial dimension of the metapopulation and hence the fluctuation-driven eradication of resistance is a phenomenon expected to hold on metapopulation lattices of any spatial dimension, see *Impact of the spatial dimension and accuracy of the critical migration prediction* in Discussion and S1 Appendix Sec. 5.5.

### Breaking it down: bottlenecks and fluctuations vs. spatial mixing

To further understand the joint influence of bottlenecks and migration on *R* eradication, we analyse typical single realisations of the metapopulation spatio-temporal dynamics when it is subject to intermediate switching rate and slow migration (*m* < *m*_*c*_), and experiences bottlenecks of moderate strength; see Fig 3 where the parameters *ν* = 0.1, *m* = 0.001, and *K*_+_/*K*_−_ = 25 (see also S3 Movie and S1 Appendix Sec. 4) satisfy the *R* fluctuation-driven eradication conditions of (8).

This resistance clearance mechanism, driven by bottlenecks and fluctuations, occurs randomly across the grid (scattered red sites in Fig 3A). The microbial composition of each deme fluctuates due to the homogeneous environmental variability (*K* switches simultaneously in time across all demes of the grid) and random birth-death events. In the regime defined by the conditions (8), strong bottlenecks cause demographic fluctuations that, after enough time, lead to *R* eradication in some demes (e.g., after *t* = 70 in Fig 3A,G). However, resistant cells can randomly migrate from neighbouring demes, recolonising *R*-free demes and favouring the spread of microbial coexistence across the metapopulation (pink clusters in Fig 3A-B). The fraction 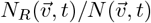 of *R* cells in a deme 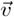 recolonised by resistance is characterised by spikes after a period of extinction, corresponding to *R* recolonisation events (e.g., at *t* ≈ 350 in Fig 3H, see also pixel 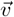 in Fig 3A-E). In summary, the occurrence of bottlenecks increases the fraction *ρ*_*S*_(*t*) of *R*-free demes across the grid (one spike in Fig 3I at each bottleneck), whereas *R* recolonisation gradually reduces *ρ*_*S*_(*t*), leading to a sequence of spikes and decreases of ρ_*S*_(*t*) (Fig 3I). Spikes are higher the stronger the bottlenecks (larger *K*_+_/*K*_−_), while the decrease of *ρ*_*S*_(*t*) steepens for faster migration (higher values of *m*). The typical stages of *ρ*_*S*_(*t*) dynamics are thus: (i) an environmental bottleneck eradicates *R* in some demes causing *ρ*_*S*_(*t*) to spike, (ii) some of these demes are then recolonised by *R* cells through migration, and *ρ*_*S*_(*t*) decreases. This is then followed by another bottleneck, that restarts the cycle of spikes and decreases of *ρ*_*S*_(*t*). The succession of steps (i) and (ii) as the environment switches back and forth, eventually leads to either *R* eradication (when *m* < *m*_*c*_) or to the persistence of resistance (long-term balance of spikes and decreases of *ρ*_*S*_). In the example of Fig 3 for a single realization of the metapopulation, the number of *R*-free demes steadily increases with the number of bottlenecks and, migration not being fast enough to restore resistance across the grid, *R* cells are eventually eradicated from the entire metapopulation (increasingly more red *R*-free demes in Fig 3E-F than in Fig 3A-D; *ρ*_*S*_(*t*) → 1 when *t* ≳ 400 in Fig 3I; see also S3 Movie and S1 Appendix Sec. 4). This is consistent with the metapopulation realisation of Fig 3 satisfying the conditions (8). See S1 Appendix Sec. 5.4 and Fig S8 for the dynamics of the absolute number of *S* and *R* cells in this example realisation of Fig 3.

When sensitive and resistant cells locally coexist on the grid, the demes containing *R* and *S* microbes (pink pixels in Fig 3A-F) consist approximately of *N*_th_ and *K*(*t*) − *N*_th_ cells of type *R* and S, respectively (see S1 Appendix Sec. 5.4 Fig S8). In the examples considered here, prior to a bottleneck, coexisting demes in the mild environment (where *K* = *K*_+_ ≫ *N*_th_) are thus made up of an overwhelming majority of *S* cells, see S1 Appendix Fig S8 and S2-S4 Movies, which is consistent with an *R* “containment strategy” [124].

### Slow migration can speed up and enhance *R* eradication: Near-optimal conditions for resistance clearance

We have disentangled the trade-off between the population bottleneck strength *K*_+_/*K*_−_ and migration rate *m*. To further clarify the interplay between fluctuations and migration in the metapopulation, we investigate how the probability *P*(*N*_*R*_(*t*) = 0) that there are no resistant cells across the entire grid after a time *t* depends on *m* and *K*_+_/*K*_−_ over time (Figs 4), and how it changes for different values of the switching rate (S1 Appendix Sec. 5.6 Fig S10). Under the conditions of (8), the probability *P*(*N*_*R*_(*t*) = 0) of overall resistance eradication increases in time (red/dark phases in Fig 4A-D): the *R* eradication mechanism driven by strong bottlenecks overcomes microbial mixing, and the red/dark phase expands in time until reaching its border where *m* ≈ *m*_*c*_ (see Eqs. (6) and (8)).

Remarkably, in Fig 4 we find that after some time (*t* ≳ 200), *R* cells are most likely to be eradicated from the metapopulation under slow but non-zero migration (in Fig 4B-D red regions are darker for *m* ∼ 10^−4^ − 10^−3^ than *m* ∼ 0 − 10^−4.5^; see S1-S2 Movies and S1 Appendix Secs. 4, and S1 Appendix Sec. 5.1 Figs S4 and S5). In Fig 4E, the probability of *R* eradication *P*(*N*_*R*_(*t*) = 0) for *t* ≥ 300 increases steadily with *m* before reaching a plateau near 1 for *m* ∼ 10^−4^ − 10^−3^ (*P*(*N*_*R*_(*t* ≥ 300) = 0) ≳ 0.7 in Fig 4E), and then sharply decreases as *m* exceeds *m*_*c*_. Since *K*_+_ ≫ *K*_−_ ≫ 1, most migration events in the intermediate regime defined by the conditions of (8) occur when demes are in the mild environmental state, where the number of microbes in each deme is typically large: Thus, *N* ≈ *K*_+_ and most individuals are of type *S*, with *N*_*S*_ *≈ K*_+_ − *N*_th_ ≫ *N*_*R*_ ≈ *N*_th_ ≫ 1 (Model & Methods and Fig 1A, and S1 Appendix Sec. 5.4 Fig S8). We hence estimate that the rate of migration per deme in the switching regime *ν* ∼ *s* ≲ 1 (see conditions (8)) is roughly *mK*_+_, and consists mostly of sensitive individuals moving into a neighbouring deme. In this context, the impact of migration is particularly significant for the eradication of *R* cells when the rate of cell migration per deme (mostly of *S* during the mild environmental state), approximately *mK*_+_, is comparable to the rate *ν* (1 − *δ*^2^)/2 at which bottlenecks arise (Model & Methods; see also S1 Appendix Sec. 1.2.1). In fact, when *mK*_+_ ≳ *ν*, the *R*-dominated demes (that have by chance been taken over by *R*) can be efficiently recolonised by *S* cells, and can then be eventually cleared from resistance by the fluctuation-driven mechanism caused by strong bottlenecks, as illustrated in S1 Appendix Sec. 5.1 and Figs S4C and S5E-H. Matching the rates at which bottlenecks and the *S*-recolonisation of *R*-dominated demes occur, yields the condition *m* ≳ *ν*/*K*_+_ for which migration can efficiently help promote the fluctuation-driven clearance of resistance; see yellow lines in Figs 4C-D. *R*-dominated demes are not effectively recolonised when the migration rate is lower than *ν*/*K*_+_, and therefore the probability of *R* eradication when *m* < *ν*/*K*_+_ is the same as for *m* = 0 (see *Population size and microbial parameter values* in the Discussion subsection “Robustness of results across parameters and scenarios”; see also S1 Appendix Sec. 5.1).

The probability *P*(*N*_*R*_(*t*) = 0) of *R* eradication for *m* ≲ *m*_*c*_ is an increasing function of *t* at fixed migration rate (Fig 4E). In fact, as this environmental regime is characterised by a sequence of strong bottlenecks, each of which can be seen as an attempt to eradicate *R* (“Background” in Model & Methods), the clearance of resistance for any 0 < *m* ≲ *m*_*c*_ is certain in the long run, i.e. *P*(*N*_*R*_(*t* →∞) = 0) →1. However, maximising the probability clearance of resistance in the shortest possible time is of great biological and clinical significance, e.g. to devise efficient antibacterial treatments [41, 32, 125]. This means that it is important to determine when the eradication of resistance is both *likely and rapid*. The second central question that we ask is therefore: *What are the conditions ensuring a quasi-certain clearance of resistance in the shortest possible time t*^∗^*?*

To address this important problem, we have determined the migration rate *m*^∗^, satisfying the conditions (8), for which the time for the eradication of *R*, here denoted by *t*^∗^, is minimal. As detailed below, in Fig 4F we determine *m*^∗^ and *t*^∗^, corresponding to the near-optimal conditions for the clearance of resistance, for the example of Fig 4 when the bottlenecks strength is *K*_+_/*K*_−_ = 400 (largest value considered in Fig 4). To this end we have computed *τ*_90_(*m*) ≡ min_*t*_ {*t* : *P*(*N*_*R*_(*t*) = 0) ≥ 0.90} as a function of *m* in the range *ν*/*K*_+_ < *m* ≲ *m*_*c*_ (all other parameters being kept fixed). τ_90_ is thus the shortest time after which there is at least a 90% chance that resistance has been cleared from the metapopulation. Similarly, we have also determined *τ*_95_(*m*) ≡ min_*t*_{*t* : *P*(*N*_*R*_(*t*) = 0) ≥ 0.95} giving the minimal time for which the *R* clearance probability exceeds 0.95. Therefore, *τ*_90_ and *τ*_95_ give respectively the 90% and 95% percentile of *R* eradication times (see S1 Appendix Sec. 3.2). The results of Fig 4F show that *τ*(*m*) has a single minimum value at essentially the same migration parameter *m* = *m*^∗^ *≈* 3 · 10^−4^ for both 90% and 95% percentiles. Since this is generally the case for strong enough bottlenecks, to shorten the notation and unless specified otherwise, we henceforth refer to *τ*_90*/*95_ simply as *τ* . In the example of Fig 4F, we find *t*^∗^ *τ*(*m*^∗^) ≈ 240 − 270, and *τ* increases sharply when *m* > *m*_*c*_ while *τ*(*m* = 0) > *t*^∗^ (not shown in Fig 4F, we have verified that *τ*(*m* = 0) > 500). Hence, the fluctuation-driven eradication of *R* is most efficient for *m* = *m*^∗^ ≈ 3 · 10^−4^ ∈ [*ν*/*K*_+_, *m*_*c*_], when the probability of resistance clearance after *t* ≈ *t*^∗^ = *τ*(*m*^∗^) microbial generations is close to 1 (see τ_90*/*95_ vs. *m* in Fig 4F). Since the eradication of *R* is here driven by the strong bottlenecks at an average frequency *ν*(1 − δ^2^)/2 ∼ *s* (see “Background” in Model & Methods), *t*^∗^ scales as 1/*s*, i.e. *t*^∗^ = 𝒪 (1/*s*). We have verified that these findings are robust since similar results are obtained for other percentiles and values of *K*_+_/*K*_−_.

Together with the necessary requirements of (8), we thus obtain the following *near-optimal conditions* for the quasi-certain fluctuation-driven eradication of resistance from the metapopulation (for *K*_+_/*K*_−_ fixed):

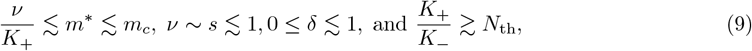

with *P*(*N*_*R*_(*t*^∗^) = 0) ≥ 0.95 after *t*^∗^ = *τ*(*m*^∗^) =𝒪 (1/*s*). Under these conditions, the probability of eradicating resistant cells from the metapopulation after *t* ≈ *t*^∗^ is *near optimal*: For a migration rate *m*^∗^ (with the other parameters fixed and satisfying the conditions (9)), the probability of *R* eradication reaches a set value close to one (Fig 4F), with the fraction *ρ*_*S*_ of *R*-free demes across the metapopulation thus approaching 1 in a time *t*^∗^ =𝒪 (1/*s*) (see S1 Appendix Fig S5H). Remarkably, this means that, under the near-optimal conditions of (9), *slow migration enhances the eradication of resistance compared to the non-spatial case* (*m* = 0, Model & Methods). (Moreover, when *K*_+_/*K*_−_ ≳ *N*_th_*L*^2^, the near-optimal conditions of (9) extend to all *m*, see *Derivation of the critical migration rate m*_*c*_ above). The condition *ν*/*K*_+_ ≲ *m* ≲ *m*_*c*_ is shown as the region within the golden and green lines in Fig 4D-E and corresponds to the near-optimal values *m* ∼ 10^−4^ − 10^−3^ found in Fig 4F. Interestingly, this range of migration rates are of the same order as those studied in Refs. [29, 35], and are consistent with typical microfluidic experiments [107, 126] (see *Translation to the laboratory* in the Discussion subsection “Future directions”). It is worth noting that the conditions of (8) and (9) are essentially independent of the spatial dimension of the metapopulation and hence the fluctuation-driven eradication of resistance is a phenomenon expected to hold on lattices of any dimension; see *Impact of the spatial dimension* in the Discussion subsection “Robustness of results across parameters and scenarios”, and S1 Appendix Sec. 5.5 and Fig S9.

We have also studied the probability of *R* eradication *P*(*N*_*R*_(*t*) = 0) as a function of *K*_+_/*K*_−_ and *m* for a range of slow, intermediate, and fast switching rates *ν* and different values of switching bias *δ*, confirming that *R* eradication occurs chiefly for 0.1 ≲ *ν* ≲ 1 (S1 Appendix Sec. 5.6 Fig S10).

These results demonstrate that not only the fluctuation-driven eradication of the resistant strain *R* arise in the two-dimensional metapopulation under the conditions (8), but that slow migration (*ν*/*K*_+_ ≲ *m* ≲ *m*_*c*_) actually *speeds up* the clearance of resistance and it *can even enhance* the probability of *R* elimination (Discussion and S2 Movie and S1 Appendix Sec. 4), with the best conditions for the fluctuation-driven eradication of *R* given by the relations (9), and corresponding to the clearance of resistance from the grid being almost certain in a near-optimal time *t*^∗^ ∼ 𝒪 (1/*s*).

An intuitive explanation for why slow migration can promote the fluctuation-driven eradication of resistance is illustrated by S1 Appendix Sec. 5.1 Figs S4 and S5: in the absence of migration, when *R* cells randomly take over a deme during periods in the harsh environment, with the low carrying capacity *K* = *K*_−_ (blue in S1 Appendix Fig S5A-D; see also S1 Movie and S1 Appendix Sec. 4), resistance cannot be eradicated from that deme in isolation. However, slow migration allows for sensitive cells to recolonise that deme, from which it is then possible to clear resistance by means of the above fluctuation-driven eradication mechanism (S1 Appendix Figs S4C and S5E-H).

## Discussion

Microbial communities generally live in time-fluctuating environments endowed with spatial structure. Migration in space, environmental variability, and fluctuations thus affect the eco-evolutionary dynamics of bacterial populations [127]. They are particularly relevant to determine the likelihood that cells resistant to antimicrobial drugs survive AMR treatments or thrive in drug-polluted environments [10, 40, 41, 42, 43]. Here, inspired by chemostat and microfluidic setups [25, 88, 93, 107, 126, 25, 94], we have shed further light on cooperative antimicrobial resistance embedded on surfaces in natural environments by investigating a metapopulation model of sensitive (*S*) and cooperative resistant (*R*) cells on a (2D) grid of demes (a one-dimensional metapopulation is considered in S1 Appendix Sec. 5.5, see Fig S9), connected through local cell migration, and subject to a constant drug input rate as well as to time-fluctuating conditions (Model & Methods and “Robustness of results across parameters and scenarios”, below).

In Ref. [26], it was shown that strong population bottlenecks, arising when the environment and deme composition vary on the same timescale (*ν* ∼*s*), can cause fluctuations leading to *R* eradication in isolated demes (no migration), see “Background” in Model & Methods and Fig 1A. This fluctuation-driven *R* eradication mechanism occurs in a biologically relevant regime in well-mixed populations [119, 94, 26]. However, it is not obvious whether, and in what form, this phenomenon still appears in the presence of spatial migration (see *Population size and microbial parameter values*, below). In fact, when *R* and *S* cells migrate at a fast rate, their long-lived coexistence is enhanced [96, 39], thereby promoting the persistence of resistance (S1 Appendix Sec. 2, Figs S2B-C, S3, and S4D, S5I–L).

To address this, we have first determined the critical migration rate *m*_*c*_, above which fluctuation-driven eradication of resistance across the metapopulation becomes unlikely (Eq. (6)). This yields the conditions (8) that ensure eradication of *R* from the metapopulation. Biologically, this occurs when environmental bottlenecks are sufficiently strong (i.e. *K*_+_/*K*_−_ is large enough) to counteract the homogenizing effect of migration. Under these conditions, resistance is cleared from local populations at a higher rate than they are recolonised by *R* cells (see Results and Figs 1B, 2, 3, and 4).

We have also found the near-optimal environmental conditions (9) ensuring a quasi-certain clearance of resistance in the shortest possible time. This has allowed us to show that fluctuation-driven eradication of *R* is fastest under slow-but-nonzero migration, when it is most likely to occur on the relaxation timescale of microbial dynamics (*t*^∗^ = 𝒪 (1/*s*); see Fig 4E and “Slow migration” in Results). Biologically, slow migration allows *S* cells to recolonise *R*-only demes, and eventually to clear resistance from these sites (S1 Appendix Sec. 5.1 Figs S4C and S5E-H, S1-S2 Movies). We note that this slow migration regime is relevant in laboratory chemostat and microfluidic experiments [107, 126], as well as in theoretical studies [29, 35] (see *Population size and microbial parameter values* below). Interestingly, previous works reported that a similar regime of cell migration is optimal for microbial survival in growth-dilution cycles [14]. As explained below, we find that our findings in fluctuating environments are consistent with those earlier results (see “Concordance of results with existing literature”).

In the next subsections we review the assumptions and limitations of our study, discuss how it advances the field in light of the existing literature, address its biological relevance for laboratory experiments, and we also outline possible model extensions.

### Robustness of results across parameters and scenarios

#### Population size and microbial parameter values

We have carried out extensive stochastic simulations of the ensuing metapopulation dynamics, and repeatedly tracked the simultaneous temporal evolution of up to a total of *K*_+_*L*^2^ = 10^7^ microbes distributed across *L*^2^ = 400 spatial demes through hundreds of realisations and thousands of different combinations of environmental parameters and migration rates (Table 1 and S1 Appendix Sec. 3). This is a rather large metapopulation model, even though many experiments are carried out with even bigger microbial populations [41, 128, 129]. In addition, some *in-vivo* host-associated metapopulations are naturally fragmented into a limited number of small demes, e.g. *L*^2^ ≈ 25 and *K* ≈ 1000 in mouse lymph nodes [130, 131, 22, 79] and *L* ≈ 300 and *K* ≈ 100 in mouse intestine crypts [132], resulting in system sizes comparable to those considered here. Our results have been obtained by neglecting the occurrence of mutations, which is acceptable since *R* fluctuation-driven eradication typically occurs on a faster timescale (see Introduction and conditions (9)) compared to mutations [133, 134, 22]. We also note that the resistance mutation rate has been shown to decrease with the population density [133, 134], suggesting that our results can hold in large bacterial populations. It is worth stressing that, according to the conditions of (8), fluctuation-driven eradication of *R* is expected whenever *K*_+_/*K*_−_ ≳ *N*_th_ (and *m* ≲ *m*_*c*_), regardless of the population size and spatial dimension of the metapopulation (see *Impact of the spatial dimension* below, and S1 Appendix Sec. 5.5, Fig S9). Remarkably, this condition is satisfied by values characterising realistic microbial communities, e.g. (*K*_+_, *K*_−_, *N*_th_) ∼ (10^11^, 10^6^, 10^5^) [26] (S1 Appendix Sec. 1.2.3). Furthermore, we note that the indicative values used in our examples for the extra metabolic cost of resistance (*s* = 0.1) and the biostatic impact of the antimicrobial drug (*a* = 0.25), are biologically plausible parameter values, with similar figures used in existing studies [135, 136, 26]. The values of *s* and *a* would typically decrease on a long evolutionary timescale due to compensatory mutations (typically after more than ∼10^3^ microbial generations in a low mutation regime [22]); however, they can be considered to remain constant on the shorter timescale considered here (a few hundred microbial generations). Additionally, results are robust against changes in values of *s* and *a* as long as 0 < *s* < *a* ≲ 10^−1^ [26] (“Background” in Model & Methods). Here we find mathematically convenient to represent the action of the drug as limiting the growth of *S* cells (bacteriostatic scenario) despite *β*-lactams being typically bactericidal drugs (increasing cell death). This choice is acceptable in the regime of low drug concentration considered here, where bactericidal and bacteriostatic antibiotics have a similar action [137, 138]. We note that higher drug concentration would increase the cooperation threshold *N*_th_ (more resistant cells needed to protect *S* individuals), thus increasing the chances that resistant cells spread and fix in many demes (see S1 Appendix Sec. 2).

We have explored a substantially broad range of values of migration rate *m*, spanning four orders of magnitude (*m* ∈ 10^−5^ − 10^−1^), in addition to the benchmark case of no migration (*m* = 0) corresponding to a metapopulation of isolated demes. (Here, *m* ≲ 10^−5^ is effectively equivalent to *m* = 0; see Fig 2A and Fig 4A-E). Slow to moderate migration, *m* ∈ 10^−5^ − 10^−1^, corresponds, on average, to the local migration of 0.001% to 10% of cells during each microbial generation, ranging from effective deme isolation to a significant mixing via dispersal. This wide range of migration rates is consistent with diverse experimental settings (see *Translation to the laboratory in “Future directions”*, below), from standard laboratory chemostats to microfluidic devices, e.g., [107, 126], as well as with theoretical studies, e.g., [29, 14, 35]. Moreover, in line with general principles [96, 39], we have shown that for a migration rate beyond the critical value *m*_*c*_ (fast migration) dispersal strengthens strain coexistence (Figs 2 and 4, and S1 Appendix Figs S2B-C, S3, S4D, and S5I-L). In our examples, the near-optimal migration rate for the fluctuation-driven eradication of *R* is in the range *m* ∼ 10^−4^ − 10^−3^ (slow migration; see below), corresponding on average to one migrant per ∼10^3^ − 10^4^ cells every generation. Overall, this study spans a broad range of migration rates, from isolated sites to fully connected demes, and identifies the parameter regimes under which fluctuation-driven eradication of resistance occurs (see “Critical migration rate” in Results).

#### Environmental assumptions

In biology and ecology, the carrying capacity provides a coarse-grained description of environmental limitations on population growth, arising from diverse factors such as nutrient availability, toxin accumulation, or other environmental conditions [139]. Since these factors fluctuate over time and space, it is natural to assume that the carrying capacity itself varies with the environment [44, 140, 141]. Here, we have thus modelled environmental variability by letting the carrying capacity *K*(*t*) change suddenly and homogeneously across the metapopulation by taking very different values when the environmental conditions are mild or harsh. For the sake of simplicity and concreteness, we have assumed that each deme is subject to the randomly switching binary carrying capacity, *K*(*t*), given by Eq. (3) (see “Environmental variability” in Model & Methods). This binary choice is a convenient way to represent environmental variability in random cycles of feast and famine [51, 52, 53, 78, 55, 54, 26, 28, 27, 79] that we can interpret here as time variations of a spatially homogeneous influx of nutrients [26, 27, 28, 51, 55] (or sequential changes in the antibiotic influx [95, 55]). Notably, it allows us to easily model the drastic population bottlenecks often experienced by microbial communities, whose role is central to this study and important in shaping microbial dynamics [80, 55, 81, 41, 82, 83, 84, 50, 85, 86, 87, 88, 89, 90, 91, 92]. This theoretical simplification is relatively close to laboratory-controlled conditions used in chemostat and microfluidic experiments ([25, 93, 94]). While other choices are possible, such as continuously varying *K*(*t*), these would introduce significant theoretical and computational challenges ([54]) and would be less suitable to capture the sharp bottlenecks that are key for our analysis. (See below for the case of periodic switching of *K*(*t*)).

#### Forms of migration

There are different ways of modelling cells’ dispersal and migration in microbial populations. Cellular movement is often directed towards areas that are rich in resources [113], but dispersal is commonly assumed to happen with a constant per capita migration rate (see, e.g., Refs. [18, 20, 79]). Inspired by directed cell motion, we have first considered a density-dependent form of dispersal, see Eq. (2a), positing that cells from demes whose occupancy is close to the carrying capacity (*N* ≈ *K*, lack of resources) have a higher rate of migration than residents from a lowly populated sites (*N* < *K*, abundance of resources). We have also considered the simpler form of dispersal where all cells can migrate onto a neighbouring deme with a constant per-capita rate *m*, see Eq. (2b). For both types of migration, we have obtained similar results regarding the influence of *m* and *K*_+_/*K*_−_ on the fluctuation-driven eradication of resistance (S1 Appendix Secs. 5.2-5.3 and Figs S6-S7, S4-S5 Movies). These additional data demonstrate the robustness of our findings that are qualitatively independent of the specific choice of dispersal considered here. Extending this work to species-specific, directional, or spatially dependent migration rates would be particularly relevant for more complex metapopulation structures [15, 20].

#### Impact of the spatial dimension and accuracy of the critical migration prediction

In this study, for the sake of concreteness, we have focused on a metapopulation model consisting of a two-dimensional (2D) grid of *L* × *L* demes connected by cell migration (Fig 1B; Model & Methods). This provides a natural framework for modelling microbial communities inhabiting surfaces where cellular migration occurs, a setting commonly used in both theoretical and experimental studies [75, 76, 31, 77]. Possible applications include the human skin [6], the digestive tract [7], plant leaf surfaces [4], the seabed [2], and other wet environments [3]. While the results presented in Figs 2–4 were obtained for the 2D model, we have also analysed a one-dimensional (1D) metapopulation consisting of a ring of demes, or cycle, in S1 Appendix Sec. 5.5 Fig S9. These results show that our predictions also hold qualitatively in 1D lattices. The main difference between Fig 4 and S1 Appendix Fig S9 is that, in the latter, eradication of *R* occurs for values of *m* up to ten times larger than in the former, which is in agreement with Eq. (6). The fact that the theoretical prediction for *m*_*c*_ (Eq. (6)) captures the critical migration rate quantitatively in 2D (Figs 2A, 4A–E; S1 Appendix Figs S6A, S7A, S10) but only qualitatively in 1D stems from Eq. (6) being a mean-field result, independent of spatial dimension. This expression neglects deme-to-deme spatial correlations, which are particularly relevant in low dimensions (“Critical migration rate” in Results). Consequently, the approximation of *m*_*c*_ provided by Eq. (6) improves with increasing spatial dimension, and is therefore expected to work even better in three-dimensional metapopulations. Note that the conditions (8) depend on spatial dimension only through the actual critical migration rate *m*_*c*_, of which the expression Eq. (6) is a mean-field approximation. As a result, fluctuation-driven eradication of resistance is expected to occur on metapopulation lattices (regular graphs) in any spatial dimension, provided that the conditions of (8) are satisfied, and to be most efficient under the near-optimal conditions (9).

The effects of spatial structures such as star graphs, island models, and cycles have been investigated for non-cooperative strain competition under slow migration in static environments [18, 22], as well as under time-varying external conditions [79]. Serial dilution experiments have also motivated studies of growth-and-dilution cycles coupled on graphs under fast migration [15, 20]. Understanding the impact of complex spatial structures on cooperative antimicrobial resistance, however, remains largely an open problem.

#### Extra vs intracellular drug inactivation

Our theoretical model of cooperative antimicrobial resistance captures the inactivation of the drug by *R* cells, either through extra or intracellular resistance enzymes (see “Metapopulation model” in Model & Methods). In the latter case, more *R* cells are required to protect *S* from drug exposure, which could be represented by a higher cooperation threshold *N*_th_ than in the extracellular case, consistently with Ref. [142]. Moreover, in this metapopulation setting we assume that the public good (shared drug protection) does not directly spread to neighbouring demes, because the drug degradation process takes place locally on a timescale much shorter than that of microbial replication (e.g., a single *β*- lactamase enzyme can hydrolyse up to ∼10^3^ antibiotic molecules per second [143]). This assumption is consistent with each deme receiving a parallel inflow of medium (including antibiotics), as in typical microfluidic setups [107, 93] and in parallel chemostats (see *Translation to the laboratory* in “Future directions”, below). However, since the active drug and resistant enzyme concentration in this study is set by the local number of *R* cells (see “Metapopulation model” in Model & Methods), their dispersal across demes can also be interpreted as an effective form of drug and public good diffusion through the metapopulation. Similarly, the diffusion of available resources across demes is indirectly captured by the density-dependent migration transition rate of Eq. (2a), in which resource-consuming individuals tend to disperse away from demes with low resource availability (i.e., when *N*/*K* is high; see “Intra- and inter-deme processes” in Model & Methods).

### Concordance of results with existing literature

On the one hand, most related studies that investigate the impact of fluctuating environments have focused on two-strain competition dynamics in well-mixed communities, either in the absence of public goods or when cooperative behaviour benefits both strains [51, 52, 53, 54, 55, 144], including the case of time-varying nutrient and toxin concentrations [27]. The eco-evolutionary dynamics of cooperative AMR in well-mixed populations, under binary time-varying environmental conditions, has been studied in Refs. [26, 28], that are directly relevant for this work and revealed when fluctuations can lead to the eradication of resistant cells.

On the other hand, the impact of spatial structure—such as star graphs and island models—has been typically studied in static environments under slow migration [18], with non-cooperative AMR considered in Ref. [22] and Ref. [15] experimentally investigating the spread of an antibiotic-resistant mutant through a star graph. Other studies have examined the dynamics of bacterial colonies of resistant and sensitive cells undergoing range expansion in constant environments [142, 145, 146]. Remarkably, recent metapopulation studies have investigated growth-and-dilution cycles coupled to fast cell migration [20], and competing wild-type and mutant cells subject to feast-and-famine cycles on metapopulation lattices [79]. However, metapopulation studies that include some form of environmental stochasticity have generally not addressed cooperative resistance. Instead, they typically focus on single-strain populations, island models, global migration, or spatially heterogeneous metapopulations [58, 59, 60, 61, 62, 63].

Moreover, a substantial body of literature has focused on *rescue dynamics*, chiefly on *demographic rescue* in structured metapopulations [57], which occurs when a declining population is rescued from local extinction by an influx of individuals migrating from neighbouring demes – for instance, by restoring resistant cells in *R*-free demes after strong bottlenecks (Fig 3). Interestingly, Ref. [14] reported that *intermediate* migration rates (similar to our slow/moderate regime) maximise species persistence time in paired batch cultures undergoing growth–migration–dilution cycles. This results from recolonization events following local extinctions and is aligned with findings from other computational studies of non-cooperative rescue dynamics [65, 66, 67, 68] and with experimental observations [69, 70, 71, 72, 73]. Here, we have found that slow-but-nonzero migration rates – of similar magnitude to those in Ref. [14] – enhance the *fluctuation-driven eradication of resistance* (Fig 4 and S1 Appendix Sec. 5.1 Figs S4 and S5), rather than rescuing resistance. These results are however compatible with those of Ref. [14]. As discussed in “Slow migration” (Results), here slow migration promotes *S*-cell recolonisation of *R*-only demes – analogous to the rescue dynamics of Ref. [14] – but subsequently renders these demes prone to fluctuation-driven eradication of resistance when the environment switches from mild to harsh conditions (S1 Appendix Sec. 5.1 Figs S4 and S5). Note that the *fluctuation-driven eradication* mechanism does not arise in Ref. [14], because the latter involves two mutually cooperative strains resistant to two drugs, and considers growth–dilution cycles rather than the binary environmental switching studied here.

The *fluctuation-driven eradication* mechanism was unveiled in Ref. [26]. While it relies on biologically realistic assumptions (see above), some of these do not correspond to commonly used laboratory setups, which explains why this phenomenon has not yet been tested experimentally. Specifically, most experiments with environmental bottlenecks involve serial dilution protocols, where each cycle consists of a period of exponential growth followed by an instantaneous dilution step [15, 147] (see *Translation to the laboratory* in “Future directions”, below). Such systems generally do not exhibit fluctuation-driven eradication of *R*, since this phenomenon typically requires the population to spend finite periods in the harsh environment (see “Background” in Model & Methods and [26]). This also applies to studies of microbial cooperation [56], including those investigating the effects of “dilution shocks” [13] or “disturbance events” [87], as well as cooperative antimicrobial resistance [14]. Moreover, as we discuss below in *Translation to the laboratory*, there is currently no clear correspondence between chemostat-inspired models and those based on serial dilution experiments. Although recent chemostat experiments have analysed the effects of environmental fluctuations in well-mixed populations switching between different nutrient sources [25], these did not consider cooperative resistance and therefore did not display fluctuation-driven eradication. Similar environmental fluctuations to those considered here have also been implemented in metapopulation microfluidic experiments, e.g., in Ref. [93] for phenotypic switching in a single strain.

Remarkably, the authors of Ref. [20] investigated a spatially structured (non-cooperative) two-strain model consisting of subpopulations connected by migration and subject to bottlenecks arising from growth–dilution cycles. They found that slow migration can amplify selection for the fittest strain, whereas fast migration tends to suppress selection. However, the lack of cooperation and the serial dilution dynamics of the system considered in Ref. [20] (inspired by batch culture setups) cannot give rise to fluctuation-driven eradication. It is worth noting that the authors of Ref. [22] studied resistance rescue in a metapopulation of sensitive cells and non-cooperative drug-resistant mutants, showing that spatial structure can facilitate the survival of resistance. Since the model of Ref. [22] is non-cooperative and subject to a single environmental change (when a biostatic drug is added), it does not exhibit fluctuation-driven resistance eradication.

Cooperative resistance in spatially structured metapopulations has also been studied experimentally, for instance in the range expansion experiments of Refs. [142, 145, 146], which were performed under constant environmental conditions. As discussed above, these are conditions under which fluctuation-driven resistance eradication cannot occur (S1 Appendix Sec. 2, Fig S2). Moreover, as detailed above, Ref. [14] investigated cooperative AMR in a spatial setup where no fluctuation-driven eradication is expected. We also note that Ref. [21] focuses on cooperative resistance rescue under environmental fragmentation into random, independent subpopulations (without migration). While this model cannot exhibit fluctuation-driven eradication (as it involves a single strain), the authors of Ref. [21] show that habitat fragmentation enhances resistance rescue.

To the best of our knowledge, no previous study has examined the optimal conditions for eradicating cooperative drug-resistant cells in a stochastic metapopulation composed of sensitive and resistant strains, where demes are connected by local migration and subject to global feast–famine cycles. Here, we present the first metapopulation study showing that slow-but-nonzero migration helps eradicate cooperative AMR in time-fluctuating environments. This is in stark contrast – yet fully consistent (see above) – with previous results showing that slow migration, in environmental conditions different from those considered here, helps maintain cooperative AMR [14] and non-cooperative strain competition [65, 66, 67, 68, 69, 70, 71, 72, 73].

### Future directions

#### Translation to the laboratory: chemostats, microfluidic setups, and batch cultures

Our modelling approach is mainly inspired by chemostat setups, which are commonly used in laboratory-controlled experiments to modulate the influx of nutrients and drugs in microbial communities. In such systems, the concentrations of resources and toxins can be adjusted to impose harsh conditions that generate population bottlenecks, whose eco-evolutionary impacts are the subject of intense study [25, 88, 93]. Here, we focus on the biologically relevant regime of intermediate environmental time variation [25, 119, 94], characterised by *ν* ≲ 1 and 0 ≤ *δ* ≲ 1, in which the population size within each deme rapidly tracks the carrying capacity, whereas the local composition (number of *S* and *R* cells in a deme) relaxes more slowly, on a timescale ∼ 1/*s* with typically *s* ≲ 10^−1^ (*s* = 0.1 in all figures here; see “Background” in Model & Methods and S1 Appendix Sec. 1.2.1). This regime corresponds to conditions fluctuating between mild (*K* = *K*_+_) and harsh (*K* = *K*_−_) environmental states with a frequency between once per hour and once per day, that is, approximately every 1 − 100 microbial generations (*ν* = 0.01 − 1; see Model & Methods). The drug influx is kept constant, and each environmental switch, theoretically treated as instantaneous, occurs rapidly in practice. While we have conveniently represented the switching of the carrying capacity as a random process at rates *ν*_*±*_ (Fig 1A), the case where *K* varies periodically between *K*_+_ and *K*_−_, with period 1/*ν*_+_ + 1/*ν*_−_, would not change the qualitative results of our study [53] (“Background” in Model & Methods, S1 Appendix Sec. 1.2.3), and could be seen as a potential laboratory implementation of this model.

Since all the above conditions can be practically implemented [25, 94], we believe that our theoretical predictions can, in principle, be probed in prospective laboratory-controlled experiments. The environmental switching of *K*(*t*) would be realised using a sequence of spatially connected, fixed-volume chemostats, each acting as a deme, see, e.g., Ref. [148]. The rate of cell migration would be set by the rate of volume exchange between neighbouring demes-chemostats (0.001% − 10% of the volume every hour). Moreover, with microfluidic devices and single-cell techniques, it is possible to perform spatially structured experiments involving as few as 10 − 100 cells per microhabitat patch [9, 107, 149]. These conditions are consistent with our modelling parameters, notably those corresponding to demes of relatively small size under harsh conditions (e.g. *K* = *K*_−_ = 80). The migration rate between microhabitats in such setups largely depends on the experimental design (e.g. number of patch-to-patch channels, channel cross-section).

It is worth noting that most laboratory experiments are performed with batch cultures, which are characterised by cycles of exponential growth followed by instantaneous dilution steps; see, e.g., Refs. [15, 147]. These setups are generally easier to operate than chemostat or microfluidic systems. However, in batch cultures, the concentrations of nutrients (microbial consumption) and drugs (enzymatic degradation by *R* cells) vary continuously over time, making their theoretical modelling particularly challenging [20, 150]. Establishing a neat correspondence between theoretical models inspired by chemostats and serial dilution cycles therefore remains largely an open problem. The gradual degradation of the drug can play a critical role in the eco-evolutionary dynamics of cooperative AMR, as shown in Ref. [21], where the metapopulation fragmentation into isolated demes enhances the maintenance of resistance. See also, e.g., the 2D experimental study of Ref. [31], and the theoretical works such as Refs. [29, 35], which show how spatial heterogeneity in drug concentration shapes the spatio-temporal dispersal of cells and resistance.

#### Beyond two strains and cooperative resistance

We have focused on a two-strain metapopulation model, but our analysis can be readily extended to cases involving multiple sensitive strains and a single resistant type. For this extension, the fraction of *R* cells in each deme should fluctuate around a low but non-zero value (here, *N*_th_/*K*_+_ ≪ 1; see Fig 1). Moreover, spatial fluctuation-driven eradication of resistance requires that the number of cells in each deme sharply decreases following a population bottleneck, while the deme composition evolves on a slower timescale. This leads to a small *R* subpopulation in each deme that is prone to extinction. In our model, the number of resistant cells per deme after a bottleneck is approximately *N*_th_ *K*_−_/*K*_+_ ≲ 1 (Fig 1; see “Background” in Model & Methods).

In contrast, in the case of non-cooperative antimicrobial resistance, the resistant strain does not share its protection with sensitive cells. Thus, when, as here, the metapopulation is subject to a steady drug influx, the spread of sensitive cells is hindered by the presence of the drug, and their fitness remains lower than that of resistant cells. In this case, no fluctuation-driven eradication occurs, since the fraction of resistant cells typically outgrows that of sensitive ones. For non-cooperative resistance, if the initial number of *R* cells is sufficiently large, resistance is expected to eventually take over the entire metapopulation [18, 20, 22, 151].

## Conclusions

Environmental variability, spatial structure, cellular migration, and demographic fluctuations are ubiquitous and key factors influencing the temporal evolution of cooperative antimicrobial resistance. The combined effects of dispersal and fluctuations in structured environments are complex and pose numerous challenges [96, 127, 123]. While dispersal generally promotes strain coexistence in spatially organised populations [39], we show that, counterintuitively, migration can instead strongly facilitate the elimination of one strain in temporally fluctuating environments.

In this study, we have demonstrated – by theoretical analysis and computational simulations – that environmental variability and, critically, slow-but-nonzero migration can lead to the efficient eradication of cooperative drug resistance in one- and two-dimensional lattice metapopulations consisting of drug-resistant and sensitive cells. We have identified the conditions under which fluctuation-driven eradication of the resistant strain occurs in general metapopulation lattices, and the near-optimal parameter regimes for this mechanism to operate in the shortest possible time. Our main findings, obtained explicitly for two-dimensional and cycle metapopulations, remain qualitatively valid for lattices of any spatial dimension.

This work demonstrates that the interplay between environmental variability, demographic fluctuations, and slow migration can enable the efficient eradication of cooperative antimicrobial resistance from an entire metapopulation through fluctuation-induced bottlenecks. This mechanism is effective when environmental changes occur on timescales comparable to microbial dynamics, providing a plausible strategy for laboratory or therapeutic interventions to eradicate resistance that would otherwise persist under static conditions. We believe that our theoretical predictions, which are robust to model variations, are relevant to realistic microbial populations and could, in principle, be tested experimentally using chemostat setups and/or microfluidic devices [25, 88, 93]. More broadly, our work illustrates how environmental fluctuations can be harnessed to achieve desired evolutionary outcomes, such as eliminating antibiotic resistance. We hope that these theoretical results will motivate further experimental investigations of environmental variability in chemostat and microfluidic systems, and inform the development of novel clinical protocols and therapeutic strategies aimed at preventing the spread of antimicrobial resistance.

## Data availability

The data generated and used within this work can be found at the Open Science Framework repository (Lluís Hernández-Navarro, Kenneth Distefano, Uwe C. Täuber, and Mauro Mobilia. 2024. Supplementary data, code, and movies for “Slow spatial migration can help eradicate cooperative antimicrobial resistance in time-varying environments”. OSF. https://doi.org/10.17605/OSF.IO/EPB28).

## Code availability

The C++ code used to generate the data and the Python and Matlab codes to process and visualize the data within this work can be found at the Open Science Framework repository (Lluís Hernández-Navarro, Kenneth Distefano, Uwe C. Täuber, and Mauro Mobilia. 2024. Supplementary data, code, and movies for “Slow spatial migration can help eradicate cooperative antimicrobial resistance in time-varying environments”. OSF. https://doi.org/10.17605/OSF.IO/EPB28).

## Acknowledgements

The authors would like to thank M. Asker, J. Jiménez, S. Muñoz Montero, M. Pleimling, A. M. Rucklidge, and M. Swailem for fruitful discussions. L. H. N. and M. M. gratefully acknowledge funding from the U.K. Engineering and Physical Sciences Research Council (EPSRC) under the Grant No. EP/V014439/1 for the project ‘DMS-EPSRC Eco-Evolutionary Dynamics of Fluctuating Populations’ (https://eedfp.com/). K. D. and U. C. T.’s contribution to this research was supported by the U.S. National Science Foundation, Division of Mathematical Sciences under Award No. NSF DMS-2128587.

## Author contributions

**Conceptualization:** Lluís Hernández-Navarro, Mauro Mobilia.

**Data curation:** Kenneth Distefano, Lluís Hernández-Navarro.

**Formal analysis:** Lluís Hernández-Navarro, Kenneth Distefano, Mauro Mobilia.

**Funding acquisition:** Mauro Mobilia, Uwe C. Täuber.

**Investigation:** Lluís Hernández-Navarro, Kenneth Distefano.

**Methodology:** Lluís Hernández-Navarro, Kenneth Distefano, Mauro Mobilia.

**Project administration:** Mauro Mobilia.

**Resources:** Uwe C. Täuber, Mauro Mobilia.

**Software:** Kenneth Distefano.

**Supervision:** Mauro Mobilia, Uwe C. Täuber.

**Validation:** Kenneth Distefano, Lluís Hernández-Navarro.

**Visualization:** Lluís Hernández-Navarro, Kenneth Distefano, Mauro Mobilia.

**Writing – original draft:** Lluís Hernández-Navarro, Mauro Mobilia, Kenneth Distefano.

**Writing – review & editing:** Lluís Hernández-Navarro, Mauro Mobilia, Kenneth Distefano, Uwe C. Täuber.

## Supporting information

**S1 Appendix**. The Supplementary Appendix comprises further technical details about the model and methods, some additional results, 10 supplementary figures, and a description of how the stochastic simulations were performed. It also contains a description of the Supplementary Movies. The Supplementary Appendix is also electronically available from the *bioRxiv* repository at https://www.biorxiv.org/content/10.1101/2024.12.30.630406v3.supplementary-material.

The Supplementary Movies are also electronically available from the *OSF repository* at https://doi.org/10.17605/OSF.IO/EPB28.

**S1 Movie**. Resistant cells can survive in switching environments when demes are fully isolated. Full description in S1 Appendix.

**S2 Movie**. Slow migration can enhance the fluctuation-driven eradication mechanism. Full description in S1 Appendix.

**S3 Movie**. Fluctuation-driven eradication of *R*. Full description in S1 Appendix.

**S4 Movie**. Fluctuation-driven eradication mechanism with density-dependent migration. Full description in S1 Appendix.

**S5 Movie**. Fluctuation-driven eradication mechanism with density-independent migration. Full description in S1 Appendix.

