## Supplementary material for "Slow spatial migration can help eradicate cooperative antimicrobial resistance in time-varying environments": S1 Appendix

February 17, 2026

In this Supplementary Information, we provide additional details and supporting material to the manuscript "Slow spatial migration can help eradicate cooperative antimicrobial resistance in time-varying environments". This information is electronically available alongside the preprint of the manuscript at the *bioRxiv* repository (https://doi.org/10.1101/2024.12.30.630406). Throughout this document, Eq. ($m$) and Fig $n$ refer, respectively, to Equation ($m$) and Figure $n$ of the manuscript.

### 1 Additional details of the model & methods

In this section, we provide additional details on the model by discussing the master equation encoding its individual-based dynamics, and giving further details of the eco-evolutionary dynamics in a single isolated deme.

#### 1.1 Master equation of the two-dimensional metapopulation model

As discussed in the main text (Model & Methods), the full metapopulation model is a continuous-time multivariate Markov process, and its dynamics with environmental fluctuations is characterised by the probability $P(\{N_R, N_S\}, \xi|t)$ that its microbial population at time $t$ in any given deme, denoted by a two-dimensional position vector $\vec{u}$, consists of $N_R(\vec{u})$ and $N_S(\vec{u})$ resistant and sensitive cells, in the environmental state $\xi(t) = \pm 1$, with $K(\xi) = K_\pm$ when $\xi = \pm 1$. The metapopulation make-up is encoded in $\{N_R, N_S\} \equiv \{N_R(\vec{u}), N_R(\vec{u'}), N_R(\vec{u''}), ..., N_S(\vec{u}), N_S(\vec{u'}), N_S(\vec{u''}), ...\}$, where the vectors $\vec{u}, \vec{u'}, \vec{u''}, ...$ denote each of the $L^2$ demes. Given the migration-mediated interactions between nearest-neighbour demes in the $L \times L$ grid (with periodic boundary conditions), the master equation characterising the stochastic time-evolution of the metapopulation reads [1]:

$$\begin{aligned}\frac{\partial P}{\partial t}=\sum_{\vec{u}}\Bigg\{&\left(\mathbb{E}_R^-(\vec{u})-1\right)T_R^+(\vec{u})P+\left(\mathbb{E}_R^+(\vec{u})-1\right)T_R^-(\vec{u})P+\left(\mathbb{E}_S^-(\vec{u})-1\right)T_S^+(\vec{u})P+\left(\mathbb{E}_S^+(\vec{u})-1\right)T_S^-(\vec{u})P\\&+\frac{1}{2}\sum_{\vec{u}'\ \text{n.n.}\ \vec{u}}\Bigg[\left(\mathbb{E}_R^+(\vec{u}')\mathbb{E}_R^-(\vec{u})-1\right)T_R^{M_i}\left(\vec{u}'\to\vec{u}\right)P+\left(\mathbb{E}_R^+(\vec{u})\mathbb{E}_R^-(\vec{u}')-1\right)T_R^{M_i}\left(\vec{u}\to\vec{u}'\right)P\\&\qquad+\left(\mathbb{E}_S^+(\vec{u}')\mathbb{E}_S^-(\vec{u})-1\right)T_S^{M_i}\left(\vec{u}'\to\vec{u}\right)P+\left(\mathbb{E}_S^+(\vec{u})\mathbb{E}_S^-(\vec{u}')-1\right)T_S^{M_i}\left(\vec{u}\to\vec{u}'\right)P\Bigg]\Bigg\}\\&+\nu_{-\xi}P(\{N_R,N_S\},-\xi|t)-\nu_\xi P(\{N_R,N_S\},\xi|t),\end{aligned}\tag{S1.1}$$

where $\mathbb{E}_{R/S}^{\pm}(\vec{u})$ are shift operators that increase $(+)$ or decrease by one $(-)$ the number of $R$ or $S$ cells in deme $\vec{u}$, i.e., they increase or decrease by one the value of $N_R(\vec{u})$ or $N_S(\vec{u})$, which are the components of the set $\{N_R, N_S\}$ that correspond to deme $\vec{u}$, and $i = 1, 2$. To simplify the notation, except in the last line, we dropped all explicit dependencies on $\{N_R, N_S\}$, $\xi$, and $t$. In (S1.1), $\sum_{\vec{u}}$ runs over all $L^2$ demes in the metapopulation, and the sum $\sum_{\vec{u}}\sum_{\vec{u}'\ \text{n.n.}\vec{u}}$ is over the four nearest neighbours (n.n.) $\vec{u}'$ of each deme $\vec{u}$.

The birth $T_{R/S}^+(\vec{u})$ and death $T_{R/S}^-(\vec{u})$ rates in deme $\vec{u}$ are given by Eq. (1). $T_{R/S}^{M_i}(\vec{u}\to\vec{u}')$ are the rate at which an $R/S$ cell migrates from deme $\vec{u}$ to one of its nearest neighbours $\vec{u}'$ given by (2a,b): It is the transition rate $T_{R/S}^{M_1}(\vec{u}\to\vec{u}') = \frac{m}{4}\frac{N(\vec{u}))}{K(t)}N_{R/S}(\vec{u})$ in the case of density-dependent migration $(i = 1)$, and the rate $T_{R/S}^{M_2}(\vec{u}\to\vec{u}') = \frac{m}{4}N_{R/S}(\vec{u}))$ for density-independent migration $(i = 2)$. In the last line of (S1.1) we adopt the notation $\nu_\xi \equiv \nu_\pm$ when $\xi = \pm1$. The first line on the right-hand side of the master equation encodes births and deaths of resistant and sensitive cells in each deme $\vec{u}$; the second and third lines describe the inward and outward migration of $R$ or $S$ microbes, respectively; and the final line accounts for the random environmental switching.

The master equation (S1.1) is specifically given for the two-dimensional lattice considered here, but it can readily be generalized to any regular lattice of $d$ dimensions (with periodic boundary conditions) consisting of $L^d$ connected demes, see, e.g., Ref. [2]. For completeness, the case of a one-dimensional metapopulation of size $L$ with periodic boundary conditions is briefly considered in Sec. 5.5 (see Fig S9).

This multivariate master equation can be simulated efficiently using the stochastic methods described in Section 3. It is worth noting that demographic fluctuations eventually lead to the extinction of the entire metapopulation, but this phenomenon is unobservable as it typically occurs after an enormous time, that grows dramatically with the system size and can generally not be observed in sufficiently large metapopulations as those considered here.

#### 1.2 Further details on the eco-evolutionary dynamics in an isolated deme

As explained in Model & Methods of the main text, it is instructive to review the properties of the eco-evolutionary dynamics in a single isolated deme (with $m = 0$), first by considering the mean-field approximation where all fluctuations are ignored, then the dynamics in a *static environment* with a constant and finite carrying capacity, and then by studying the piecewise-deterministic approximation of the eco-evolutionary dynamics in a time-fluctuating environment. We notice that the dynamics in a single deme is fully described by the master equation when we formally set $L = 1$ and $m = T_{R/S}^{M_i} = 0$.

##### 1.2.1 Eco-evolutionary dynamics in an isolated deme: Mean-field approximation

It is instructive to first ignore fluctuations entirely and consider the mean-field dynamics in an isolated deme $\vec{u}$ where the carrying capacity is assumed to be constant and extremely large, $K = K_0 \to \infty$. The deme is thus unlikely to go extinct, and with the transition rates (1), the mean-field eco-evolutionary dynamics in $\vec{u}$ is characterised by rate equations for the deme size $N = N(\vec{u}, t)$ and the fraction $x = N_R(\vec{u}, t)/N$ of resistant cells in the deme, which read [3, 4]:

$$\begin{aligned}\dot{N} &= T_R^+ - T_R^- + T_S^+ - T_S^- = N\left(1-\frac{N}{K_0}\right),\\ \dot{x} &= \frac{T_R^+ - T_R^-}{N} - x\frac{\dot{N}}{N} = \frac{f_R - f_S}{\bar{f}}x(1-x) = \begin{cases}-\frac{sx(1-x)}{1-sx} & \text{if } x \geq N_{\text{th}}/N,\\ \frac{(a-s)x(1-x)}{1-a+(a-s)x} & \text{if } 0 \leq x < N_{\text{th}}/N,\end{cases}\end{aligned}\tag{S1.2}$$

where the dot indicates the time derivative and we have used the expressions of $T^{\pm}_{R/S}$ given by (1), with $f_R = 1 - s$, $f_S = 1$ when $x \geq N_{\rm th}/N$ and $f_S = 1 - a$ when $x < N_{\rm th}/N$ $(0 < s < a < 1)$, and $\overline{f} = (xf_R + (1-x)f_S)$ (Model & Methods) [5, 6]. The logistic rate equation for $N$ predicts the relaxation of the deme size towards the constant carrying capacity $N \to K_0$ on a timescale $t \sim 1$. Clearly, the fraction of resistant cells $x$ is coupled to the deme size $N$: it decreases on a timescale $t \sim 1/s$ when $x \geq N_{\rm th}/N$, whereas it grows on a timescale $t \sim 1/(a-s)$ when $x < N_{\rm th}/N$. Since $s \lesssim 1$ and $a \lesssim 1$, in this mean-field picture, the fraction $R$ relaxes on a slower timescale $t \sim 1/|f_R - f_S| > 1$, with $N_R = Nx \to N_{\rm th}$, yielding a long-time fraction $x \to N_{\rm th}/K_0$ of $R$ cells in the deme [5, 6]. Similarly, the number of $S$ cells relax towards $N_S = N(1-x) \to K_0(1-x) = K_0 - N_{\rm th}$ on timescale $t \sim 1$ and the fraction of $S$ approaches $1 - N_{\rm th}/K_0$ on the longer timescale $t \sim 1/|f_R - f_S|$. In all our examples, we consider $|f_R - f_S| \sim s \ll 1$, ensuring a clear timescale separation between the dynamics of the deme size and its make-up, with $N$ and $x$ being respectively the fast and slow variables.

##### 1.2.2 Eco-evolutionary dynamics of an isolated deme subject to a static environment

In a static environment where the carrying capacity is large but *finite* and constant, $K = K_0 \gg 1$, the deme is unlikely to go extinct in an observable time and its size fluctuates about the carrying capacity, with $N(\vec{u}) \approx K_0$, see Eq. (3), with the deme composition that changes via the intra-deme cell division and death according to the reactions process given in the main text (see Model & Methods) occurring with the transition rates (1).

In this static environment, the intra-deme dynamics can be aptly approximated by a Moran process for a deme $\vec{u}$ of constant and finite size $N(\vec{u}) = K_0$ [7, 8, 3, 4, 5, 6] and we can describe the intra-deme dynamics by tracking the number of resistant and sensitive cells in $\vec{u}$, respectively simply denoted by $N_R$ and $N_S = N - N_R = K_0 - N_R$. In the realm of the Moran approximation, the deme composition $(N_R, N_S) = (N_R, K_0 - N_R)$ thus changes according to the Moran process [7, 8, 9, 10]

$$\begin{aligned}
(N_R, N_S) &\xrightarrow{\widetilde{T}^+_R} (N_R + 1, N_S - 1), \\
(N_R, N_S) &\xrightarrow{\widetilde{T}^-_R} (N_R - 1, N_S + 1),
\end{aligned} \tag{S1.3}$$

where the Moran transition rates are defined in terms of $T^{\pm}_{R/S}$, given by (1) with $K(t) = K_0$, according to [3, 4, 5, 6, 11, 2]

$$\begin{aligned}
\widetilde{T}^+_R(N_R) &= \frac{T^+_R T^-_S}{K_0} = \frac{f_R}{\overline{f}} \frac{N_R N_S}{K_0} = \frac{f_R}{\overline{f}} N_R \left(1 - \frac{N_R}{K_0}\right), \\
\widetilde{T}^-_R(N_R) &= \frac{T^-_R T^+_S}{K_0} = \frac{f_S}{\overline{f}} \frac{N_R N_S}{K_0} = \frac{f_W}{\overline{f}} N_R \left(1 - \frac{N_R}{K_0}\right).
\end{aligned} \tag{S1.4}$$

These effective transition rates correspond to the increase and decrease in the number of resistant cells in the isolated deme $\vec{u}$ of finite size $K_0$. The Moran process defined by (S1.3) and (S1.4) conserves the deme size $N = K_0$ by accompanying each birth of an $R/S$ by the simultaneous death of an $S/R$ cell, and is characterised by the absorbing states $(N_R, N_S) = (K_0, 0)$ ($R$ fixation) and $(N_R, N_S) = (0, K_0)$ ($S$ fixation). The fixation probability and mean fixation of this Moran process can be computed analytically using classical techniques [8, 12, 9, 10]. When the initial number of $R$ cells in the deme is at equilibrium, i.e. $N_R = N_{\rm th}$ (equilibrium fraction of $N_{\rm th}/K_0$ resistant cells, see above), with $0 < s < a < 1$, it has been shown that the fixation probability of the $R$ strain for this Moran process is [5]

$$\phi(K_0) \simeq \frac{1}{1 + \frac{1}{(1-s)^{K_0 - K_0^*}}}, \tag{S1.5}$$

when $K_0 \gg N_{\rm th} \gg 1$, where

$$K_0^* \equiv N_{\rm th} \frac{\ln(1-a)}{\ln(1-s)} - \frac{\ln[s(1-a)/(a-s)]}{\ln(1-s)} \tag{S1.6}$$

is the deme size for which both strains have the same fixation probability 1/2 [5]. The result (S1.5) shows that $R$ is most likely to fixate the deme ($x \to 1$) if the equilibrium fraction of resistant cells is sufficiently high, namely if $K_0 < K_0^*$ and thus $N_{\rm th}/K_0 \gtrsim \frac{\ln(1-s)}{\ln(1-a)}$ (see (S1.6)), whereas the fixation of $S$ ($x \to 0$) is the most likely outcome when $K_0 > K_0^*$, i.e. if $N_{\rm th}/K_0 \lesssim \frac{\ln(1-s)}{\ln(1-a)}$, see Figs 3 and S1 in Ref. [5]. The expression of the mean fixation time can be found in Ref. [5] where it is found to increase with $K_0$ and $N_{\rm th}$, and become unobservable for large values of these parameters [5, 6]. As a consequence, an isolated deme is doomed to be taken over by resistant cells if the $R$ equilibrium fraction is high enough (greater than $\ln(1-s)/\ln(1-a)$). Otherwise, there is long-lived coexistence of resistant and sensitive cells [5]. In Refs. [5, 6, 11], the coexistence of the strains is considered to be long-lived when it persists for periods exceeding $2K_0$ (twice the typical deme size).

##### 1.2.3 Eco-evolutionary dynamics in an isolated deme subject to a time-fluctuating environment: Piecewise-deterministic approximation and fluctuation-driven eradication of resistance

When deme size is sufficiently large for demographic fluctuations to be negligible at all times, randomness only arises from environmental variability via the time-switching carrying capacity.

In this work, we consider a binary time-varying carrying capacity endlessly switching between values to represent "feast and famine" cycles of alternating mild and harsh conditions [3, 4, 13, 14, 15, 16, 17, 11, 2]. Here, each deme is assumed to have the same time-varying carrying capacity $K(t)$, where

$$K(t) = \frac{1}{2}\left[K_+ + K_- + \xi(t)(K_+ - K_-)\right] \tag{S1.7}$$

is driven by the dichotomous Markov noise (DMN) $\xi(t) \in \{-1, 1\}$ (Eq. (5) in Model & Methods). Therefore, $K(t)$ randomly switches between two possible values, $K = K_+ \gg 1$ in the mild environment and $K = K_- < K_+$ in the harsh environmental state, at rates $\nu_+ = \nu(1-\delta)$ and $\nu_- = \nu(1+\delta)$ according to $K_- \underset{\nu_+}{\overset{\nu_-}{\rightleftharpoons}} K_+$, with a stationary average given by $\langle K(t)\rangle = \left(\frac{1-\delta}{2}\right)K_- + \left(\frac{1+\delta}{2}\right)K_+$; see Model & Methods and Figs 1, S1B,C and S4A.

The size in an isolated deme is in turn driven by the time-switching carrying capacity $K(t)$, and its dynamics is well approximated by the piecewise deterministic Markov process ($N$-PDMP) [18, 3, 4, 13, 15, 16, 17, 11, 5, 6, 2] defined by

$$\dot{N} = N\left(1 - \frac{N}{K(t)}\right) = \begin{cases} N\left(1 - \frac{N}{K_-}\right) & \text{if } \xi = -1, \\ N\left(1 - \frac{N}{K_+}\right) & \text{if } \xi = 1, \end{cases} \tag{S1.8}$$

where we have used (S1.7). In the realm of the the $N$-PDMP approximation, the deme size thus satisfies a deterministic logistic equation in each environmental state $\xi = \pm 1$, subject to a carrying capacity $K \in \{K_-, K_+\}$ that switches according to (S1.7) when the environment changes ($\xi \to -\xi$). Within the $N$-PDMP approximation, the fraction $x$ of $R$ cells in the deme still obeys Eq. (S1.2) and is the slow variable, but $x$ is now coupled to Eq. (S1.8) and hence depends on the time-fluctuating environment encoded in (S1.7).

The stationary marginal probability density of the $N$-PDMP (S1.8) with environmental parameters $\{\nu, \delta\}$, is [5, 6, 13, 19, 20, 21, 3, 4, 11, 2]:

$$\rho(N) = \frac{\mathcal{Z}}{N^2}\left(\frac{K_+ - N}{N}\right)^{\nu(1-\delta)-1}\left(\frac{N - K_-}{N}\right)^{\nu(1+\delta)-1}, \tag{S1.9}$$

where $\mathcal{Z}$ is a normalisation constant and $N \in [K_-, K_+]$ is treated as a continuous variable. Despite ignoring demographic fluctuations, the $N$-PDMP and its stationary density (S1.9) provide a faithful description of the deme size dynamics when it is subject to the time-switching carrying capacity (S1.7) [3, 4, 13, 15, 16, 17, 11, 5, 6, 2], see Fig S1D-F. For instance, the long-time average deme size $\langle N\rangle$ is accurately approximated by $\int_{K_-}^{K_+} N\rho(N)dN$ [3, 4, 11, 13], see Fig S1E.

Guided by the expression of (S1.9) (with $|\delta| < 1$), we distinguish three dynamical regimes:
(i) When environmental switching is much slower than the ecological timescale ("slow switching"), $\nu \ll 1$, $N$ is effectively constant and close to either $K_-$ or $K_+$ with respective probability $(1 \mp \delta)/2$. Hence, when $\nu \ll 1$, the long-time distribution of $N$ is bimodal, a feature well captured by $\rho(N)$, and $(N_R, N_S) \approx (N_{\rm th}, K_\mp - N_{\rm th})$ with probability $(1 \mp \delta)/2$; see Fig S1A,D.

(ii) When the rate of environmental variability is much higher than that of the ecological dynamics ("fast switching"), $\nu \gg 1$, $N$ is not able to track $K(t)$ and environmental fluctuations self-average with the deme size fluctuating about the effective carrying capacity [3, 4, 13, 17, 11, 5]

$$\mathcal{K} = 1/\langle 1/K(t) \rangle = 2K_- K_+ / \left[ (1+\delta)\, K_- + (1-\delta)\, K_+ \right]. \tag{S1.10}$$

Thus, when $\nu \gg 1$, the quasi-stationary distribution of $N$ is unimodal and centred about $\mathcal{K}$, with $N \approx \mathcal{K}$ when $\mathcal{K} \gg 1$, as aptly reproduced by $\rho(N)$, and $(N_R, N_S) \approx (N_{\rm th}, \mathcal{K} - N_{\rm th})$ with probability $(1 \mp \delta)/2$; see Fig S1C,F.

(iii) When the timescale of environmental variability is similar to that of the ecological dynamics ("intermediate switching"), $\nu \lesssim 1, 0 \le \delta \lesssim 1$, the deme size tracks the carrying capacity, with $N_R$ and $N_S$ fluctuating respectively about $N_{\rm th}$ and $K(t) - N_{\rm th}$ ($N_R \approx N_{\rm th}$ and $N_S \approx K(t) - N_{\rm th}$ when $K(t) \gg 1$), but the quasi-stationary distribution of $N$ cannot be simply related to an effective static carrying capacity or to a linear superposition of $K_\pm$. In the intermediate switching regime, the long-time distribution of $N$ is shaped by environmental variability, a property well captured by $\rho(N)$; see Fig S1E. Moreover and quite remarkably, this dynamical regime is characterised by *bottlenecks* when $\nu \lesssim 1, 0 \le \delta \lesssim 1$. Bottlenecks arise when the carrying capacity switches from $K_+$ to $K_- < K_+$, leading to a drastic reduction of the deme size and cause important fluctuations; see Fig S1B. Since the average time spent in the environmental state $\xi = \pm 1$, where $K = K_\pm$, is $1/\nu_\pm$, the mean time between two successive bottlenecks is $1/\nu_- + 1/\nu_+ = 2\nu/(\nu_+\nu_-)$ and therefore bottlenecks occur at a rate $\nu_+\nu_-/(2\nu) = \nu(1-\delta^2)/2$ (average bottleneck frequency) [13, 5].

In the regimes of slow/fast switching, $R$ and $S$ can coexist for extended periods and resistant cells can prevail in an isolated deme [5, 6]. (In Refs. [5, 11, 6], there is long-lived strain coexistence when its duration exceeds $2\langle N \rangle$). However, the intermediate regime where $\nu \sim s \lesssim 1$, $0 \le \delta < 1$, is characterised by population bottlenecks that lead to a likely fluctuation-driven clearance of resistant cells from the isolated deme when $K_+/K_- \gtrsim N_{\rm th}$ (and $1 \ll N_{\rm th} < K_- \ll K_+$) after experiencing a sequence of bottlenecks [5] (Model & Methods), see Figs 1A and S1B. Importantly, the condition $K_+/K_- \gtrsim N_{\rm th}$ depends on the bottleneck strength rather than on the typical deme size in each environment, and it can be satisfied by values encountered in realistic microbial communities, such as $(K_+, K_-, N_{\rm th}) \sim (10^{11}, 10^6, 10^5)$ [5]. The intermediate switching regime, characterised by the "fluctuation-driven eradication" of resistance in a time scaling with $1/s$, corresponds to environmental variations occurring on a similar timescale $1/s$ as the intra-deme dynamics (i.e. the bottleneck frequency is comparable to the rate at which the deme composition changes; see Eq.(S1.2)) [5].

In an isolated deme, the long-time distribution size $N$ is independent of its composition, see (S1.8) and Fig S1. Its approximation by the PDMP density (S1.9) is hence expected to hold also in the presence of migration at any values of the environmental switching parameters $(\nu, \delta)$, as found also in Ref. [2]. In particular, slow migration ($m \ll 1$), the most relevant regime for this study has only a minor influence on the distribution of $N$ [2]. The analysis based on the PDMP approximation therefore holds when $m \ll 1$, and we similarly expect that the PDMP-based description to hold also for intermediate and fast migration rates. For the latter ($m \gg 1$), all $L^2$ demes can be viewed as being fully connected as in the island model [22, 23].

### 2 Coexistence of resistant and sensitive cells in static and under slow/fast switching environmental conditions

It is known that on the one hand migration increases the intra-deme diversity (rising alpha diversity) and, on the other hand, dispersal mixes strains between patches, hence reducing the inter-deme differences and causing spatial homogenization of the population (lowering beta diversity) [24, 25]. In this context, here we assess how spatial migration influences the survival of $R$ and $S$ cells in *static environments*, where the carrying capacity $K = K_0$ is constant and, after a short transient, each deme has approximately the same size $N \approx K_0$ (Sec. 1.2.1). We also assess the cases of "slow switching" ($\nu \ll 1$) and "fast switching" ($\nu \gg 1$) environments, which can be effectively mapped into a linear combination of two static environments ($K = K_-$ and $K = K_+$) and a single static environment ($K = \mathcal{K}$), respectively (see below and Sec. 1.2.3; Fig S1 panels A,D and C,F).

When the grid consists of fully isolated demes that are entirely disconnected ($m = 0$), these evolve independently of each other. In the case of static environments, see Sec. 1.2.2 and Model & Methods,

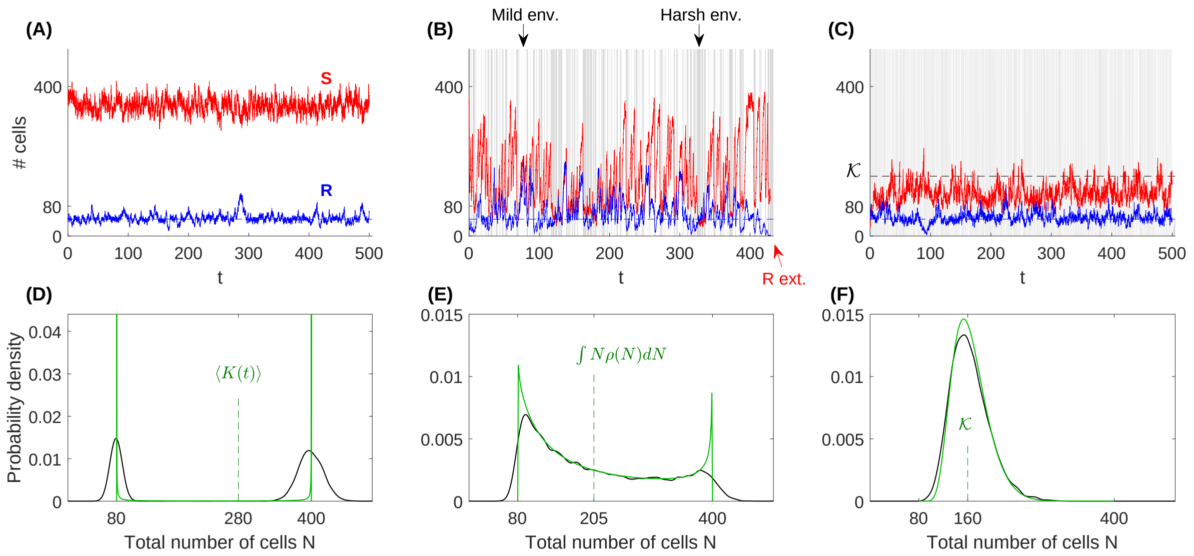


Figure S1: **Microbial dynamics in an isolated deme subject to a switching environment. (A)** Example evolution of the number of $S$ (red curve) and $R$ cells (blue) in an isolated deme (in microbial generation time, see Sec. 3) for a slow-switching environment. Parameters are $N_{\text{th}} = 45$ (blue dashed line), $K_+ = 400$, $K_- = 80$, $\nu = 0.001$, $\delta = 0.25$, $s = 0.1$, and $a = 0.25$. Here, the slow-switching environment stays in the mild state ($K = K_+$), where $N_S \approx K_+ - N_{\text{th}}$ while $N_R \approx N_{\text{th}}$ (Model & Methods and Sec. 1.2.3). The dynamics is thus the same as in a static environment with carrying capacity $K = K(t = 0)$. **(B)** Example realisation as in panel (A) for an intermediate-switching environment with $\nu = 0.75 \sim s$. The environment switches back and forth between harsh ($K = K_-$, grey background shade) and mild states ($K_+$, white shade). Frequent environmental bottlenecks (white-to-grey) are accompanied by a sequence of transient $N_R$ dips leading to the fluctuation-driven clearance of resistance (red arrow, Model & Methods and Sec. 1.2.3). **(C)** Same as in panels (A,B) for a fast-switching environment with rate $\nu = 10$. Environmental variations is so rapid that the carrying capacity self averages, with $K \approx \mathcal{K}$ and $N \approx \mathcal{K} \gg 1$ (not shown), $N_R \approx N_{\text{th}}$ and $N_R \approx \mathcal{K} - N_{\text{th}}$ (Sec. 1.2.3). **(D)** Bimodal quasi-stationary probability density of the total population $N = N_S + N_R$ in an isolated deme, sampled from $10^4$ realisations run for $t = 100$ microbial generations, with the same parameters as in (A) (black, Model & Methods and Sec. 1.2.3). The solid green line shows the stationary PDMP density $\rho(N)$ given by Eq. (S1.9) (Sec. 1.2.3). The dashed vertical green line indicates the average $\langle K(t)\rangle = (K_+ + K_-)/2 + \delta(K_+ - K_-)$, which is close to the average deme size $\langle N \rangle$ (Model & Methods). **(E)** As in (D), with the same parameters as in panel (B). The dashed green vertical line shows the PDMP approximation of the average deme size, computed using Eq. (S1.9) according to $\int_{K_-}^{K_+} N\rho(N)\, dN$. **(F)** Same as in (D) and (E), with the same parameters as in panel (C). The dashed vertical green line indicates $\mathcal{K} = 2K_-K_+/\left[(1+\delta)\, K_- + (1-\delta)\, K_+\right]$, which is close to $\langle N \rangle$ and to its PDMP approximation (Sec. 1.2.3).

the dynamics in each deme thus tends to a coexistence equilibrium of $R$ and $S$, with the number of resistant and sensitive cells fluctuating about $N_R \approx K_0 x \approx N_{\text{th}}$ and $N_S \approx K_0(1-x) \approx K_0 - N_{\text{th}}$. (Note that in Fig S2, $R$ dominates because $K_0 < K_0^*$, i.e. $\frac{N_{\text{th}}}{K_0} > \frac{\ln(1-s)}{\ln(1-a)}$ [5], see Eq. (S1.6) and Sec. 1.2.2). In the absence of migration, there is a long-lived strain coexistence across the metapopulation, with a slow increase in time of the number of $R$-only (along with a few $S$-only) demes (lightest curve $m = 0$ in Figs S2C-D). The fraction of resistant cells ($x \equiv N_R/N \approx N_R/K_0$) at long times thus follows a bimodal distribution for $m = 0$, with a dominant peak at $x = 1$, as $R$ cells take over most isolated demes, and a secondary peak at $x = 0$ corresponding to sites fortuitously taken over by $S$ (darkest curve $m = 0$ in Fig S2A; $t = 1500$ microbial generations).

Demes become interconnected when the cell migration rate increases ($m > 0$). The extinction of either $R$ or $S$ strain from a deme is thus no longer irreversible, since each single-strain $R/S$-only deme can be recolonised by $S/R$ microbes migrating from neighbouring sites ('asterisk' and 'cross' demes in Fig 1B, Model & Methods). Migration in static environments therefore generally enhances and promotes the coexistence of strains across the metapopulation [24, 25]. This is illustrated by Fig S2B where faster migration is shown to promote a large number of demes where $R$ and $S$ coexist for extended periods.

At sufficiently high migration rates (for $m \geq 10^{-2}$ in Fig S2B), deme mixing is sufficient to ensure

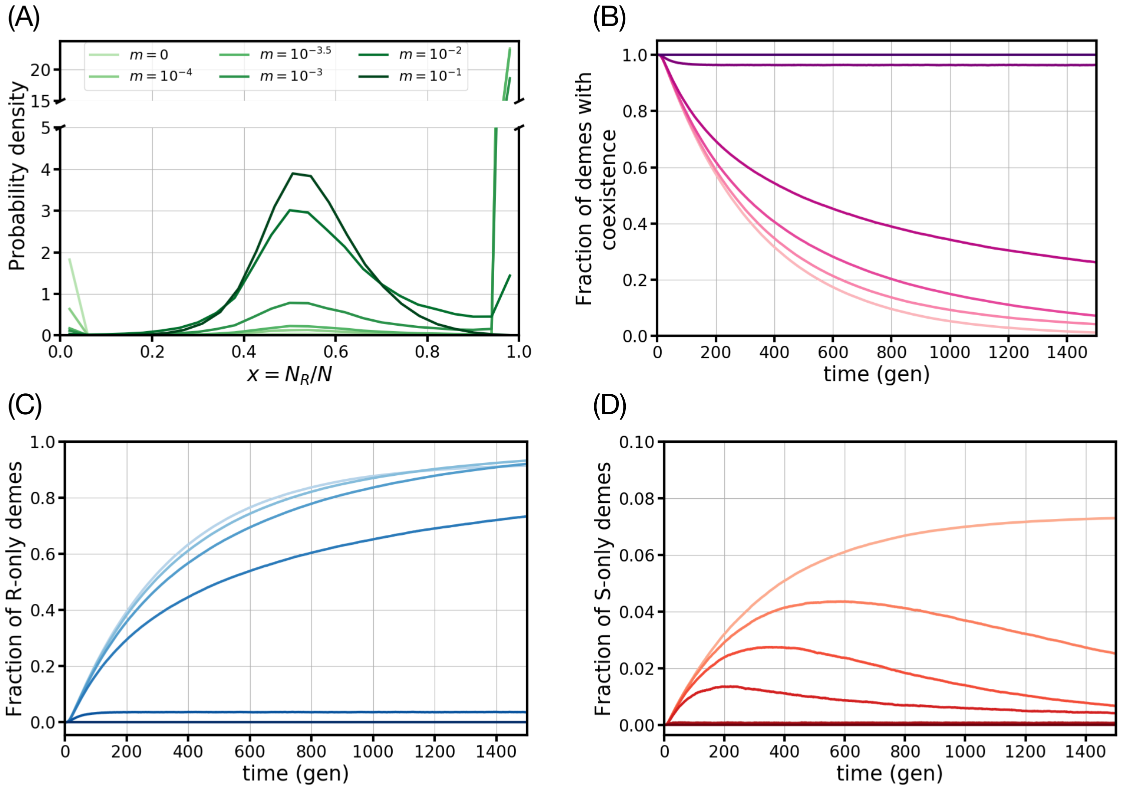


Figure S2: **Migration shapes the coexistence of $S$ and $R$ cells in static environments.** Migration rates are $m \in \{0, 10^{-4}, 10^{-3.5}, 10^{-3}, 10^{-2}, 10^{-1}\}$ (light to dark colour) according to Eq. (2a), the constant carrying capacity is $K_0 = 80$ and the cooperation threshold is $N_{\rm th} = 40$. Other parameters are: resistance metabolic cost $s = 0.1$, drug impact on sensitive growth $a = 0.25$, on a square grid of $L \times L = 100 \times 100$ demes. Data have been averaged over $\mathcal{R} = 50$ realisations, and error bars are smaller than the line thickness. **(A)** Probability density of the microbial composition $x$ giving the fraction of $R$ cells in a deme after $t = 1500$ microbial generations computed in bins of width $\Delta x$=0.04. Clearly, the probability density in is centred around $x \approx N_{\rm th}/K_0 = 0.5$ (where $N \approx K_0$ in each deme), with a sharper peak as $m$ is increased. **(B)** Time evolution of the fraction of demes in the metapopulation with coexisting $R$ and $S$ cells. **(C)** Fraction of $R$-only demes across the metapopulation. **(D)** Fraction of $S$-only demes across the grid. In all panels the migration rate increases from light to darker colour.

the maintenance of coexisting demes, where the fraction of $R$ cells fluctuates about $x \approx N_{\rm th}/K_0$, see Fig S2A. In the limit of fast migration $m \gg 1$, demes can be regarded as fully connected, and the distribution of the fraction $x = N_R/N$ of $R$ cells in a deme concentrates narrowly about $N_{\rm th}/K_0$ (not shown in Fig S2A). After an unobservably long time (when $K_0 \gg 1$, see Secs. 1.1 and 3), the metapopulation will most likely become a homogeneous monoculture of $R$ cells since these typically fixate faster than $S$ cells (Fig S2C,D). As noted in Sec. 1.1, the final state is the full extinction of the metapopulation, but it will be attained after much longer (generally unobservable) time.

Remarkably, we also find that slow-but-nonzero migration rate here enhances the fixation probability of $R$ cells. This is in stark contrast from our key result obtained in *time-varying environments* where we have shown that slow migration promotes the eradication of resistance in the intermediate switching regime (Discussion and Fig S4C). Perhaps counterintuitively, we find that the largest fraction of $R$-only demes in a *static environment* occurs at slow yet nonzero migration (for which the fraction of $S$-only demes is already low). For instance, in Fig S2C the $m = 10^{-4}$ and $10^{-3.5}$ lines overtake the $m = 0$ curve at $t \approx 1100$ and $t \approx 1400$. Although quantitatively small, this effect unveils a qualitatively relevant phenomenon: In a static environment (zero switching), slow-but-nonzero migration is strong enough to foster $R$ recolonisation and sufficiently weak to prevent $S$ recolonisation. When the metapopulation is subject to a time-fluctuating environment in the intermediate switching regime (Sec. 1.2 and Model & Methods), and there is slow cell migration, we have exhaustively discussed in the main manuscript that a similar phenomenon of much stronger effect enhances the eradication of resistance (Results & Discussion, see S2 Movie).

We have therefore shown that the parameters yielding efficient eradication of $R$ in intermediate switching environments (see Figs 2-4 and Figs S6-S10) lead to long-lived coexistence of $R$ and $S$ in static environments, with $R$ gradually dominating (Fig S2; see also Sec. 1.2.2).

All the above results for static environments can readily be used to shed light on the eco-evolutionary dynamics under *time-fluctuating environmental conditions* when the switching rate is either very low ($\nu \ll 1$) or very high ($\nu \gg 1$), i.e. under slow/fast switching environments (Sec. 1.2.3).

- When $\nu \ll 1$ (slow switching): Each deme spends long periods in either the harsh or mild environment, so that its size is $N \approx K_{\pm}$ with probability $(1 \pm \delta)/2$; see Sec. 1.2.3. Therefore, when $K_{+} \gg K_{-} > N_{\rm th} \gg 1$, $K_{-} < K_0^*$, and $K_{+} > K_0^*$ (see Eq. (S1.6)), as in all our examples, the $R$ fixation probability given by Eq. (S1.5) is close to 1 in the harsh environment and vanishingly small in the mild one ($\phi(K_0 = K_{-}) \approx 1$, $\phi(K_0 = K_{+}) \approx 0$). Hence, under these conditions, in the absence of migration, the most likely outcomes are either the fixation of $R$ in the harsh environment, occurring with probability $\frac{(1-\delta)}{2}\phi(K_{-})$, or, with probability $\frac{(1+\delta)}{2}$, the coexistence of $R$ and $S$ in the mild environment, with a fraction $x \approx N_{\rm th}/K_{+}$ of resistant cells per deme. When migration is present ($m > 0$), in the harsh environment, coexistence about $x \approx N_{\rm th}/K_{-}$ becomes more likely than fixation of $R$. Under fast migration, long-lived coexistence of $R$ and $S$ thus dominates, with a typical resistant fraction $x \approx \frac{(1-\delta)}{2}\frac{N_{\rm th}}{K_{-}} + \frac{(1+\delta)}{2}\frac{N_{\rm th}}{K_{+}} = \frac{N_{\rm th}}{\mathcal{K}}$ when averaged across harsh and mild environments, see Eq. (S1.10).

- When $\nu \gg 1$ (fast switching): Each deme experiences an effective carrying capacity $\mathcal{K}$ (Sec. 1.2.3 Eq. (S1.10)). Thus, when $\mathcal{K} > K_0^*$ (as in our examples), we have $\phi(K_0 = \mathcal{K}) \approx 0$, and long-lived coexistence of $R$ and $S$ is almost certain, with a resistant fraction $x \approx N_{\rm th}/\mathcal{K}$ in each deme for any migration rate. Increasing $m$ prolongs coexistence and sharpens the distribution of $x$ around this value.

Moreover, as shown in Ref. [5], the mean fixation time in an isolated deme depends strongly on $N_{\rm th}$ (see also Sec. 1.2.2). When $N_{\rm th}$ is not particularly large (as in our examples) and $K > K_0^*$, long-lived coexistence under no or slow migration is not guaranteed, as there is a finite probability of $S$ fixation [5] after only some hundred microbial generations (see Fig. 3 in Ref. [5]).

The essence of this analysis is illustrated in Fig S3, where $N_{\rm th} = 40$ and the mean fixation time in an isolated deme when $K > K_0^*$ is comparable to the time considered in Fig S3C [5]. In Fig S3A (slow environmental switching, $\nu \ll 1$), when $m = 0$ and $m = 10^{-3.5}$ (slow migration), the spike near $x \approx 1$ corresponds to the fixation of $R$ in the harsh environment, whereas the peak about $x = 0$ stems from the coexistence of both strains in the mild environment (with a small fraction $x \approx N_{\rm th}/K_{+} \approx 0.007$ of $R$) in each deme and also from the possible fixation of $S$. Coexistence becomes predominant at higher migration rates (density more centred about $x \approx N_{\rm th}/K_{-} = 0.5$ in the harsh environment and about $x \approx N_{\rm th}/K_{+} \approx 0.007$ in the mild environment as $m$ increases, see darker curves in Fig. S3A), as migration facilitates the recolonisation of $R$-only and $S$-only demes (see Figs S4B-D and S5A-D). Fig S3B shows that the fraction of $R$-only demes decreases with $m$, and vanishes under fast migration rate (for $m = 0.1$ in Fig S3). In Fig. S3C (fast environmental switching, $\nu \gg 1$), the distribution of $x = N_R/N$ has a clear peak near $N_{\rm th}/\mathcal{K}$ that becomes narrower and sharper for higher $m$, and a spike near $x \approx 0$ when $m = 0$ and $m = 10^{-3.5}$. The latter stems from the fixation of $S$ that cannot be neglected for $N_{\rm th} = 40$ and $\mathcal{K} > K_0^*$ (the mean fixation time in an isolated deme of size $\mathcal{K}$ and $N_{\rm th} = 40$ is comparable to $t = 500$; see Fig 3 of Ref. [5]), and disappears under faster migration. Fig. S3D shows that the fraction of $R$-only demes is almost zero in this fast environmental switching example – both under zero and slow migration ($m = 0$ and $m = 10^{-3.5}$) – and it vanishes under fast migration ($m = 0.1$), when strain coexistence is predominant.

In summary, for all examples considered in this work, long-term coexistence of $R$ and $S$ across the grid is always possible under both slow and fast environmental switching, when there is non-zero migration (coexistence is likely when $m$ is not too small). Fixation of $R$ or $S$ can occur under slow migration rates (or $m = 0$), but coexistence dominates otherwise. In the absence of migration, demes are fully disconnected, and resistance is very likely to persist – either through coexistence or fixation of $R$ – for extended periods under both fast and slow environmental variation.

Moreover, as indicated in the Discussion, in this study we consider a low drug concentration regime. However, cooperative resistance has been shown to be prevalent even at high drug concentrations in

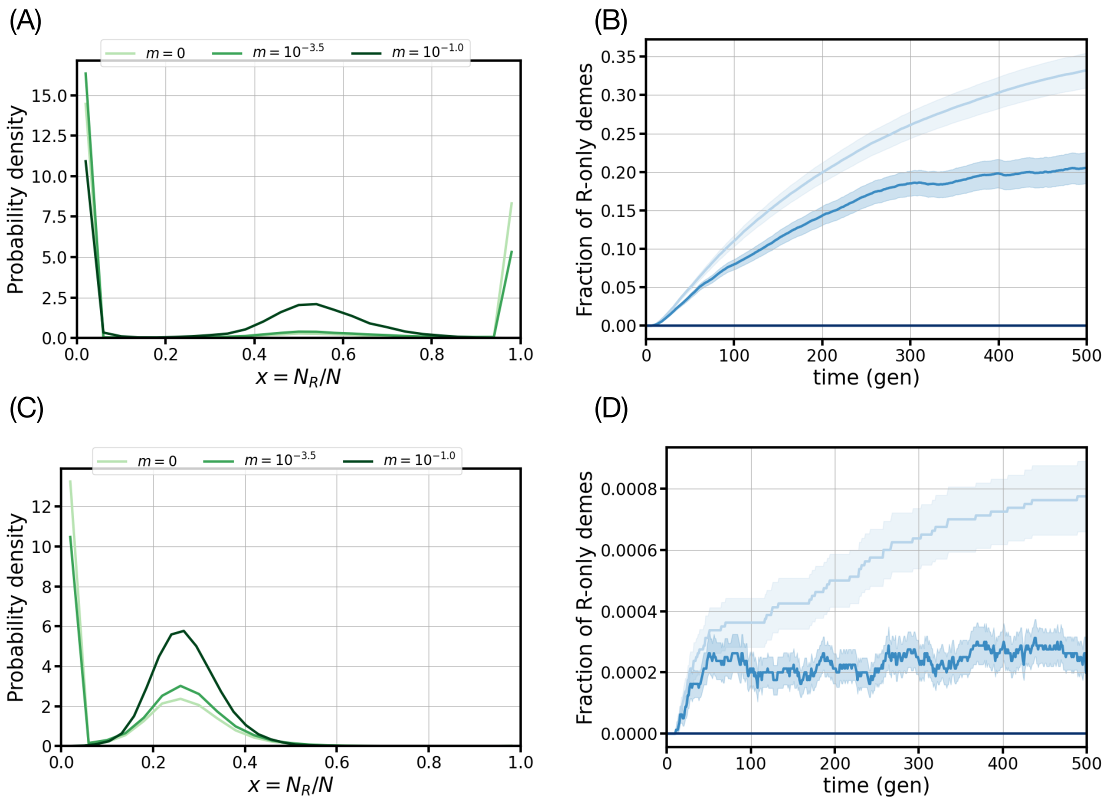


Figure S3: **Comparing migration effects on the coexistence of $S$ and $R$ cells in slow (A,B) and fast (C,D) switching environments.** Eco-evolutionary dynamics of $20 \times 20$ metapopulations with migration according to Eq. (2a), with migration rates $m \in \{0, 10^{-3.5}, 10^{-1}\}$ (light to dark colour). Other parameters are: $K_- = 80$, $K_+ = 5657$, $\delta = 0$ (symmetric switching, no environmental bias), $N_{\rm th} = 40$ (cooperation threshold), $s = 0.1, a = 0.25$. Slow/fast switching rates are $\nu = 0.001$ in (A,B) and $\nu = 10$ in (C,D). Results have been averaged over 200 independent simulations. **(A)** Probability density of the microbial composition $x = N_R/N$ (fraction of $R$ cells in a deme) after $t = 500$ microbial generations computed in bins of width $\Delta x$=0.04 for $\nu = 0.001$ (slow switching). When $m = 0$ (no migration) and $m = 10^{-3.5}$ (slow migration), the probability density is characterised by a spike at $x = 1$ ($R$ fixation) and a stronger spike near $x \approx 0$ (corresponding to $S$ fixation and $S/R$ coexistence with a low fraction $N_{\rm th}/K_+$ of $R$ cells in the mild environment; see Sec. 2). In the presence of faster migration, coexistence is more likely, as indicated by a pronounced peak around $x \approx N_{\rm th}/K_- = 0.5$ in the harsh environment for $m = 0.1$, while the intensity of the spike near $x \approx 0$ decreases with $m$ ($S$ fixation in the mild environment is less likely as $m$ increases). **(B)** Fraction of $R$-only sites in the metapopulation as a function of time for the same parameters as in panel A. Error bars show the standard error of the mean (see Sec. 3.3.2). As the migration rate increases (light to dark blue), the fraction of $R$-only demes decreases to zero, as coexistence is predominant when $m = 0.1$. Results similar to those of Fig S2C. **(C)** As in panel A for fast switching environment, with $\nu = 10$. In this regime, each deme is subject to the effective carrying capacity $\mathcal{K} \approx 158$; see (S1.10). The probability density has thus a sharp peak $x \approx N_{\rm th}/\mathcal{K} \approx 0.25$ for all values of $m$, with the peak becoming sharper and narrower as $m$ is increased (coexistence is most likely under fast migration, as shown here for $m = 0.1$). When $m = 0$ and $m = 10^{-3.5}$, there is also a spike near $x \approx 0$ stemming from the fixation of $S$ that is here possible when $m = 0$ or $m \ll 1$ (since the mean fixation time in an isolated deme is comparable to $t = 500$; see text and Fig 3 of [5]). **(D)** Same as in panel B in a fast environmental switching regime, with $\nu = 10$. The fraction of $R$-only demes is almost zero when $\nu \gg 1$ under zero and slow migration ($m = 0$ and $m = 10^{-3.5}$), and it vanishes under fast migration ($m = 0.1$), when each deme of the grid is characterised by strain coexistence.

well-mixed populations [26]. While this has not been thoroughly investigated in our work, we note that high drug concentration would increase the cooperation threshold $N_{\text{th}}$ (more $R$ cells would be needed to protect $S$ individuals from exposure to the active drug). We have shown that the number of $S$ cells in a deme tends to $N_S \approx K(t) - N_{\text{th}}$ (see Secs. 1.2.1 and 1.2.3). Therefore, sensitive cells would likely become extinct across all demes and $R$ cells fix in the metapopulation as $N_{\text{th}}$ is increased due to a higher drug concentration.

### 3 Computational methods: Monte Carlo algorithm, simulation parameters, additional plotting details

In this section, we explain how the extensive stochastic simulations of the metapopulation dynamics were performed, summarise the parameters that we have used and provide further details on the figures.

#### 3.1 Metapopulation simulations

Spatially extended models of microbial communities are often investigated by means of agent-based lattice simulations [27]. This work employs a stochastic Monte Carlo algorithm on a two-dimensional, $L \times L$, square lattice (or grid) metapopulation with periodic boundary conditions to investigate interaction effects of migration-coupled neighbouring microbial sub-populations (demes), see Model & Methods. In all the examples in the main text, we have chosen to consider grids of linear size $L = 20$, with $L$ larger than the metapopulation composition's correlation length $\ell$ in static environments (having preliminarily checked that the correlation length is thus always much smaller than the lattice size, with $\ell < 5 \ll L$), but low enough to ensure computational feasibility. Monte Carlo algorithms similar to the one used here are particularly useful tools to investigate the properties of spatially extended systems and have been abundantly employed across various fields, see, e.g., Ref. [28]. The Gillespie algorithm [29] (that generate statistically exact sample paths) was also considered, but due to the number of cells within each deme and overall size of the metapopulation, it was deemed less computationally efficient. Each lattice deme contains sensitive and resistant cells whose populations are governed by a birth-death process with migration (Model & Methods), whose transition rates are given by Eqs. (1), (2a) and (2b), and are subject to a fluctuating carrying capacity (S1.7). To minimize initial transients, at $t = 0$ a total of $N_{\text{th}}L^2$ resistant and $(K - N_{\text{th}})L^2$ sensitive cells (with $K \in \{K_-, K_+\}$) are uniformly distributed at random among all $L^2$ demes in the metapopulation. (Similarly, in the one-dimensional lattice of Sec. 5.5, an initial total number of $N_{\text{th}}L$ and $(K - N_{\text{th}})L$ cells of type $R$ and $S$ are distributed across the cycle). The expected initial number of resistant and sensitive cells, denoted by $N^0_{R,S}$, here matches their respective stationary values in a static environment, i.e. $N^0_R = N_{\text{th}}$ and $N^0_S = K - N_{\text{th}}$ (Methods). Further, the environmental state of the metapopulation at $t = 0$ begins at stationarity, $K(t = 0) = \langle K(t) \rangle = \frac{1}{2}[K_+ + K_- + \delta(K_+ - K_-)]$ with the mean of the dichotomous Markov noise equalling that of the environmental switching bias, $\langle \xi(t) \rangle = \delta$ [5, 13]. Thus, the system begins in a harsh ($K_-$) or mild ($K_+$) environment with a probability of $(1 - \delta)/2$ or $(1 + \delta)/2$, respectively (Model & Methods, Sec. 1.2.3). We ran $\mathcal{R} = 200$ realisations for each parameter set of Figs 2A, 4A-D, S6A, S7A, S9A-E, and S10, and performed $\mathcal{R} = 50$ realisations for the parameter sets of Fig S7A,B. This means that for each parameter set, we have generated $k = 1 \dots \mathcal{R}$ realisations (simulation runs) of the metapopulation dynamics and the probabilities shown in the heatmaps of those figures have been obtained by sampling over $\mathcal{R}$ samples. Moreover, the trajectories of the fraction $\rho_S(t)$ of demes without $R$ cells, shown in Figs 2B-H, 3I, S6C-K, and S7C-K, have been computed according to Eq. (7) from one typical sample metapopulation realisation. Similarly, the number $N_{R_c/S_c}(t)$ of $R/S$ cells in coexisting demes across the lattice, shown in Fig S8 of Sec. 5.4, have been computed for a single metapopulation realisation, see Eq. (S5.12).

Each panel of Fig 3 features the same single simulation realisation. Simulation results reported in Figs 1A and S4A for the dynamics in a single isolated deme were obtained using the classical Gillespie algorithm [29].

The system evolves in time units of microbial generations, where we consider one generation to equal one Monte Carlo Step (MCS), e.g., bacteria replicate on a scale of approximately once every $\sim 1$ hour. Within every generation, the environment can switch at rate $\nu$, and cells are chosen at random to attempt birth, death, or migration with rates given by (1), (2a), (2b); see Model & Methods.

The same general Monte Carlo stochastic simulation algorithm used in two-dimensions was applied to the one-dimensional special case (Sec. 5.5). Both versions of the code are electronically available on the Open Science Foundation repository at https://doi.org/10.17605/OSF.IO/EPB28. We have used two forms of migration: one, with transition rate (2a), where the per capita migration rate depends on the deme's local population density; and the other, see (2b), with a simpler constant per capita dispersal rate (Model & Methods). The simulation of migration has been implemented in the same way for both formulations. A single MCS is completed once the sum of death or birth reactions equals twice the initial number of cells within the system at the beginning of the step, i.e. 1 MCS $=2\mathcal{N}= 2\sum_{\vec{u}}(N_{\rm S}(\vec{u}) + N_{\rm R}(\vec{u}))$ where the sum is over all the demes $\vec{u}$, see also below. There are thus $\sim 10^4$ birth/death events in 1 MCS when $N(\vec{u}) = N_S(\vec{u}) + N_R(\vec{u}) \approx K_-$ in each deme $\vec{u}$, and $\sim 10^5 - 10^7$ events when $N(\vec{u}) \approx K_+$. Accordingly, on average, each cell in the system attempts one birth and one death reaction within a generation. Setting the simulation time unit (1 MCS) as one microbial generation, migration reactions and environmental switches do not contribute to the above reaction count. This allows for a direct comparison between simulations at different migration $m$ and environmental rates $\nu$ on the eco-evolutionary timescale of (S1.2) and (S1.8), see also below. All finite stochastic systems will eventually reach their final, absorbing state. In our model, the ultimate absorbing state is characterised by the total extinction of both microbial strains. However, this is unobservable in a reasonable amount of computational time as this phenomenon occurs on a timescale that diverges dramatically with the total population and the size of the grid of demes (Sec. 1.1). Moreover, we have considered values of $K_-$, $K_+$ and $N_{\rm th}$ large enough for demes never to go extinct during our simulations, and able to generate strong bottlenecks and lead to fluctuation-driven resistance in an isolated deme [5] (Results). In our study, we ran simulations for up to 500 generations, which is of the order of $10^2$ experimental hours (Discussion). This is sufficiently long to observe the fluctuation-driven eradication of $R$ cells in the metapopulation (when feasible), while maintaining computational efficiency.

##### 3.2 Simulation parameters

To run a single simulation, 12 parameters are specified: the side length $L$ of the lattice of demes; the duration of the simulation (number of microbial generations), $t_{\rm max}$; the average initial number of sensitive and resistant cells per deme at $t = 0$, $N_S^0$ and $N_R^0$; the impact of the drug on the fitness of exposed sensitive cells, $a$; the constant metabolic cost for resistant cells to generate the resistance enzyme, $s$; the migration rate, $m$; the mild and harsh carrying capacity in each deme, $K_+$ and $K_-$; the resistant cooperation threshold, $N_{\rm th}$; the average environmental switching rate, $\nu$; and the environmental bias, $\delta$ (see Table 1). Throughout this work, we fixed seven parameters in all lattice simulations for computational convenience: $L = 20$ for the grid of Figs 2-4 ($L = 100$ for the cycle of Sec. 5.5), $t_{\rm max} = 500$ (but $t_{\rm max} = 1500$ for Fig S2), $N_R^0 = 40 = N_{\rm th}$, $a = 0.25$, $s = 0.1$, and $K_- = 80$. We set the average initial number of $S$ cells per deme to $N_S^0 = K(t = 0) - N_R^0$, where the starting carrying capacity $K(0)$ is either $K_-$ or $K_+$ with probability $(1 - \delta)/2$ and $(1 + \delta)/2$, respectively (Sec. 1.2.3). The remaining parameters are specified within the figure captions. All parameters are summarised in Table 1 of the main text. In our figures, we have explored and characterised the spatial fluctuation-driven eradication of $R$ across the two-dimensional metapopulation by tuning the average switching rate $\nu$, the environmental switching bias $\delta$ determining the relative time spent in mild/harsh conditions, the migration rate $m$, and the bottleneck strength $K_+/K_-$ (keeping $K_-$ fixed). We proceeded similarly to obtain the results reported in Fig S9 for a one-dimensional metapopulation.

The explored range of migration rates varies from small to large migration values, $m \in [10^{-5}, 10^{-1}]$, as well as the case of absent migration, $m = 0$ (separated by vertical dashed white lines in Figs 2, 4, S6A, S7A, S9A-E, and S10). We simulated a range of values of the demes' carrying capacity $K_+$ in the mild environment, with $K_+$ spanning from $10^3$ to $3.2 \cdot 10^4$ (Discussion). The environmental bottleneck strength $K_+/K_-$ thus ranged from 12.5 to 400. The upper limit in $K_+$ and the side length of the square grid $L$ set the maximum total number of cells across the grid of demes ($\sim L^2 K_+$), which was bounded by $\sim 10^7$ due to computational constraints (Discussion). We also tested an extended range of intermediate environmental switching parameters to corroborate our results, with $\nu \in \{0.01, 0.1, 1\}$ and $\delta \in \{0.25, 0.5, 0.75\}$, see Fig S10, as well as $\nu \in \{0.001, 10\}$ for $\delta = 0$, see Fig S3.

#### 3.3 Additional plotting details

##### 3.3.1 Indicating harsh environment as green bands

In many panels of Figs 3 and S5-S8, green bands are shown to indicate when and for how long the metapopulation is experiencing a harsh environment (where $K = K_{-} = 80$). As outlined in Sec. 3.1, a single generation is completed when a combination of $2\mathcal{N}$ birth or death reactions are attempted, which can be envisioned as the number of ticks of the "Monte Carlo clock". Hence, there are $2\mathcal{N}$ ticks within one generation: the clock ticks forward once at every birth or death reaction. As explained above, environmental switches or migration events are not considered ticks so the clock is not moved forward when these occur. The duration of a generation is thus comparable with other metapopulation realisations with similar system sizes, but whose environmental switching and migration rates differ. The system's observables, such as the population density of $S$ and $R$ across the entire metapopulation $n_{S/R} = \left(\sum_{\vec{u}} N_{S/R}(\vec{u})\right)/L^2$ (fraction of $S$ and $R$ across the grid), the number $N_{S/R}(\vec{u})$ of $S$ and $R$ cells in deme $\vec{u}$, and the current carrying capacity ($K \in \{K_{-}, K_{+}\}$), are recorded at the end of each generation. Therefore, the green bands seen within Figs 3G-I, S5D,H,L, S6C-K, S7C-K, and S8, indicate the last environmental state that the system is experiencing after the final generational tick. These are intended to give the reader additional context to interpret the figure panel, but attention should be paid to the fact that, as explained below, they do not fully reflect the complete environmental switching dynamics.

Monte Carlo (MC) simulations require a suitable time-discretisation and the introduction of an elementary MC step (1 MCS) for which this work defines to be a single generation. The latter acts like a resolution limit: the MC simulation cannot resolve dynamical events happening faster than 1 MCS. Here, 1 MCS is the time for $2\mathcal{N}$ birth-death events (Sec. 3.1) to have occurred across the metapopulation ($\mathcal{N}$ is fixed and determined at the beginning of the step). This is in contrast to other stochastic simulation algorithms, such as the Gillespie algorithm [29], where each event time is drawn individually. During 1 MCS, the next event (environmental switch, birth, death, or migration event) depends on the current propensities of the metapopulation (current overall population on the grid) which are updated after every event. The resolution limit implies that the MC algorithm is well-defined for slow to intermediate environmental switching (for $\nu \lesssim 1$). However, MC algorithm attains its resolution limit when $\nu > 1$. In this case, the likelihood of more than one switch occurring within a single generation becomes non-negligible, and the green bands cannot discern the associated environmental variations (e.g., it cannot distinguish between $K_{\pm} \to \cdots \to K_{\pm}$ and no switches at all), and hence typically shows less time than expected in the harsh environment for a given value of $\delta$ when $\nu > 1$. In other words, the green bands in Figs 3G-I, S5D,H,L, S6C-K, S7C-K, and S8, do not capture the time shorter than 1 microbial generation (1 MCS) spent in the harsh environmental state, and hence provides a partial description of the environmental switching dynamics. Here, most of our results for environmental switching have been obtained for $\nu \lesssim 1$, and we have compared the predictions of the MC algorithm when $\nu > 1$ against analytical results obtained in the fast switching regime, with $\nu \gg 1$ (Model & Methods, Sec. 1.2.3). We have also checked our results against those obtained from the Gillespie algorithm for small systems. This analysis indicates that the MC algorithm that we have used gives quantitatively faithful and accurate results for the range of environmental parameters considered in this study.

##### 3.3.2 Wald interval

Figs 2, 4, S6, S7, S9, and S10 explore a vast region of the parameter space to show the optimal bottleneck strength ($K_{+}/K_{-}$) and migration rate ($m$) for extinction of $R$ cells through the fluctuation-driven eradication mechanism. Each ($m$, $K_{+}/K_{-}$) value pair shown in Figs 2A, 4A-D, S6A, S9A-E, and S10E,G is an average of 200 independent simulations at a single time, $t$. Each ($m$, $K_{+}/K_{-}$) value pair shown in Figs S7A and S10A-D,F,H,I is an average over 50 independent simulations. To compute the probability of $R$ eradication at time $t$, denoted by $P(N_R(t) = 0)$, we checked whether the total lattice population of $R$ cells, $N_R(t)$, was zero or not in each realisation. Accordingly, the error bars in Figs 4E, S6B, S7B, and S9F, represent binomial confidence intervals computed using the Wald method. If $n$ denotes the number of realisations with $N_R(t) = 0$ (successful eradication) and $\mathcal{R}$ the total number of realisations (here $\mathcal{R} = 200$ or 50), then the confidence interval is given by

$$P(N_R(t) = 0) = \frac{n}{\mathcal{R}} \pm \frac{z_\alpha}{\sqrt{\mathcal{R}}}\sqrt{\frac{n}{\mathcal{R}}\left(1 - \frac{n}{\mathcal{R}}\right)}, \tag{S3.11}$$

where $z_\alpha = 1$, corresponding to one standard deviation (approximately a 68% confidence interval), was used for all values of the migration rate $m$. We have used (S3.11) to assess the standard error of the mean for the results reported in Figs 2, 4, and Figs S3, S6-S10, that is estimated to be below 4% when the average is over $\mathcal{R} = 200$ realisations and below 7% when $\mathcal{R} = 50$. The code to reproduce the heatmaps seen in Figs 2A, 4A-D, S6A, S7A, S9A-E, and S10, the complementary time evolution of the $R$ eradication probability plots (Figs 4E, S6B, S7B, and S9F), and the implementation of Eq. (S3.11) can be found on the Open Science Foundation repository, and electronically available at https://doi.org/10.17605/OSF.IO/EPB28.

##### 3.3.3 Calculation of the 90th and 95th percentile of the $R$ eradication time

Fig 4F shows the plots of the 90th and 95th percentile of $R$ eradication time as a function of migration, that is $\tau_{90}(m)$ and $\tau_{95}(m)$, respectively. In this example, the environment switches at a rate $\nu = 0.1$, with an environmental bias $\delta = 0.5$ and the bottleneck strength is $K_+/K_- = 400$. Additional simulation parameters can be found in Table 1.
Here, given a particular percentile, $q \in [0, 100]$, and a migration rate, $m$, the $q$th percentile of resistance eradication time, denoted by $\tau_q(m)$, was computed by (i) recording the first time step in which $R$ eradication occurs in every of the $\mathcal{R}$ realisations ($\mathcal{R} = 200$ in this case). If a realisation does not have complete $R$ eradication within the allotted simulation time ($t_{max} = 500$ generations), then a NaN is recorded instead. Next, the recorded eradication times are (ii) sorted from least to greatest with all NaN entries being placed at the end of the list. Finally, the $\lfloor(\frac{q}{100} \cdot \mathcal{R})\rfloor$th entry of the list is (iii) selected to be the time in which $q$ percent of realisations experience $R$ eradication. Within Fig 4F, for small and large migration rates, it can be seen that the $R$ eradication time exceeded the simulation time, hence no data point is included for $\tau_{95}\left(m \in \{0, 10^{-5}, 10^{-1}\}\right)$ and $\tau_{90}\left(m \in \{0, 10^{-1}\}\right)$. Specifically, we have checked separately the case $m = 0$ and always found that $\tau_q(m = 0) > 500$ and thus $\tau_q(0) > \tau_q(m^*)$ (see "Slow migration can speed up and enhance $R$ eradication: Near-optimal conditions for resistance clearance" in Results).

### 4 Supplementary simulation movies

In this section we describe the supplementary movies, also uploaded in the Open Science Foundation repository and electronically available at https://doi.org/10.17605/OSF.IO/EPB28. The movies have been tested for compatibility on Chrome and Firefox (while they may not play on Safari).

**S1 Movie: Resistant cells can survive in switching environments when demes are fully isolated.** Example of spatial microbial dynamics for a single realisation of a grid metapopulation without migration ($m = 0$) subject to a intermediate switching environment (Model & Methods, Results). The simulation parameters are $L = 20$, $\nu = 0.01$, $\delta = 0$, $K_- = 80$, $K_+ = 1414$, $N_{\text{th}} = 40$, $s = 0.1$ and $a = 0.25$; all the other parameters are as listed in Table 1. The simulation was run for 500 Monte Carlo Steps (i.e., microbial generations; Sec. 3). **Left:** microbial composition of the $20 \times 20$ square grid of demes evolving in time. Red pixels indicate demes with $S$ cells only, blue pixels depict $R$-only demes, and pink pixels are demes where both $R$ and $S$ cells coexist. **Top right:** temporal evolution of the average number of $S$ (red trace) and $R$ cells (blue) over all multi-strain demes (pink demes in left panel) where both strains coexist. **Bottom right:** temporal evolution of the fraction of single-strain demes where only $S$ (red traces) or $R$ cells (blue) survive (compare to Fig 3). Since demes are fully isolated (no microbial migration $m = 0$), the extinction of $R$ or $S$ cells is irreversible in each deme (when a pink pixel in left panel becomes red or blue, it remains so as no recolonisation is possible; blue and red traces in the bottom right panel do not decrease in time). In persistent harsh conditions (low total number of cells, top right panel), $R$ can by chance take over some demes due to demographic fluctuations, while $S$ only takes over a few demes (several blue pixels and a few red pixels replace pink pixels in left panel, blue trace increases faster than red trace in bottom right panel, Results). In a mild environment (high total number of cells, mostly $S$, top right panel), $S$ can still take over a few demes, but $R$ typically cannot (a few additional red pixels replace pink ones in left panel, the red trace still increases while the blue stays constant in the bottom right panel; Results). When the metapopulation experiences an environmental switch from mild to harsh conditions (occurrence of a bottleneck), the fluctuation-driven eradication mechanism wipes out resistance in many coexisting demes (many pink pixels become red in left panel, dip in the red and blue traces in top right panel, big red spike in the red trace of the bottom right panel; Results). After a sufficiently long time, following a sequence of bottlenecks, the metapopulation will be composed of many $S$-only demes (red pixels in left panel) and fewer, spatially scattered $R$-only demes (blue pixels), without any demes where both strains coexist (no pink pixels). For these set of parameters, resistance thus survives in the metapopulation.

**S2 Movie: Slow migration can enhance the fluctuation-driven eradication mechanism.** Legend is as in S1 Movie but with slow cell migration at rate $m = 0.001$, implemented according to the transition rate (2a). Local strain extinction is now reversible as $R$ and $S$ can recolonise any deme through cellular migration (blue and red pixels can become pink again in left panel, traces can decrease in time in the bottom right panel; Results). Recolonisation events are greatly enhanced when the environment switches to the mild state $K = K_+$ since the number of migration attempts increases with the total population size (higher total number of cells, mostly $S$, in top right panel, almost all the pixels become pink in the left panel; both traces almost reach zero again in the bottom right panel; Model & Methods and Results). As in S1 Movie, population bottlenecks lead to fluctuation-eradication of resistance in many coexisting demes (many pink pixels suddenly become red in left panel, dip in red and blue traces in top right panel, red spike in bottom right panel; Results). This process continues cyclically until $R$ cells are eradicated across the whole metapopulation, an outcome that is impossible to achieve for these parameters in the absence of migration (Results and Discussion). Note that full $R$ eradication is not shown in this movie as it is expected to occur some time after $t = 500$.

**S3 Movie: Fluctuation-driven eradication of $R$.** Legend is as in S1-S2 Movies, but for environmental parameters $(\nu, \delta) = (0.1, 0.5)$, high carrying capacity $K_+ = 2000$, and migration rate $m = 0.001$. This movie corresponds to the same realisation that is shown in Fig 3. In this case, the slow-but-nonzero migration rate $m$ and the shorter time spent in the harsh environment than in the mild one ($\delta > 0$) prevents $R$ cells to take over any demes (no blue pixels in left panel and zero blue curve in bottom right panel at all times). The intermediate environmental switching causes many successive bottlenecks leading to the elimination of $R$ from an increasing number of demes, hence overcoming cellular mixing driven by migration (accumulation of burst of red pixels in left panel, accumulation of spikes in the red trace of the bottom right panel; Results and Discussion). Ultimately, after $t = 500$ microbial generations, the $R$ strain is almost entirely wiped out from the metapopulation, and will be fully cleared after a few more generations, which is not shown here due to the set time cap (only two red pixels remain in the left panel, red trace almost reaches 1 at $t = 500$ in the bottom right panel). An example of full eradication within $t = 500$ is shown in Fig S5E-H.

**S4 Movie: Fluctuation-driven eradication mechanism with density-dependent migration.** Legend as in S3 Movie, but with high carrying capacity $K_+ = 16000$ and migration rate $m = 0.01$. As in S1-S3 Movies, the density-dependent migration is implemented according to the transition rate (2a). This movie corresponds to the realisation shown in Supplementary Fig S6I. In this case, the observed behaviour is similar to that of S3 Movie: $R$ cannot take over any deme (no blue pixels in left panel, and no visible blue curve in bottom right panel) and the fluctuation-driven eradication almost eliminates resistance after 500 generations (very few remaining pink pixels in the left panel, red trace almost reaches 1 in the bottom right panel). Full $R$ eradication will occur in the next few microbial generations (not shown here due to the $t = 500$ time limit).

**S5 Movie: Fluctuation-driven eradication mechanism with density-independent migration.** Legend and parameters as in S4 Movie, but here migration is density-independent according to the transition rate (2b). This case corresponds to the realisation shown in Fig S7I. A direct comparison with S4 Movie reveals qualitatively similar results in all panels.

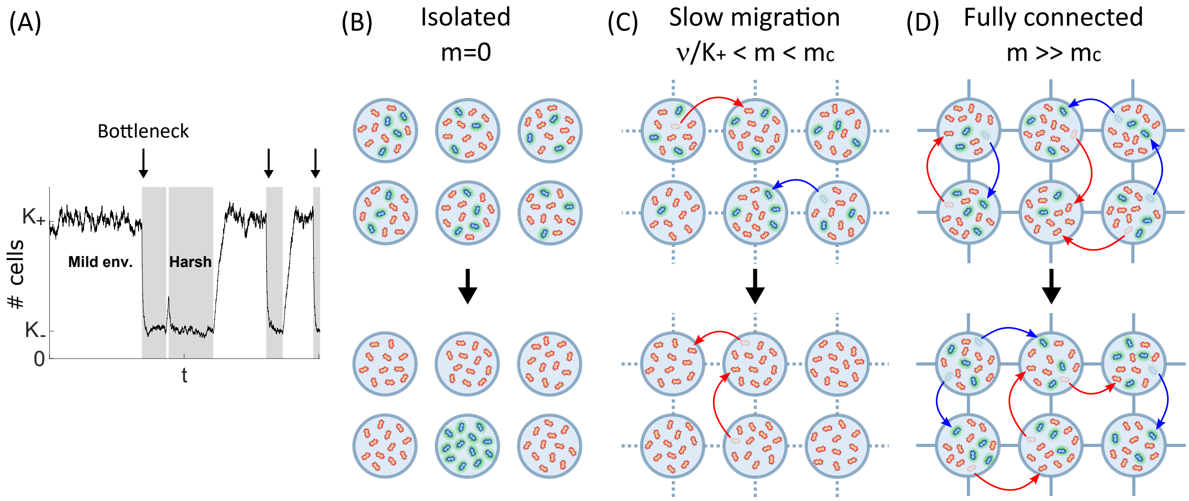


Figure S4: **How slow migration enhances the eradication of resistant cells.** **(A)** Example evolution of the total number of cells in a deme ($N = N_S + N_R$), driven by the fluctuating carrying capacity $K(t)$, in an environment switching at intermediate rates $\nu_\pm \lesssim 1$. As the environment continuously alternates between mild and harsh conditions, the carrying capacity varies suddenly from $K = K_+$ (mild, white background) to $K = K_- \ll 1$ (harsh, grey background), bottlenecks appear (arrows), and the fluctuation-driven eradication of $R$ cells sets in (Model & Methods, Results and Discussion, see also Fig 1). Here, parameters are $\nu_- = 0.125$, $\nu_+ = 0.075$, $K_- = 80$, $K_+ = 400$. **(B)-(D)** schematically illustrate the state of the metapopulation before (top row) and after (bottom) several consecutive bottlenecks in the regime intermediate switching (Model & Methods, Results, Sec. 1.2.3) for three migration scenarios. **(B)** The metapopulation initially consists of demes containing both $R$ and $S$ cells (top, blue and red cells). After several environmental bottlenecks (black downward arrow, $N$ dynamics as panel (A)), the fluctuation-driven eradication mechanism can clear resistance from most demes (Model & Methods). However, when $m = 0$ (no migration), $R$ cells may randomly take over a few demes (bottom, blue-only; see S1 Movie) and survive locally (see Fig S5A-D). (Following bottlenecks, $R$ and $S$ could still coexist in some demes, and more bottlenecks would be needed to eradicate $R$, this is not shown here). **(C)** Same as in (B) with demes connected by slow migration of cells (red/blue curved arrows indicating $S/R$ migrations, $\nu/K_+ < m < m_c$; see conditions (8) and (9) in Results and Discussion). In this scenario, even if $R$ cells take over some demes, $S$ cells can migrate and recolonise those (S2 Movie), which then become prone to fluctuation-driven eradication of resistance (bottom, all red, $m = 10^{-4} - 10^{-3}$ in Fig S5E-H, Results and Discussion). **(D)** Same as in (B)-(C) when cells migrate at a high rate ($m \gg m_c$,), where the metapopulation effectively consists of fully-connected demes. In this dispersal regime, many migration events occur (red and blue arrows) and continuously mix up the composition of the demes, hence preventing the fluctuation-driven eradication of resistance; $R$ and $S$ cells typically coexist in all demes (see Fig S5I-L).

### 5 Supplementary figures

In this section we present and discuss supplementary figures corroborating a number of aspects of our Results and Discussion.

#### 5.1 How slow migration enhances the eradication of $R$

The figures of this subsection illustrate the critical role of migration on the eradication of resistance when each deme is subject to the environmental binary switching shown in Fig S4A, see also Sec. 1.2.

In the absence of migration ($m = 0$), demes are disconnected and the fluctuation-driven eradication of resistance occurs in much of the metapopulation, but resistance may, by chance, take over some demes. Without migration, these demes cannot be recolonised by $S$ cells, as schematically shown in Fig S4B. This intuitive picture is corroborated by the simulation results reported in Fig S5A-D: the blue pixels in the snapshots of Fig S5A-C indicate the persistence of $R$-only demes across the grid when $m = 0$, while Fig S5D shows that there is rapidly a steady fraction of approximately 10% of $R$-only demes ($\rho_R \approx 0.1$ after $t \approx 100$ microbial generations; see Eq.(7)).

When $\nu/K_+ < m < m_c$ (slow migration; see Eq. (6) and conditions (8) and (9)), demes are

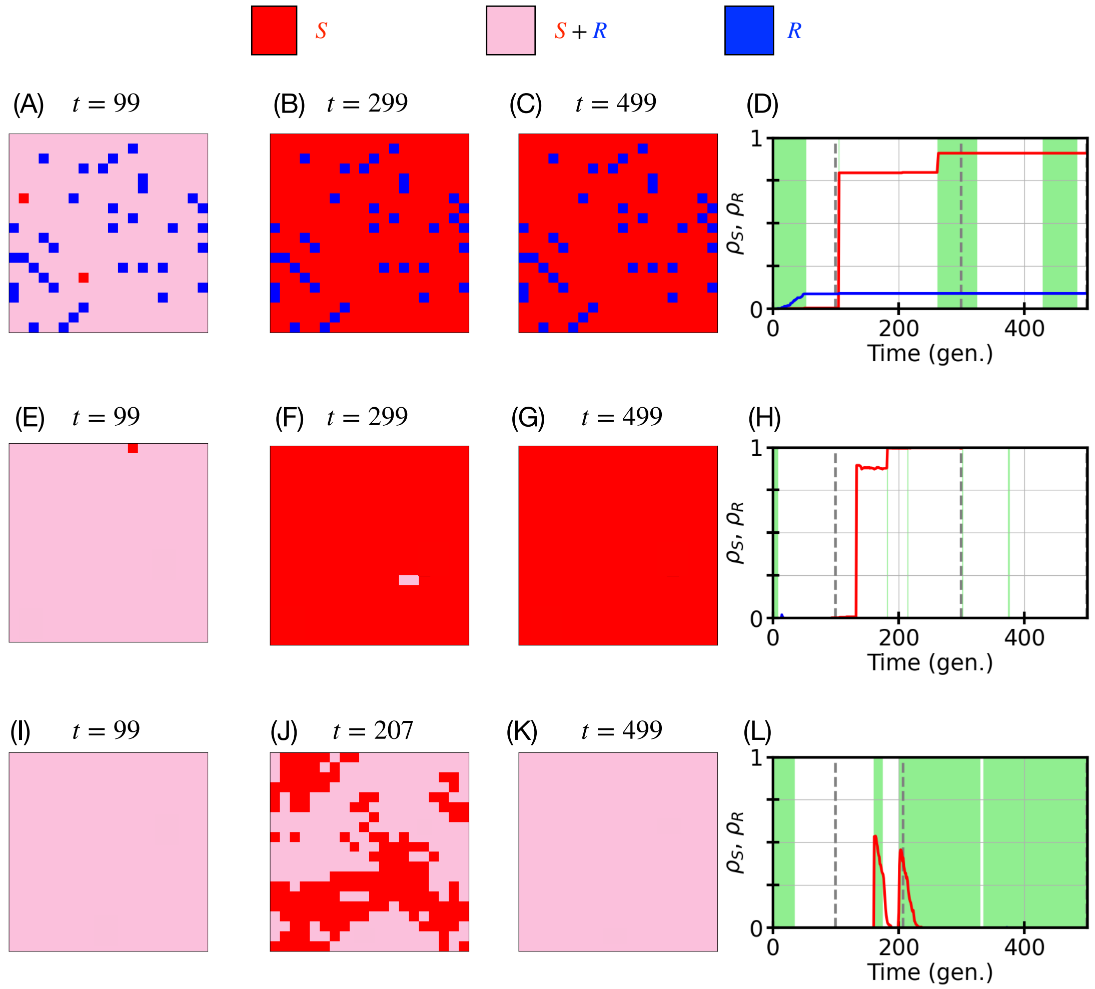


Figure S5: **The role of migration on the eradication of resistance.** Three typical independent realisations of the grid metapopulation eco-evolutionary dynamics (top, middle, and bottom row) corresponding to the three different migration scenarios of Fig S4B-D (with dispersal implemented according to the migration transition rate (2a)). Each realisation begins in the harsh environment ($K_- = 80$) where the average populations of $S$ and $R$ are initially 40 cells per deme; the grid size is $L \times L$, with $L = 20$. Top: No migration ($m = 0$), $K_+ = 22627$, environmental parameters are $(\nu, \delta) = (0.01, 0.25)$. Middle: Migration rate is $m = 10^{-3.5} \approx 3 \cdot 10^{-4}$ (slow migration), $K_+ = 32000$, $(\nu, \delta) = (0.01, 0.75)$. Bottom: $m = 10^{-1}$ (fast migration), $K_+ = 16000$, $(\nu, \delta) = (0.01, 0)$. See Table 1 for the other parameters. **(A-C, E-G, I-K)** Snapshots of the metapopulation at the specified times. Each deme can be of three types: demes with $S$ cells only (red pixels), $R$-only demes (blue pixels), and demes where $S$ and $R$ cells locally coexist (pink pixels). **(D, H, L)** Fractions $\rho_S$ (red) and $\rho_R$ (blue) of $S/R$-only demes across the metapopulation as a function of time (see Eq.(7)). Green bands indicate when the metapopulation is in the harsh environment (Sec. 3.3.1). Vertical dashed lines indicate the respective snapshot times of panels (A-C,E-G,I-K). We find that complete clearance of $R$ cells occurs for slow migration (middle, panels E-H), when, according to the scenario sketched in Fig S4C, there is fluctuation-driven eradication of resistance from the entire grid. For no migration, there is resistance clearance in much of the metapopulation, but a finite fraction of $R$-only demes survives (top, panels A-D). In the absence of migration, resistance is not entirely eradicated from the grid as schematically shown in Fig S4B. When migration is fast, the fluctuation eradication mechanism is countermanded by highly motile $R$ cells recolonising $S$-only areas of the metapopulation (as sketched in Fig S4D), resulting in the metapopulation consisting of coexisting demes (bottom, panels I-L).

connected by a few migration events occurring at a similar rate as environmental bottlenecks (Results), which efficiently promotes the recolonisation of $R$-only demes by $S$ cells and enhances the fluctuation-driven eradication of $R$ across the entire grid (Results, Discussion), as sketched in Fig S4C. This schematic picture is supported by the simulation results of Fig S5E-H, where there are red ($S$-only) and pink ($S$ and $R$ coexistence) demes, but no blue ($R$-only) in the snapshots of Fig S5E-G. We also note that the pink/coexistence demes of Fig S5E, are replaced by red/$S$-only demes in Fig S5F-G, indicating the fluctuation-driven eradication of resistance from the grid, achieved after two bottlenecks (arising after $t \approx 120$ and 180 microbial generations). This is illustrated by Fig S5H, where we find the fraction of $S$-only demes reaching one after $t \approx 180$, i.e. $\rho_S(t \gtrsim 180) = 1$ (while $\rho_R = 0$). Note that according to (8), fluctuation-driven eradication of $R$ occurs whenever $m < m_c$ in the regime of intermediate switching ($\nu \lesssim 1, 0 \leq \delta \lesssim 1$). However, when $m < \nu/K_+$ the migration is too slow to enhance the clearance of $R$ (not enough recolonisation events) beyond its fluctuation-driven eradication in the absence of migration. In other words, the probability of $R$ eradication when $m < \nu/K_+$ is the same as for $m = 0$ (Results).

When $m \gg m_c$ (fast migration, see (6) and conditions (8) and (9)), migration efficiently mixes up the demes and "homogenises" the metapopulation (Results, Discussion and Sec. 2 above). This prevents the clearance of resistance and the metapopulation thus consists of fully connected demes made up of resistant and sensitive cells, as intuitively shown in Fig S4D. This picture is corroborated by the data of Fig S5I-L: The snapshots of Fig S5I-K are dominated by pink/coexistence demes, while Fig S5L clearly shows that in the long run there are no $R/S$-only demes across the metapopulation ($\rho_R \to 0, \rho_S \to 0$). Moreover, as explained in Results, in the regime of intermediate switching ($\nu \sim s \lesssim 1, 0 \leq \delta \lesssim 1$), fluctuation-driven eradication of resistance occurs for very strong bottlenecks, $K_+/K_- \gtrsim N_{\rm th}L^2$, regardless of the value of $m$ (not shown in Figs S4 and S5).

Figure S4 and the results of Fig S5 therefore intuitively and quantitatively explain how the fluctuation-driven eradication of resistance can work efficiently on a lattice metapopulation in the presence of migration, and why — rather surprisingly — slow migration enhances this stochastic phenomenon beyond the case of no migration.

#### 5.2 Fluctuation-driven $R$ eradication with density-dependent migration

In complement to Fig 4 of the main manuscript, we present here additional results to shed further light into the eco-evolutionary dynamics of the grid metapopulation in the regime of intermediate switching (Results, Discussion).

In Fig S6, we revisit the fluctuation-driven eradication of $R$ cells in the case of Fig 4 by focusing on the dynamics leading to clearance of resistance from the metapopulation when the conditions (8) are satisfied. Panels C-K of Fig S6 show how the fraction $\rho_S(t)$ of demes consisting only of $S$ cells (i.e. from which $R$ has been eliminated; see Eq.(7)) varies in time for different values of the parameters $m$ and $K_+/K_-$ in a typical realisation of the metapopulation dynamics.

In panels C and D of Fig S6 (for $m = 0$), we have $\rho_S \to 1$ and thus $R$ clearance after $t \approx 200$ microbial generations. In Fig S6F, we find that for a slow migration rate ($m \approx 10^{-3}$), resistance is cleared after $t \approx 150$, with $\rho_S$ approaching 1 in a "zigzag" pattern due to recolonisation events following bottlenecks. The fact that $R$ eradication occurs faster in Fig S6F than in Fig S6C,D can be traced back to slow migration enhancing the fluctuation-driven eradication of $R$ cells (see Eq. (9)), as discussed in Results and Discussion. In panels E, G, H, J and K of Fig S6, $\rho_S < 1$ after $t = 500$ microbial generations, and therefore in all these cases resistance is still present in the metapopulation after $t = 500$. When $\rho_S < 1$, $R$ and $S$ coexist on a fraction $1 - \rho_S$ of the grid (here $\rho_R \approx 0$), see Fig 3, and the makeup of the coexisting demes is discussed in Sec. 5.4; see Fig S8.

#### 5.3 Fluctuation-driven $R$ eradication with density-independent migration

The results reported in Figs 2-4 and Figs S2, S3, S5, and S6 for the fluctuation-driven eradication of resistance have been obtained with density-dependent migration whose transition rate is given by (2a). In Fig S7 we present the results obtained for the same parameters as in Fig S6, but obtained with *density-independent* migration implemented according to the (simpler) transition rate (2b).

The comparison of Figs S6 and S7 shows a striking resemblance, with similar results obtained in both cases of density-dependent (Fig S6) and density-independent (Fig S7) migration. This indicates

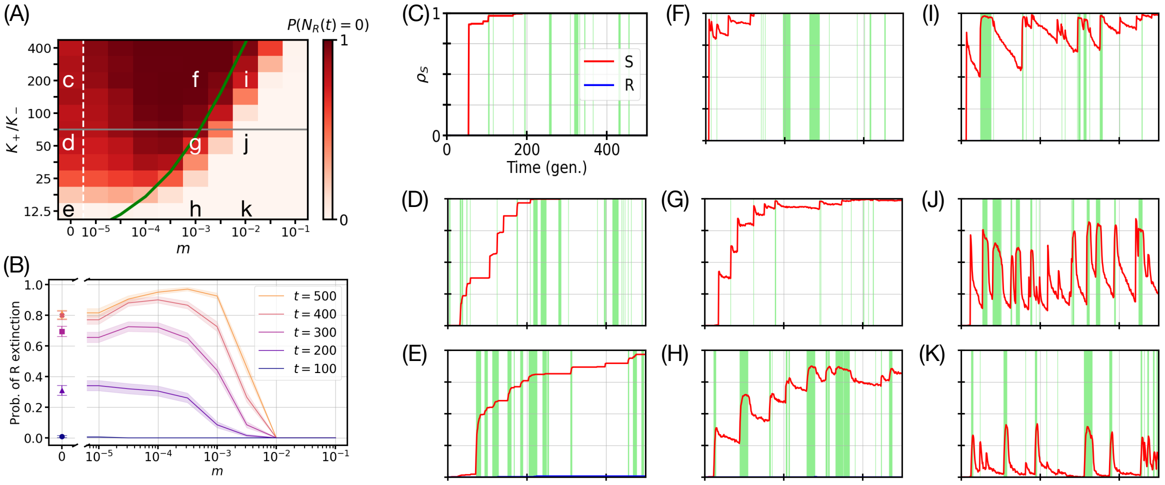


Figure S6: **Revisiting the fluctuation-driven eradication of $R$ cells for intermediate environmental switching with density-dependent migration.** Environmental parameters are $(\nu, \delta) = (0.1, 0.5)$ and density-dependent migration was implemented according to Eq. (2a). All other parameters $(L, s, a, t_{max}, N_S^0, N_R^0, K_-, N_{\text{th}})$ are as listed in Table 1 (Discussion). **(A)** Same heatmap as in Fig 4D showing the probability $P(N_R(t) = 0)$ of total extinction of $R$ (resistant) microbes as a function of bottleneck strength, $K_+/K_-$, and migration rate $m$ after 500 microbial generations ($t = 500$). The colour bar in panel (A) varies from light to dark red, with the darkest red corresponding to eradication of $R$ in all $\mathcal{R} = 200$ realisations, $P(N_R(t) = 0) = 1$ (standard error of the mean in $P(N_R(t = 500) = 0)$ is below 4%; see S1 Appendix Sec. 3.3.2). The white dashed line represents an axis break separating $m = 0$ and $m = 10^{-5}$ on a logarithmic scale. The green line shows the theoretical prediction of Eq. (6) for the critical migration rate. The grey line indicates the bottleneck strength used within panel (B). Annotation letters at specific $(m, K_+/K_-)$ values refer to the corresponding panels (C-K). **(B)** Same panel as Fig 4E showing how $P(N_R(t) = 0)$ vs migration rate $m$ changes in time (for $t = 100, 200, 300, 400, 500$, from bottom to top) at fixed bottleneck strength, $K_+/K_- = 70.7$, illustrating that slow migration enhances the eradication of $R$; see Eq. (9) in Results. The lines/symbols denote the mean across $\mathcal{R} = 200$ realisations and areas/error bars indicate confidence intervals computed via a Wald interval (Sec. 3.3.2). **(C-K)** Temporal evolution of the fraction $\rho_S(t)$ of $R$-free demes across the metapopulation for one example realisation in each panel (see Eq.(7)). As indicated in (A), migration rates are $m \in \{0, 10^{-3}, 10^{-2}\}$ (from left to right panels) and bottleneck strengths are $K_+/K_- \in \{200, 50, 12.5\}$ (top to bottom). Green bands indicate the times in which each deme experiences a harsh environment, ($K_- = 80$; see Sec. 3.3.1). The almost unnoticeable blue line in (E), indicates a very small fraction $\rho_R(t)$ of $R$-only ($S$-free) demes under vanishing migration ($m \to 0$); see Sec. 5.1 and Fig. S5A-D. S4 Movie shows the full spatial metapopulation dynamics for the parameters of panel (I).

that the fluctuation-driven eradication of resistance is robust against the specific form of migration, and that its characterisation by (8) and (9) applies regardless of the specific choice made for the dispersal of the metapopulation.

#### 5.4 Number of $R$ and $S$ cells in coexisting demes

In this work, we have studied in detail how the interplay of environmental and demographic fluctuations can eradicate resistance from a grid metapopulation (we have also considered a cycle and $d$-dimensional lattices in Sec. 5.5). For the sake of completeness, here we briefly discuss the dynamics in demes when resistant and sensitive cells *coexist* ("coexisting demes"). The coexistence of strains can be for a short transient preceding the fluctuation-driven eradication of $R$ (when conditions (8) are satisfied), or for extended periods of time (e.g. under fast switching; see Fig S3C).

To characterise how the composition of coexisting demes changes in time, we compute the average number $N_{R_c/S_c}(t)$ of $R$ and $S$ cells in demes where they coexist at time $t$ across the grid:

$$N_{R_c/S_c}(t) = \frac{1}{L^2} \sum_{\vec{u}} N_{R/S}(\vec{u}, t) \cdot \mathbb{1}_{\{N_{R/S}(\vec{u},t)>0\}} \cdot \mathbb{1}_{\{N_{S/R}(\vec{u},t)>0\}}, \qquad \text{(S5.12)}$$

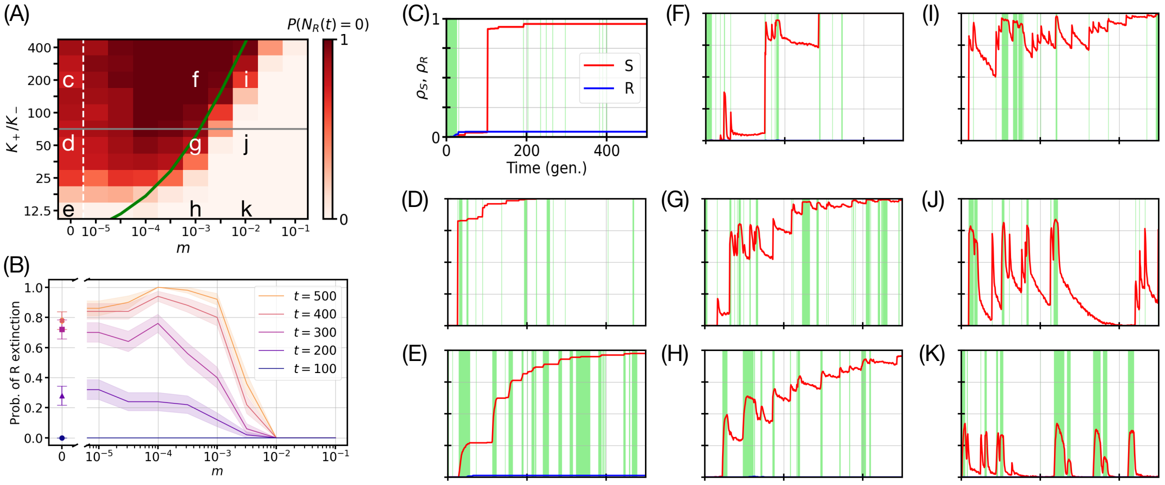


Figure S7: **Fluctuation-driven eradication of resistance with density-independent migration.** Same environmental parameters $(\nu, \delta) = (0.1, 0.5)$ (see Methods and Table 1 for all parameters), and formatting as in Fig S6, but with density-independent migration implemented according to Eq. (2b). **(A)** Same heatmap as in Fig 4D and S6A showing the probability $P(N_R(t) = 0)$ of total extinction of $R$ (resistant) microbes as a function of bottleneck strength, $K_+/K_-$, and migration rate $m$ after 500 microbial generations ($t = 500$). The colour bar in panel (A) varies from light to dark red, with the darkest red corresponding to eradication of $R$ in all $\mathcal{R} = 50$ realisations, $P(N_R(t) = 0) = 1$ (standard error of the mean in $P(N_R(t = 500) = 0)$ is below 7%; see S1 Appendix Sec. 3.3.2). As in Fig S6A, the white dashed line represents an axis break separating $m = 0$ and $m = 10^{-5}$ on a logarithmic scale. The green line shows the prediction of Eq. (6) for the critical migration rate. The grey line indicates the bottleneck strength used in (B). Annotation letters at specific $(m, K_+/K_-)$ values refer to the corresponding panels (C-K). **(B)** Probability of $R$ extinction $P(N_R(t) = 0)$ as a function of migration rate $m$ for a bottleneck strength of $K_+/K_- = 70.7$ at five different times, for $t = 100, 200, 300, 400, 500$ (from bottom to top). The lines/symbols denote the mean across $\mathcal{R} = 50$ realisations and areas/error bars indicate confidence intervals computed via a Wald interval (Sec. 3.3.2). **(C-K)** Fraction of demes without $R$ cells ($\rho_S$, red) and without $S$ cells ($\rho_R$, blue) as a function of time for different values of the density-independent migration rate and bottleneck strength: (left to right) $m \in \{0, 10^{-3}, 10^{-2}\}$ and (top to bottom) $K_+/K_- \in \{200, 50, 12.5\}$. In (C) and (E), we notice a small fraction $\rho_R$ of $R$-only ($S$-free) demes under vanishing migration ($m \to 0$); see Sec. 5.1 and Fig. S5A-D. Green bands indicate the times in which each deme experiences a harsh environment, ($K_- = 80$, see Sec. 3.3.1). S5 Movie shows the full spatial metapopulation dynamics for the parameters of panel (I).

where $\mathbb{1}_{\{N_{S/R}(\vec{u},t)>0\}}$ is the indicator function, equal to 1 if $N_{S/R}(\vec{u}, t) > 0$ and 0 otherwise (Results). In Fig S8, $N_{R_c/S_c}(t)$ correspond to the number of $R$ and $S$ cells averaged over the demes of the grid in which they coexist at time $t$ in a *single realization* of the metapopulation. These quantities have been computed for the parameter set of Fig 3 and are reported in Fig S8.

These results show that demes where $R$ and $S$ microbes coexist (indicated by pink pixels in Fig 3A-F) consist approximately of $N_{\text{th}}$ and $K(t) - N_{\text{th}}$ cells of type $R$ and $S$ in the regime of intermediate switching ($\nu \sim s \lesssim 1, 0 \leq \delta \lesssim 1$). More specifically, in the examples considered here where environmental variations are characterised by strong bottlenecks (when $N_{\text{th}} < K_- \ll K_+$), the coexisting demes consist of an overwhelming majority of $S$ cells in the mild environmental state (where $K = K_+$). Bottlenecks thus cause sharp decays in the number of $R$ and $S$ in each deme, following which demographic fluctuations can drive $R$ cells to local extinction when the conditions (8) are satisfied (Results, Discussion); see S2-S4 Movies.

#### 5.5 Slow migration can speed up and enhance the eradication of R cells in a one-dimensional metapopulation

As explained in Results and Discussion (see also Sec. 3.1), this work focuses on a two-dimensional lattice (grid) metapopulation to investigate the effects of cell migration on the eradication of resistant cells, but most of the results can be generalised to metapopulation lattices (regular graphs) of any

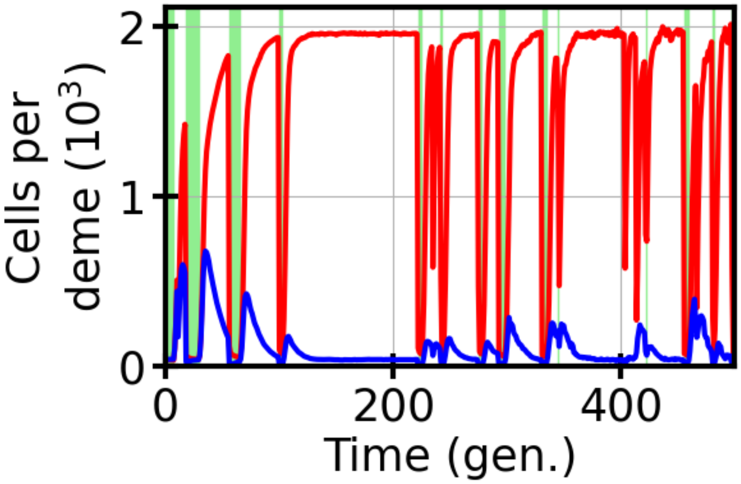


Figure S8: **Number of $R$ and $S$ cells in coexisting demes.** Number $N_{S_c}(t)$ of $S$ (red) cells and number $N_{R_c}(t)$ of $R$ (blue) microbes in coexisting demes averaged across the metapopulation grid as a function of time $t$; see Eq. (S5.12). $N_{S_c}$ and $N_{R_c}(t)$ thus represent the average number of $S$ and $R$ cells through the pink pixels of Fig 3A-F at a given time (microbial generation). The parameters are the same as in Fig 3: $L = 20, K_+ = 2000, K_- = 80, s = 0.1, a = 0.25, \nu = 0.1, \delta = 0.5$ and $m = 0.001$ (with density-dependent migration according to Eq. (2a)), see also Table 1. Green bands indicate periods of a harsh environmental state (Sec. 3.3.1).

spatial dimension $d$ (Results, Discussion). In this section, for the sake of concreteness we extend our discussion to the case of the fluctuation-driven eradication of resistance from a periodic one-dimensional lattice, and then briefly consider the general case of a $d$-dimensional lattice.

Before diving into the one-dimensional case, it is worth noting that in a metapopulation arranged on a $d$-dimensional lattice of linear size $L$, the total number of sensitive and resistant cells scales as $\mathcal{O}(K(t)L^d)$. Here we consider $L^d = 100 - 1000$. As explained in Sec. 3, our simulations correspond to a good compromise between computational efficiency and experimental relevance and allow us to characterise the phenomenon of fluctuation-driven eradication in lattice metapopulations. While the two-dimensional case ($d = 2$, with $L = 20$) is extensively covered in the main text, we present here results obtained for the one-dimensional (1D) case ($d = 1$) with $L = 100$, and demonstrate the robustness of our main findings.

Similarly as in Model & Methods, we specifically consider microbial metapopulations that can be seen as a periodic 1D lattice (a cycle or ring) of size $L$, containing $L$ demes denoted by $u \in \{1, 2, \ldots, L\}$, with $L = 100$ in our example. Again, the demes represent (well-mixed) subpopulations that are connected to their two nearest neighbours via migration, at a per capita rate proportional to $m$, see Eqs. (2a)-(2b). Each deme of the cycle has the same carrying capacity, denoted by $K$, and at time $t$ consists of $N_{S/R}(u)$ cells of type $S/R$.

The simulations of the eco-evolutionary dynamics on the cycle metapopulation follow the prescription described in Section 3.1. As in the case of the grid, we have investigated the fluctuation-driven eradication of resistance from the cycle and tested the conditions (8) and (9) governing this phenomenon (Results, Discussion). To this end, we have computed the probability $P(N_R(t) = 0)$ of $R$ eradication for the same parameters used in Fig 4 for the grid. Our results are presented in Fig S9, that should be compared with Fig 4 of which it is the one-dimensional counterpart. This comparison shows that Fig S9 has similar qualitative features as those obtained in two dimensions, with the fluctuation-driven eradication of resistance enhanced by slow migration when $\nu/K_+ < m \lesssim m_c$. These results therefore qualitatively confirm the predictions of (8) and (9); see Results and Discussion. However, we note in Fig S9 that the eradication of resistant cells ($P(N_R(t) = 0) = 1$) occurs for a broader range of values ($m$, $K_+/K_-$) than in Fig 4. In particular, we notice that in Fig S9 resistance eradication can occur for values of $m$ exceeding significantly the prediction (6) of the critical value $m_c$ (red/dark areas in Fig S9 outgrow the green curves). This stems from (6) being a mean-field prediction that ignores spatial correlations that are more important in 1D than in 2D (Results and Discussion).

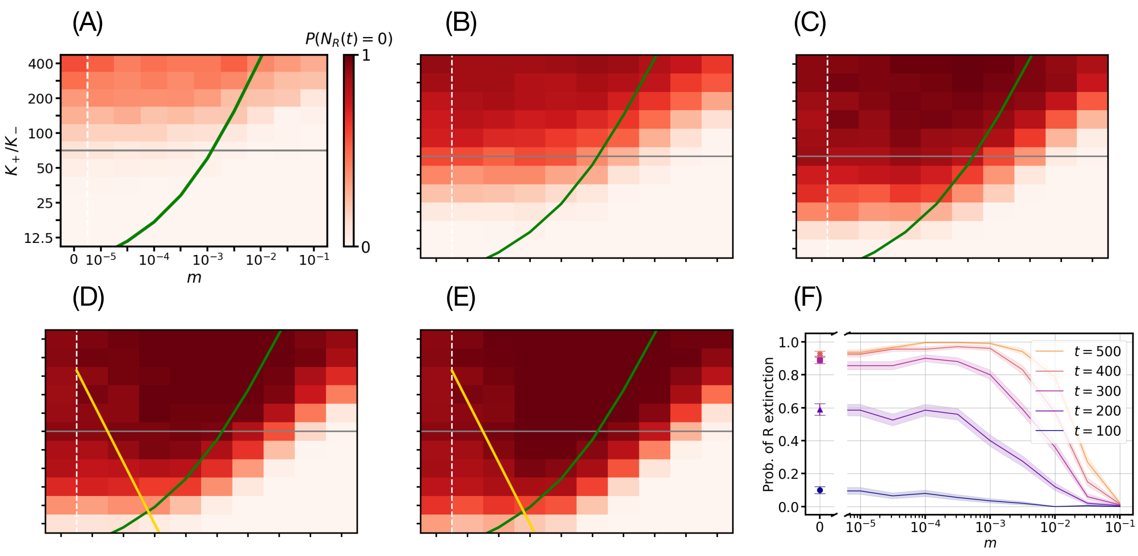


Figure S9: **One-dimensional metapopulation lattice: Slow migration can speed up and enhance the eradication of $R$ cells on a cycle.** Time evolution of the heatmap showing the probability of $R$ extinction $P(N_R(t) = 0)$ on a cycle metapopulation of length $L = 100$ as a function of bottleneck strength, $K_+/K_-$, and migration rate $m$ (implemented according to Eq. (2a) at time (microbial generation) **(A)** $t = 100$, **(B)** $t = 200$, **(C)** $t = 300$, **(D)** $t = 400$, and **(E)** $t = 500$ with environmental parameters $(\nu, \delta) = (0.1, 0.5)$; other parameters are as in Table 1 (Discussion). Each $(m, K_+/K_-)$ value pair is averaged over an ensemble of $\mathcal{R} = 200$ independent metapopulation simulations. The colour bar showing the value of $P(N_R(t) = 0)$ ranges from light to dark red indicating the fraction of the simulations for which $R$ has been eradicated from the cycle by time $t$ (standard error of mean in $P(N_R(t = 500) = 0)$ is below 4%; see Sec. 3.3.2). The green and dashed white lines represent the theoretical prediction of Eq. (6) and an eye-guiding axis break, respectively. The golden lines in panels (d-e) show $K_+/K_- = \frac{\nu}{mK_-}$, with $P(N_R(t) = 0) \approx 1$ in the (upper) region between the golden and green lines, according to Eq. (9). The grey horizontal lines in panels (A-E) indicate the example bottleneck strength of panel (F). **(F)** Probability of $R$ extinction $P(N_R(t) = 0)$ as a function of migration rate $m$ at bottleneck strength $K_+/K_- = 70.7$ for $t = 100, 200, 300, 400, 500$ (bottom to top). Solid lines (full symbols at $m = 0$) indicate average across $\mathcal{R} = 200$ realisations; shaded areas (error bars at $m = 0$) indicate binomial confidence interval computed via the Wald interval (Sec. 3.3.2).

The comparison of the results obtained for the grid and cycle metapopulations, therefore leads us to draw the following conclusions:

- The fluctuation-driven eradication of resistance holds on a grid and a cycle, and its general features are well characterised by the conditions (8) and (9). This phenomenon being triggered by strong bottlenecks (and enhanced by slow migration), regardless of the spatial dimension, is robust and expected to hold on lattice of any dimension $d$ (Results, Discussion).

- The conditions (8) and (9) characterising the mechanism of fluctuation-driven eradication of $R$ depend on the spatial dimension $d$ only through the the critical migration rate $m_c$, of which Eq.(6) is a mean-field approximation. As such, the expression (6) ignores spatial correlations, and is therefore a better approximation for 2D than 1D metapopulation lattices. Therefore, while (8) and (9), with (6), allow us to quantitatively describe the fluctuation-driven eradication of $R$ on the grid, they provide a useful qualitative description of this phenomenon on the cycle. Accordingly, we expect (8) and (9), with (6), to accurately characterise the fluctuation-driven eradication of resistance on three-dimensional lattices and, more generally, on lattices of dimension $d \geq 2$ (Results, Discussion).

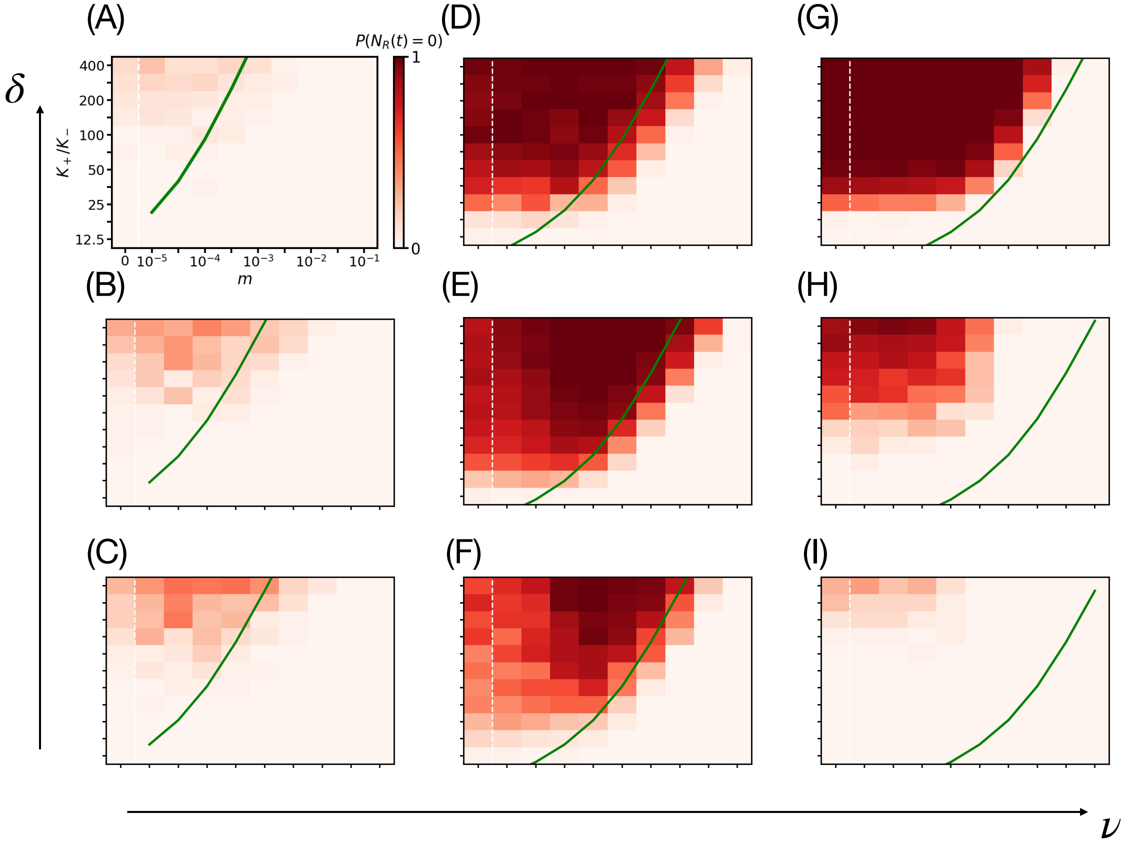


Figure S10: **Further exploration of fluctuation-driven eradication of resistance in the $(\nu, \delta)$ parameter space. (A-I)** Additional heatmaps showing for a set of environmental parameters $(\nu, \delta)$ the probability $P(N_R(t = 500) = 0)$ of $R$ eradication from a grid of size $L = 20$ as a function of the migration rate $m$ (implemented according to Eq. (2a)) and bottleneck strength $K_+/K_-$ after 500 microbial generations. Here, $s = 0.1, a = 0.25, N_{\text{th}} = 40$; see Table 1 (Discussion) for the other parameters. The colour bar varies from light to dark red, where the darkest red corresponds to the eradication of $R$ in all realisations. The white dashed lines and solid green lines respectively indicate axes breaks on the logarithmic scale and theoretical prediction of Eq. (6) for the critical migration rate (Results). Panels (E) and (G) correspond to Figs 4D and 2A, with $\mathcal{R} = 200$ realisations per pixel (standard error of the mean in $P(N_R(t = 500) = 0)$ is below 4%; see Sec. 3.3.2), whereas each pixel in the other panels (A)-(D), (F), and (H)-(I) results from $\mathcal{R} = 50$ independent simulations (standard error of the mean below 7%). The environmental switching rate varies from left to right panels, $\nu \in \{0.01, 0.1, 1\}$, and the environmental bias varies from top to bottom, $\delta \in \{0.75, 0.5, 0.25\}$.

#### 5.6 Fluctuation-driven eradication of resistance for further sets of environmental parameters $(\nu, \delta)$

In Figs 2-4 and Figs S5, S6, S7, and S8 we have investigated the fluctuation-driven eradication of resistance for a fixed small number of environmental parameters $\nu, \delta$.

Here, we corroborate the characterisation of this phenomenon by considering further sets of environmental parameters, namely for $\nu \in \{0.01, 0.1, 1\}$ and $\delta \in \{0.75, 0.5, 0.25\}$ in Fig S10. These results confirm that the fluctuation-driven eradication of resistance occur in the intermediate switching regime ($\nu \sim s \lesssim 1, 0 \lesssim \delta \lesssim 1$), as it can hardly be observed in Fig S10A,I (we checked that this is not observed for $\nu = 10$ either, e.g., see Figs S1C and S3C).
